# Neonatal NMDA receptor blockade disrupts prefrontal cortical regulation of social affective behavior: implications for schizophrenia

**DOI:** 10.64898/2026.11.04.764439

**Authors:** Reina Tachihara, Natsumi Hayashi, Carlos González del Rosal, Tingbi Xiong, Keita Igarashi, Takaaki Ozawa, Miyo Hori, Kazuo Yamada, Michimasa Toyoshima

## Abstract

Developmental N-methyl-D-aspartate receptor (NMDAr) hypofunction has been implicated in the pathophysiology of schizophrenia, including cognitive and emotional deficits. However, the neural circuit mechanisms linking developmental NMDAr hypofunction to social affective impairments remain poorly understood. Here, we investigated schizophrenia-related social affective dysfunction induced by neonatal exposure to the NMDAr antagonist MK-801 and examined the contribution of the medial prefrontal cortex (mPFC), a key hub for socio-cognitive function. Sprague-Dawley rat pups received subcutaneous injections of saline or MK-801 (MK-Low: 0.2 mg/kg; MK-High: 0.4 mg/kg) during the neonatal period. In adulthood, social affective behaviors were assessed using the Social Affective Preference (SAP) and Social Interaction (SI) tests. MK-801-treated rats, particularly the MK-High group, displayed impaired social affective responses, characterized by reduced preference for and investigation of distressed conspecifics. These impairments were ameliorated by chemogenetic inhibition of pyramidal cells in the mPFC, whereas activation of these neurons disrupted social affective behavior in naïve rats. Histological analyses revealed reduced parvalbumin-positive interneuron (PVIN) density in the mPFC of MK-801-treated rats. Consistent with the involvement of PVIN dysfunction, selective activation of mPFC PVINs alleviated social affective deficits in MK-801-treated rats. These findings identify impaired PVIN-mediated inhibitory regulation within the mPFC as a circuit mechanism underlying social affective dysfunction following neonatal NMDAr blockade, providing insight into the pathophysiology of schizophrenia-related social impairments.

## Introduction

Schizophrenia is a severe and heterogeneous psychiatric disorder that affects approximately 23 million people worldwide. Patients exhibit a wide range of symptoms, including delusions, hallucinations, cognitive dysfunction, and negative symptoms [1]. These symptoms, alone or in combination, significantly impair daily functioning. Among the consequent functional impairments, social deficits represent a major problem and are closely linked to cognitive dysfunction and negative symptoms [2]. Patients with schizophrenia often experience difficulties in processing social information, such as recognizing and responding to others’ emotions [3], as well as in higher-order social cognition, including mentalizing and Theory of Mind (ToM) [4]. In particular, patients show aberrant emotional responses to others’ affective states, which constitute a core component of social functioning. Recent studies have reported that impaired social cognitive abilities and empathic deficits may be attributed, at least in part, to disturbances in the activity of the medial prefrontal cortex (mPFC) [3], a region that plays a crucial role in social functioning in mammals [5]. The importance of the mPFC is also supported by studies using animal models of schizophrenia, in which mPFC dysfunction is associated with social impairments [6–8]; however, direct causal evidence linking specific mPFC microcircuit abnormalities to defined domains of social dysfunction, particularly social affective responses, remains limited.

One of the leading pathophysiological accounts of schizophrenia is the “glutamate hypothesis,” which posits that the disorder results from hypofunction of N-methyl-D-aspartate-type glutamate receptors (NMDAr) [9]. While the “dopamine hypothesis” has been primarily invoked to explain the emergence of positive symptoms such as delusions and hallucinations [10], the glutamate hypothesis is considered to more directly account for the negative and cognitive symptoms of schizophrenia. Furthermore, several schizophrenia risk genes [11], such as DISC1 [12] and NRG1 [13], are implicated in the regulation of NMDAr function or downstream signaling pathways, further supporting the glutamate hypothesis.

Studies using laboratory animals also support a link between NMDAr hypofunction and schizophrenia-related phenotypes, particularly cognitive, attentional, and social deficits. Rodents treated with NMDAr antagonists such as PCP or MK-801 exhibit schizophrenia-like behavioral abnormalities, including impairments in set shifting, working memory, and prepulse inhibition [14–16]. Moreover, genetic deletion of NMDAr subunits [17] or disruption of NMDAr–dependent signaling pathways [18,19], including those involving DISC1 and NRG1, also produces working memory and attentional deficits. Together, these findings suggest that loss or hypofunction of NMDArs contributes to a broad spectrum of schizophrenia-related behavioral abnormalities, ranging from positive-like symptoms to cognitive impairments.

The NMDAr hypofunction theory also provides a framework for explaining the neurobiological and anatomical abnormalities observed in patients. In schizophrenia, disturbances of inhibitory circuitry in the neocortex are frequently reported, particularly involving parvalbumin-expressing interneurons (PVINs) [20,21]. Levels of GABA-synthesizing enzymes such as GAD67 and parvalbumin protein, as well as the mRNAs encoding these proteins, are decreased in postmortem brains of patients with schizophrenia [22]. Specifically, a reduction in GAD67 expression in PVINs is sometimes observed [23], suggesting that PVINs are preferentially affected and represent a critical cellular target for schizophrenic phenotypes. PVINs highly express NMDArs during early postnatal development [24]. In the absence of these receptors, PVINs fail to acquire normal functional properties and connectivity, particularly at excitatory–inhibitory synapses. Because PVIN activity is partially driven by excitatory input from pyramidal neurons, loss of these synapses leads to reduced PVIN activity [25]. PVIN-mediated fast-spiking inhibition is essential for cortical oscillations, especially for the generation of gamma-band activity [26], which is markedly impaired in schizophrenia [27]. Notably, abnormalities in gamma oscillations are prominent during behavioral tasks, and correlate with working memory [28] and social cognitive [29] deficits in patients, suggesting that these neurobiological changes are closely linked to the symptomatology of schizophrenia, including social dysfunction.

Recently, animal studies have begun to elucidate a causal relationship between NMDAr hypofunction, PVIN dysfunction, and schizophrenia-related behavioral phenotypes. PVIN abnormalities are frequently observed in animals treated with NMDAr antagonists such as MK-801 or ketamine during the neonatal period [30,31]. In addition to reproducing schizophrenia-like cognitive and attentional deficits, mice with NMDAr deletion during early postnatal development also exhibit behavioral abnormalities and disruption of PVIN functions in adulthood [32], underscoring the importance of NMDAr function in PVINs during a critical developmental window. Moreover, acute MK-801–induced cognitive deficits can be ameliorated by optogenetic stimulation of PVINs at gamma frequencies [33], highlighting PVINs as a promising therapeutic target. However, most of these studies have focused on working memory, attention, or general social interaction, and it remains largely unknown how neonatal NMDAr hypofunction in PVINs contributes to specific domains of social dysfunction, such as impaired social affective responses.

Here, we hypothesized that hypofunction of NMDArs during early development induces schizophrenia-like social affective deficits, and that these deficits are mediated by dysfunction of inhibitory circuits in the mPFC, particularly those involving PVINs. To test this hypothesis, we chronically treated rat pups with MK-801 and subsequently assessed their social behavior in adulthood. We employed the Social Affective Preference (SAP) test [34] and a one-on-one social interaction test to examine whether their social affective responses toward distressed conspecifics were impaired, as observed in patients with schizophrenia. These assessments revealed that the social affective response toward a distressed conspecific was selectively impaired in MK-801–treated rats, especially at higher doses, in both the SAP and one-on-one interaction paradigms. Consistent with previous reports, MK-801–treated rats also exhibited compromised inhibitory circuitry in the mPFC, with particularly pronounced alterations in PVINs. Finally, we demonstrated that enhancing mPFC inhibition, including PVIN-targeted manipulations, restored social interest toward a distressed conspecific in MK-801–treated rats. Together, these findings provide causal evidence linking PVIN hypofunction in the mPFC to social affective deficits in this model and highlight PVINs as a potential therapeutic target for the social and cognitive domains of schizophrenia.

## Methods

### Animals

Sprague–Dawley rats were obtained from Jackson Laboratory Japan (Kanagawa, Japan) and Japan SLC Inc. (Shizuoka, Japan). Rat pups born from timed-pregnant dams were used as subjects. Pregnant rats were individually housed for parturition. Male and female demonstrator rats for behavioral tests were brought into the animal facility at 6 weeks of age and housed in groups of 2–3 per cage. All rats were maintained under a 12:12 h light/dark cycle with food and water available ad libitum. All experimental procedures were approved by the University of Tsukuba Committee on Animal Research and were conducted in accordance with institutional guidelines.

### Chronic MK-801 injections during the neonatal period

(+)-MK-801 hydrogen maleate (MK-801; Cat#: M107, Sigma-Aldrich) dissolved in physiological saline (Otsuka Pharmaceutical Co., Ltd.) was used for neonatal treatment. The day of birth was defined as postnatal day (PND) 0. On PND 6, pups were divided into three treatment groups. From PND 7 to 20, they received subcutaneous injections of physiological saline (vehicle), MK-801 at 0.2 mg/kg, or MK-801 at 0.4 mg/kg, twice daily, with injections separated by at least 8 h. Animals were weaned on PND 25-28 and subsequently housed in groups of 2–3 per cage with food and water ad libitum.

### Stereotaxic surgery and chemogenetic manipulation

For chemogenetic manipulation, we used the designer receptors exclusively activated by designer drugs (DREADDs) system.

#### Viral vectors

AAV5-CaMKIIa-hM4Di-mCherry (Cat#: 50477-AAV5, Lot: v167951, 1.6 × 10^13 GC/mL; Lot: v206877, 7.1 × 10^12 GC/mL), AAV5-CaMKlla-hM3Gq-mCherry (Cat#: 50476, Lot: v176266, 1.1 × 10^13 CG/mL), AAV5-CaMKlla-mCherry (Cat#: 114469-AAV5, Lot: v152046, 2.3 × 10^13 CG/mL), and AAV9-S5E2-Gq-P2A-dTomato (Cat#: 135635-AAV9, Lot: v214407, 1.9 × 10^13 GC/mL) were purchased from Addgene.

#### Surgical procedure

At 8 weeks of age, subjects were anesthetized with an intraperitoneal injection of ketamine (60 mg/kg; Fujita Pharmaceutical Co., Ltd.) and xylazine (6 mg/kg; Fujita Pharmaceutical Co., Ltd.). Rats were then mounted on a stereotaxic frame (Kopf Instruments) and received bilateral infusions of viral vectors into the mPFC (coordinates: AP +3.1 mm, ML ±0.6 mm, DV −3.8 mm from bregma; 1 µL per side at 0.2 µL/min) using a 10 µL Hamilton syringe (Cat#: 4015-11010, Hamilton) filled with white mineral oil (Cat#: M5904, Merck). After each infusion, the syringe was left in place for an additional 10 min to allow diffusion of the virus. Immediately after surgery, rats received a subcutaneous injection of meloxicam (Boehringer Ingelheim Animal Health Japan Co., Ltd.) for postoperative analgesia. Rats were allowed to recover for at least 2 weeks before behavioral testing.

#### Chemogenetic manipulation

Compound 21 (C21; Cat#: HB6124, Hello Bio) was used as a ligand for DREADD activation. In the DREADD-activated condition, subjects received an intraperitoneal injection of C21 dissolved in saline (0.5 mg/kg) 90 min before behavioral testing. In the control condition, subjects received an intraperitoneal injection of saline (1 mL/kg) instead of C21.

### Behavioral experiments

Behavioral tests were performed in adulthood during the light phase of the 12:12 h light/dark cycle.

#### Social Affective Preference (SAP) test

The SAP test was conducted in a three-chambered acrylic box (40 cm × 80 cm × 50 cm; Medical Workshop, Research Facility Center for Science & Technology, University of Tsukuba, Ibaraki, Japan) equipped with two wire cylinders. Prior to testing, subject rats were habituated to the apparatus for 20 min on each of 2 consecutive days. On the first day (arena habituation), they were placed in the arena with two empty cylinders. On the second day (social habituation), they were exposed to two cylinders containing age- and sex-matched unfamiliar conspecifics. On the test day, subjects were exposed to two cylinders containing conspecifics in different affective states: one neutral (intraperitoneal saline injection) and one distressed (intraperitoneal injection of lithium chloride (LiCl) solution at 150 mg/kg or 65 mg/kg). Subjects were allowed to freely explore the apparatus for 10 min. Sniffing behavior directed toward each cylinder and self-grooming behavior during the test were recorded. A preference index was calculated using the following formula:

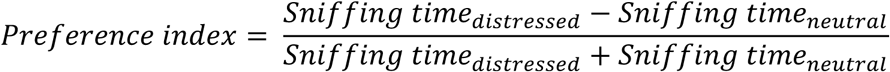

#### Social interaction (SI) test

The SI test was conducted in an acrylic box (30 cm × 40 cm × 40 cm). On Day 1, subjects underwent arena habituation for 15 min. On Day 2 (social habituation), they were allowed to engage in one-on-one interaction with an age- and sex-matched unfamiliar conspecific in the box for 15 min. On one test day, subjects encountered a conspecific (demonstrator) in a distressed state induced by an intraperitoneal injection of LiCl (150 mg/kg). On the other test day, subjects encountered a conspecific in a neutral state (saline-injected). The order of experimental conditions was counterbalanced across subjects. Social interactions between subjects and demonstrator rats were recorded for 15 min. The time spent in allo-grooming (grooming behavior directed toward the partner), sniffing, self-grooming (grooming directed toward the self), and digging was measured.

### Histology

#### Tissue preparation

For immunostaining and verification of viral expression, fixed brain samples were collected from subject rats. Rats were deeply anesthetized with isoflurane (Cat#: 095-06573, FUJIFILM Wako Pure Chemical Corporation) and transcardially perfused with 0.02 M phosphate-buffered saline (PBS), followed by 4% paraformaldehyde (Cat#: 02890-45, Nacalai Tesque, Inc.) in 0.1 M phosphate buffer (PB). Brains were post-fixed in 4% paraformaldehyde for 24 h at 4°C and then cryoprotected in 30% sucrose (Cat#: 190-00013, FUJIFILM Wako Pure Chemical Corporation) in 0.1 M PB. At least 4 days later, coronal sections (40 µm thick) were cut using a microtome (Retratome, Yamato Koki Industry Co., Ltd.).

#### Verification of viral expression

Sections were washed with 0.02 M PBS (3 × 10 min), mounted on microscope slides, and coverslipped with ProLong Glass Antifade Mountant with NucBlue (Cat#: P36981, Invitrogen). Fluorescent images were acquired using a fluorescence microscope (BZ-X710, Keyence) with a 4× objective to determine the localization of viral expression. Subjects with absent or weak viral expression in either hemisphere of the mPFC (Brodmann area 32; Paxinos and Watson, 2015) were excluded from the analysis.

#### Fluorescent immunohistochemistry

To visualize GABA-positive and parvalbumin-positive neurons in the mPFC, fluorescent immunohistochemistry was performed. Sections were washed with 0.02 M PBS (3 × 10 min) and then incubated in a blocking solution for 30 min at room temperature (for GABA staining: 1% bovine serum albumin (BSA; Cat#: G9023, Sigma-Aldrich Japan) in 0.02 M PBS; for parvalbumin staining: 1% normal goat serum (NGS; Sigma-Aldrich Japan) in PBS-T (0.1% Triton X-100 in 0.02 M PBS)). Sections were then incubated overnight at 4°C with a rabbit anti-GABA antibody (1:5000, Cat#: P2052, Abcam) or a mouse anti-parvalbumin antibody (1:4000, Cat#: P3088, Abcam). After washing with 0.02 M PBS (3 × 10 min), sections were incubated with a goat anti-rabbit IgG Alexa Fluor 488 antibody (1:300, Cat#: ab150077, Abcam) or a goat anti-mouse IgG Alexa Fluor 488 antibody (1:300, Cat#: ab150080, Abcam) for 70 min at room temperature. Following additional washes (3 × 10 min), sections were mounted on microscope slides and coverslipped with ProLong Glass Antifade Mountant with NucBlue.

Fluorescent images were captured using a 20× objective on a fluorescence microscope (BZ-X710, Keyence). The number of GABA-positive and PV-positive neurons in the mPFC was manually quantified using the cell counter plugin in ImageJ2 (version 2.14.0, National Institutes of Health). For each subject, expression levels were determined by averaging counts from 3-4 images across 2 sections.

### Statistics

All statistical analyses were performed using R (version 4.5.2) in RStudio (version 2025.09.2+418; RStudio, PBC).

For behavioral analysis in the SAP test, sniffing time was analyzed using a two-way ANOVA with group and affective state of the demonstrators (distressed vs. neutral) as factors, followed by Bonferroni post hoc tests. One-sample t-tests were used to compare the preference index of each group against chance level (0), which corresponds to equal exploration of neutral and distressed demonstrators. In addition, following a one-way ANOVA, pairwise t-tests were performed to examine between-group differences in the SAP test. Correlation analyses between time spent in sniffing and self-grooming during the SAP test were conducted using Pearson’s correlation.

A two-way ANOVA was used to analyze the time spent on each behavior during the SI test, followed by multiple comparisons with the demonstrators’ group and state (distressed or neutral) as factors.

The proportion of behavioral categories during the SI test was visualized as pie charts and analyzed using generalized linear models with the subjects’ group and the demonstrator (the experiment shown in Figure 1) or experimental condition (the experiments shown in Figures 2,3, and 5) as factors, followed by Type III Wald chi-square tests and post hoc comparisons using estimated marginal means.

**Figure 1.**
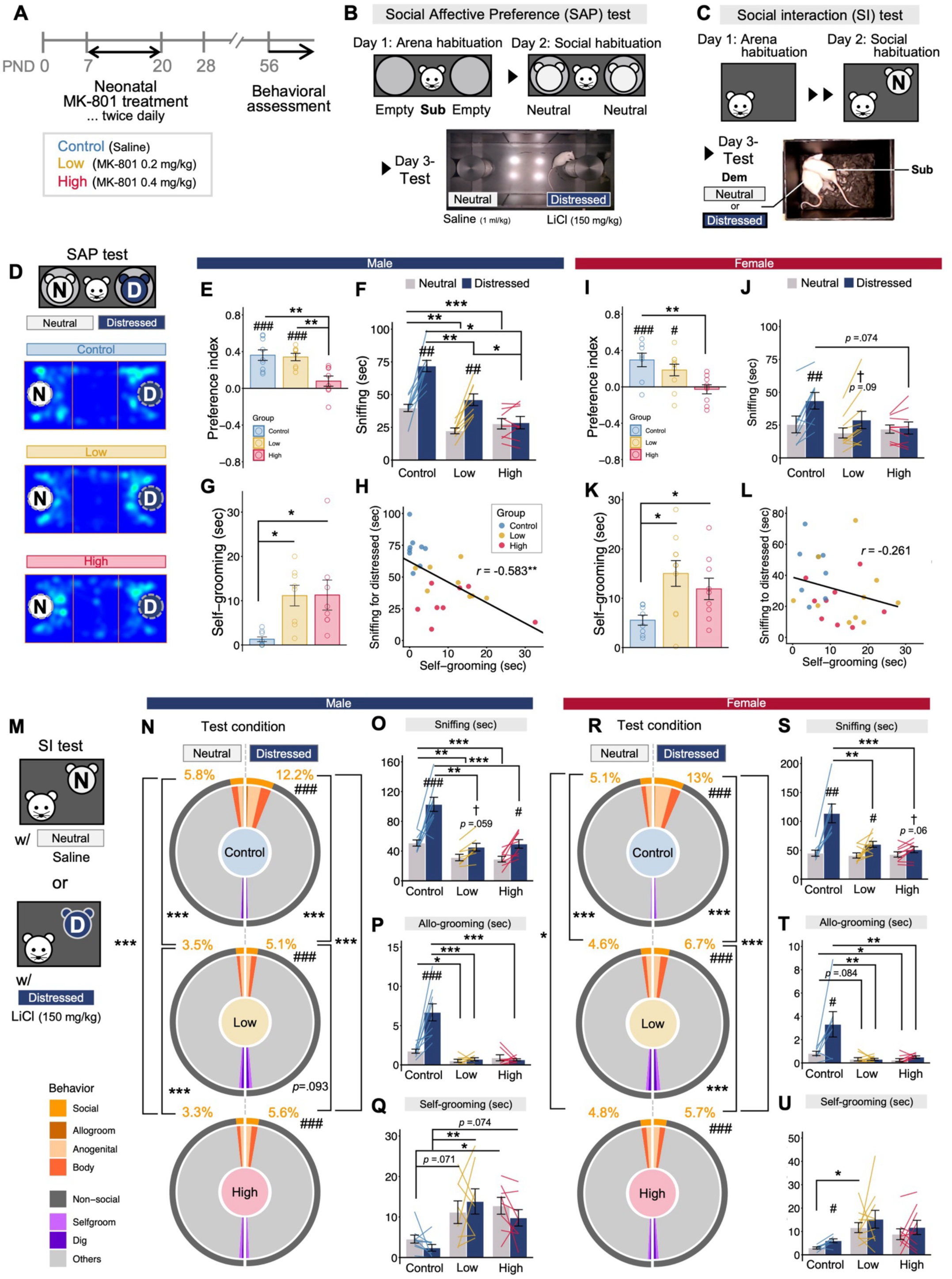
Social affective responses were impaired in rats chronically treated with MK-801 during the neonatal period. **A** Diagram of the experimental procedure. Subjects received daily injections of MK-801 (Low: 0.2 mg/kg; High: 0.4 mg/kg) or saline from postnatal day (PND) 7 to 20. Behavioral assessments were conducted in adulthood after PND 56. **B** Schematic illustration of the social affective preference (SAP) test paradigm. **C** Schematic illustration of the social interaction (SI) test paradigm. **D** Representative heatmap showing the time spent in each area during the SAP test. **E–L** Data were obtained from a total of 51 subjects (Control: male n = 9, female n = 8; Low: male n = 8, female n = 10; High: male n = 8, female n = 9). **E–H** Behavioral measurements during the SAP test in males: preference index (**E**), time spent sniffing each demonstrator (**F**), time spent self-grooming during the test (**G**), and correlation between time spent sniffing the distressed demonstrator and self-grooming behavior (**H**). **I–L** Behavioral measurements during the SAP test in females: preference index (**I**), time spent sniffing each demonstrator (**J**), time spent self-grooming during the test (**K**), and correlation between time spent sniffing the distressed demonstrator and self-grooming behavior (**L**). **M** Experimental conditions in the SI test. Each subject was tested under two conditions across two test days: SI with a neutral partner and SI with a distressed partner. **N–U** Data were obtained from a total of 50 subjects (Control: male n = 9, female n = 7; Low: male n = 8, female n = 10; High: male n = 8, female n = 9). **N–Q** Behavioral measurements during the SI test in males: pie chart showing the proportion of behavioral categories during the SI test (**N**), time spent sniffing (**O**), allo-grooming (**P**), and self-grooming (**Q**). **R–U** Behavioral measurements during the SI test in females: pie chart showing the proportion of behavioral categories during the SI test (**R**), time spent sniffing (**S**), allo-grooming (**T**), and self-grooming (**U**). Bar and line graphs show the mean ± s.e.m. \**p* < 0.05, \*\**p* < 0.01, and \*\*\**p* < 0.001 indicate significant differences between groups. #*p* < 0.05, ##*p* < 0.01, ###*p* < 0.001, and †*p* < 0.10 indicate significant differences or trends between neutral and distressed demonstrators in panels **F**, **J**, and **N–U**, or significant differences or trends from chance level in the preference index in panels **E** and **I**.

**Figure 2.**
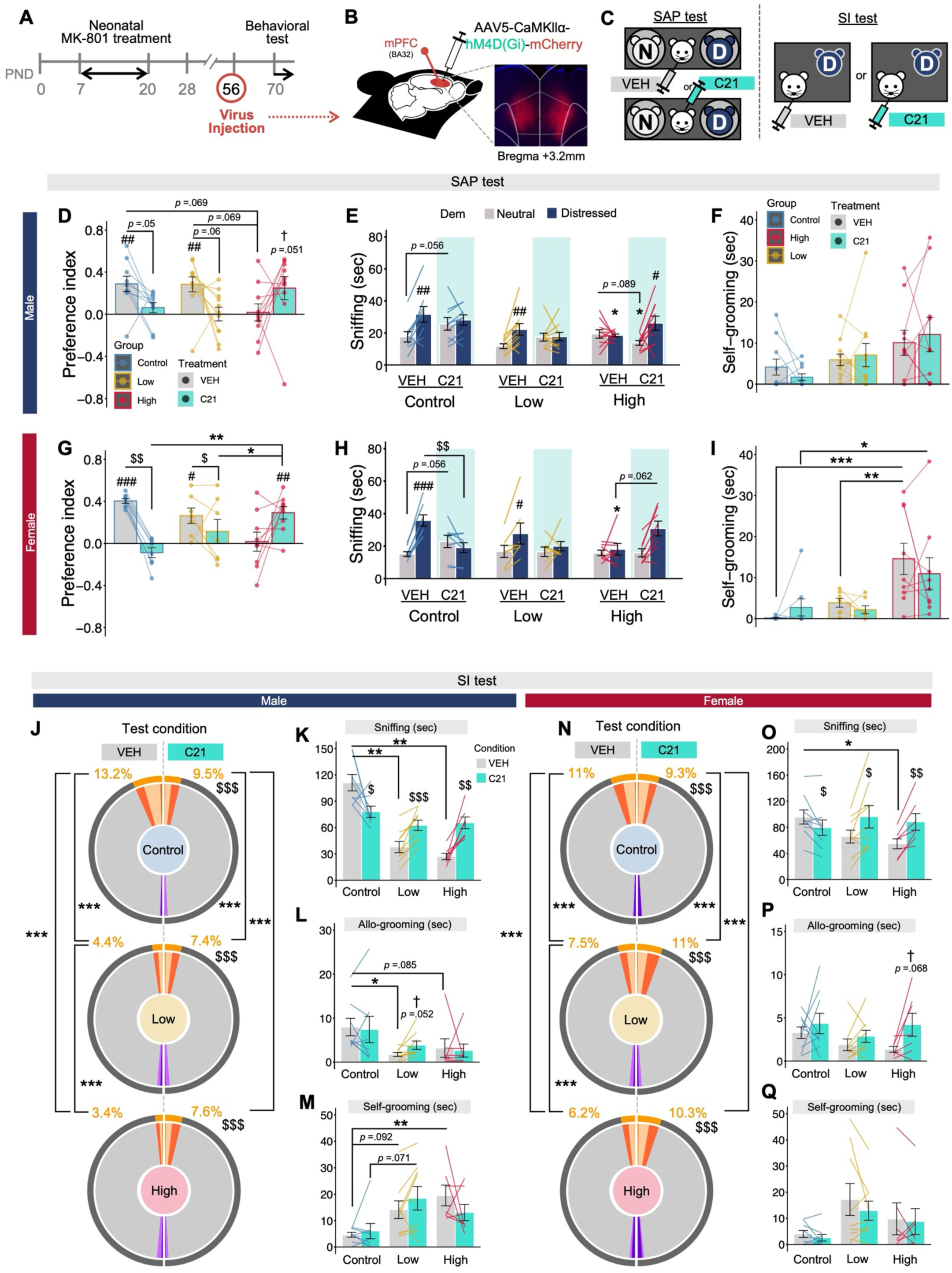
Chemogenetic inhibition of mPFC pyramidal cells partially alleviates social affective impairments in MK-High rats but disrupts them in Control rats. A, Experimental design for the SAP and SI tests with DREADD manipulation. **B** Schematic of viral vector injection. An inhibitory DREADD targeting mPFC pyramidal cells was introduced into the mPFC. **C** Within-subject design for vehicle (VEH) and Compound 21 (C21) treatment conditions in the SAP test (left) and SI test (right). **D–I** Data were obtained from 51 subjects (Control: male n = 9, female n = 8; Low: male n = 10, female n = 7; High: male n = 10, female n = 9). **D–F** SAP test results in males: preference index (**D**), time spent sniffing each demonstrator (**E**), and self-grooming during the test (**F**). **G–I** SAP test results in females: preference index (**G**), time spent sniffing each demonstrator (**H**), and self-grooming during the test (**I**). **J–Q** Data were obtained from a total of 50 subjects (Control: male n = 9, female n = 9; Low: male n = 9, female n = 9; High: male n = 7, female n = 7). **J–M**, SI test results in males: proportion of behavioral categories (**J**), time spent sniffing (**K**), allo-grooming (**L**), and self-grooming (**M**). **N–Q** SI test results in females: proportion of behavioral categories (**N**), time spent sniffing (**O**), allo-grooming (**P**), and self-grooming (**Q**). Bar and line graphs show the mean ± s.e.m. \**p* < 0.05, \*\**p* < 0.01, and \*\*\**p* < 0.001 indicate significant differences between groups. #*p* < 0.05, ##*p* < 0.01, ###*p* < 0.001, and †*p* < 0.10 indicate significant differences or trends between neutral and distressed demonstrators in panels **E**, **H**, and **J–Q**, or from chance level in panels **D** and **G**. $*p* < 0.05, $$*p* < 0.01, and $$$*p* < 0.001 indicate significant differences between vehicle- and C21-treated conditions.

**Figure 3.**
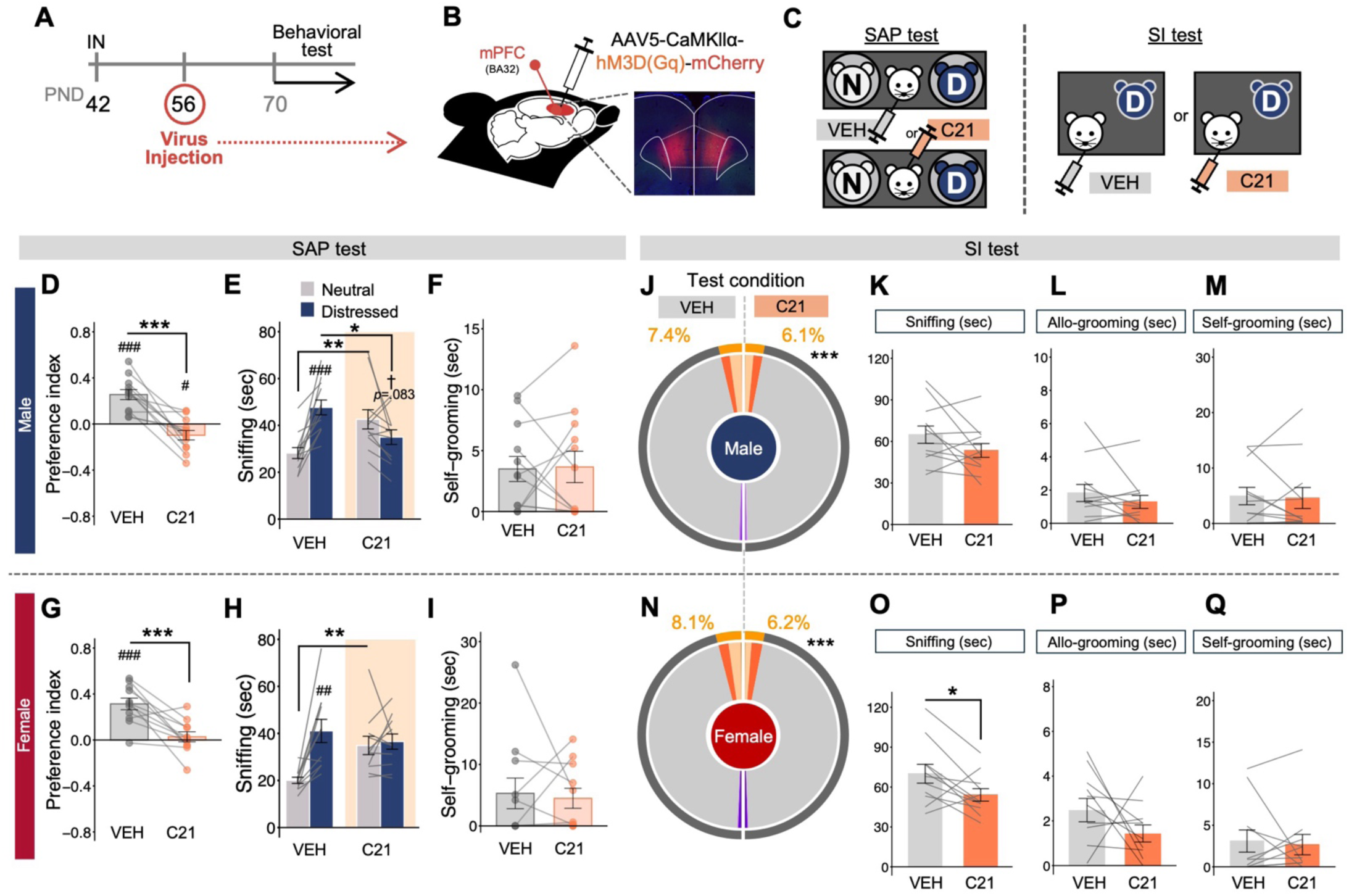
Activation of mPFC pyramidal cells replicates social affective impairments observed in MK-801-treated rats. **A** Experimental timeline for DREADD-mediated activation of mPFC pyramidal cells. **B** Schematic of viral vector injection. An excitatory DREADD targeting mPFC pyramidal cells was introduced into the mPFC. **C** Within-subject design for vehicle (VEH) and Compound 21 (C21) treatment conditions in the SAP test (left) and SI test (right). **D–Q** Data were obtained from 23 subjects (male n = 12, female n = 11). **D–F** SAP test results in males: preference index (**D**), time spent sniffing each demonstrator (**E**), and self-grooming during the test (**F**). **G–I** SAP test results in females: preference index (**G**), time spent sniffing each demonstrator (**H**), and self-grooming during the test (**I**). **J–M** SI test results in males: proportion of behavioral categories (**J**), time spent sniffing (**K**), allo-grooming (**L**), and self-grooming (**M**). **N–Q** SI test results in females: proportion of behavioral categories (**N**), time spent sniffing (**O**), allo-grooming (**P**), and self-grooming (**Q**). Bar and line graphs show the mean ± s.e.m. \**p* < 0.05, \*\**p* < 0.01, and \*\*\**p* < 0.001 indicate significant differences between vehicle- and C21-treated conditions. †*p* < 0.10, #*p* < 0.05, ##*p* < 0.01, and ###*p* < 0.001 indicate trends or significant differences between neutral and distressed demonstrators in panels **E**, **H**, and **J–Q**, or from chance level in the preference index in panels **D** and **G**.

For the SAP test with DREADD manipulation, sniffing behavior was analyzed using a three-way ANOVA with group, affective state of the demonstrators, and DREADD condition (vehicle-treated vs. C21-treated) as factors, followed by multiple comparisons with Bonferroni correction (for the experiment shown in Figure 4 only, a two-way ANOVA with demonstrator and treatment as factors was used). Preference indices were compared against chance level using one-sample t-tests and then analyzed using a two-way ANOVA with group and DREADD condition as factors, followed by Bonferroni post hoc tests (For the experiment shown in Figure 4, a one-way ANOVA with treatment as a factor was used).

**Figure 4.**
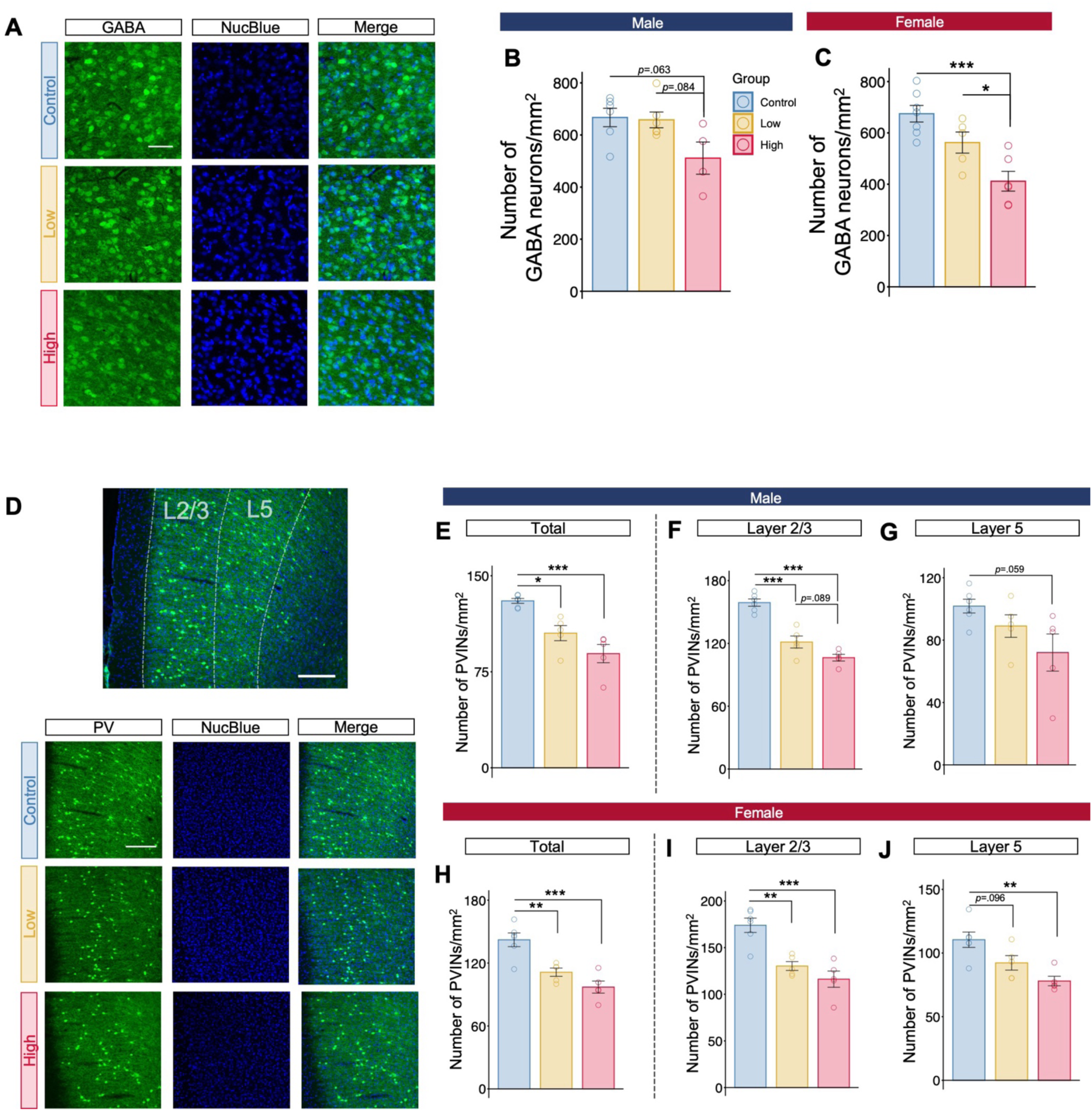
Neonatal NMDAr antagonism leads to persistent reductions in mPFC inhibitory neuron populations, particularly PVINs. **A** Representative immunofluorescence images of GABA-positive neurons. **B**, **C** Density of GABA-positive neurons in the mPFC in males (**B**) and females (**C**). **D** Representative immunofluorescence images of PVINs. **E–G** PVIN density in the mPFC in males: total mPFC (**E**), layers II/III (**F**), and layer V (**G**). **H–J** PVIN density in the mPFC in females: total mPFC (**H**), layers II/III (**I**), and layer V (**J**). Scale bars indicate 10 µm in **A**, 100 µm in the upper image in **D**, and 200 µm in the lower image in **D**. Bar and line graphs show the mean ± s.e.m. \**p* < 0.05, \*\**p* < 0.01, and \*\*\**p* < 0.001 indicate significant differences between groups.

Similarly, behaviors during the SI test with DREADD manipulation were analyzed using a two-way ANOVA with group and treatment (Vehicle or Compound 21) as factors, followed by Bonferroni post hoc tests. For the analysis of the DREADD experiment without the group factor (Figure 4), following a one-way ANOVA with treatment as a factor, pairwise t-test with Bonferroni correction was performed.

## Results

### Rats chronically treated with an NMDAr antagonist MK-801 during the neonatal period exhibit impaired behavioral responses toward a distressed conspecific

All subjects received neonatal treatment from postnatal day 7 to 20, and their social and emotional behaviors were assessed during adulthood (Figure 1A).

In the elevated-plus maze test, MK-High rats showed increases in the time spent in the open-arms, the number of open-arm entries, head dips, and reaching (Figure S1B-J). This indicates reduced anxiety and heightened exploratory motivation in the MK-High rats, rather than increased anxiety-related phenotypes. In the open-field test, there were no significant differences in behavioral indices among groups in either sex, except that male MK-Low rats tended to spend less time in the center area (Figure S1L-Q). These results demonstrate that NMDAr dysfunction during neonatal stage does not impair general emotional behaviors.

Next, we assessed social affective behaviors in the MK-801-treated rats. In the social affective preference (SAP) test (Figure 1B, D), Control and MK-Low rats spent more time sniffing distressed conspecifics injected with LiCl than neutral ones, whereas MK-High rats did not exhibit this tendency (Figure 1F, J, Figure S2K, M). Consistently, preference indices indicated significant preferences for a distressed conspecific in the Control and MK-Low, but not in MK-High groups (Figure 1E, I). Furthermore, both MK-Low and MK-High rats exhibited increased self-grooming during the SAP test (Figure 1G, K). In males, self-grooming duration was negatively correlated with the duration of sniffing toward distressed demonstrators (Figure 1H), whereas no such correlation was observed in females (Figure 1L). Distance travelled during the SAP test did not differ among groups (Figure S2J, L). Notably, neonatal treatment did not influence sniffing, self-grooming or locomotion behaviors during the social habituation phase (Figure S2A-H). Similar behavioral tendencies during the SAP test were also observed when subjects faced distressed demonstrators exposed to a low dose of LiCl or restraint stress (Figures S3, S4).

In the social interaction test (SI test; Figure 1M), all groups spent more time engaging in social behavior with a distressed conspecific than with a neutral one. However, MK-Low and MK-High rats spent less time engaging in social behavior compared to Control rats under both conditions (Figure 1N, R). Consistent with the SAP test, MK-Low and MK-High rats showed reduced sniffing and allogrooming toward distressed demonstrators compared with

Control rats (Figure 1O, P, S, T, Figure S2N, O, Q, R). Increased self-grooming was also observed in MK-801-treated rats during the SI test, similar to the SAP test (Figure 1Q, U). No significant differences in digging behaviors were observed among groups in either sex (Figure S2P, S).

These findings indicate that neonatal NMDAr blockade impairs social affective responses toward conspecifics in an aversive state, as reflected by reduced prosocial behaviors and increased self-grooming in social contexts.

### Neonatal NMDAr antagonism does not influence social novelty or odor preferences

To examine whether neonatal NMDAr antagonism impacts other aspects of social abilities, we next assessed social novelty discrimination and odor preference behaviors (Figure S5A, E). In the social novelty test, MK-801-treated rats exhibited intact discrimination performance (Figure S5 B-D). Similarly, neonatal MK-801 treatment did not alter preference for odors collected from distressed conspecifics (Figure S5 F, G). These results suggest that impaired social affective behaviors observed in MK-treated rats are specifically expressed when distressed conspecifics are physically present, and do not stem from a general inability to recognize social information.

### Chemogenetic inhibition of mPFC pyramidal cells partially ameliorates social affective deficits in MK-High rats but disrupts them in Control rats

We hypothesized that neonatal NMDAr hypofunction disrupts the development of cortical inhibitory systems, thereby biasing mPFC circuits toward an excessive excitatory state and contributing to schizophrenia-related social affective deficits. To test this hypothesis, we examined how the chemogenetic inhibition of mPFC pyramidal cells influences social affective behaviors. Subjects underwent stereotaxic surgery for the viral transduction of an inhibitory DREADD selectively targeting mPFC pyramidal cells (Figure 2A, B). Each subject was tested under two conditions, vehicle and Compound 21 (C21), in both the SAP and SI tests (Figure 2C).

Under the vehicle condition, Control and MK-Low rats, but not MK-High rats, spent more time exploring distressed demonstrators than neutral ones (Figure 2E, H). Notably, this pattern was reversed under the C21 condition: mPFC inhibition restored the preference for distressed demonstrators in MK-High rats, whereas it disrupted these preferences in Control and MK-Low rats (Figure 2D, G). These changes were primarily attributable to altered sniffing toward neutral demonstrators in males (Figure 2E, Figure S6B), whereas in females, they were mainly driven by altered exploration of distressed demonstrators (Figure 2H, Figure S6D). Although mPFC inhibition significantly modulated social exploratory behavior, it did not affect self-grooming or locomotion in the SAP test (Figure 2F, I, Figure S6A, C).

Similar behavioral modulation was observed in the SI test. Under the vehicle condition, MK-Low and MK-High rats showed lower levels of social behavior than Control rats. C21 administration increased the time spent engaging in social behavior in MK-Low and MK-High rats, whereas it decreased social behavior in Control rats (Figure 2J, N). These changes were largely reflected in sniffing behavior, which increased in MK-Low and MK-High rats but decreased in Control rats following C21 administration (Figure 2K, O, Figure S6E, F, H, I). Other behaviors were not significantly modulated by C21 administration, except for a tendency toward increased allogrooming in the male MK-Low and female MK-High groups (Figure 2L, M, P, Q, Figure S6G, J).

In contrast to the chemogenetic inhibition experiment, C21 treatment did not modulate any behaviors during the SAP test in either the Control or MK-High groups in experiments using control virus injections (Figure S7).

Together, these findings indicate that chemogenetic inhibition of mPFC pyramidal cells bidirectionally modulates social affective behaviors depending on neonatal treatment history, partially ameliorating social affective deficits in MK-801-treated rats while disrupting normal social affective responses in Control rats.

### Activation of mPFC pyramidal cells induces social affective impairments

To determine whether an excitatory-biased cortical state is sufficient to induce social affective impairments resembling those observed in MK-801-treated rats, we assessed the effects of chemogenetic activation of mPFC pyramidal cells on social affective behaviors in naive rats. Naive rats were obtained at 6 weeks of age and received viral injections targeting mPFC pyramidal cells during adulthood (Figure 3A, B).

In the SAP test (Figure 3C), vehicle-treated rats showed a preference for distressed demonstrators, whereas this preference was abolished by C21 administration (Figure 3D, G). Consistently, increased sniffing toward distressed demonstrators was observed under the vehicle condition but was lost following C21 administration (Figure 3E, H, Figure S8B, D). Self-grooming and locomotion were not modulated by C21 administration (Figure 3F, I, Figure S8A, C).

In the SI test, the proportion of social behaviors was reduced by C21 administration (Figure 3J, N). In males, total sniffing time did not differ between treatments (Figure 3K); however, further analysis by a body region revealed that anogenital sniffing, but not body/head sniffing, was decreased by C21 administration (Figure S8E, F). In females, rats under the C21 condition exhibited less sniffing than under the vehicle condition, especially in anogenital area (Figure 3O, Figure S8H, I). Similar to the previous experiment, mPFC activation did not influence allogrooming, self-grooming, or digging behaviors (Figure 3L, M, P, Q, Figure S8G, J).

These results indicate that chemogenetic activation of mPFC pyramidal cells disrupts social affective responses toward distressed conspecifics, supporting the possibility that an excitatory-biased mPFC state contributes to NMDAr blockade-like social affective impairments.

### Neonatal NMDAr antagonism leads to persistent reductions in mPFC inhibitory neuron populations, particularly PVINs

To examine the effects of neonatal chronic MK-801 treatment on the inhibitory system in the mPFC, we performed immunofluorescence staining for GABA-positive cells (Figure 4A). The MK-High group had fewer GABA-positive cells than in the Control and MK-Low groups, particularly in females (Figure 4B, C).

Given the particular relevance of PVINs to schizophrenia-related cortical dysfunction, we next examined PVINs in the mPFC (Figure 4D). Compared with the Control group, the number of PVINs in the mPFC was significantly reduced in both MK-Low and MK-High groups in both sexes (Figure 4E, H). Layer-specific analysis in males revealed significant reductions in layer II / III PVINs in both the MK-Low and MK-High groups compared with the Control group, with a trend toward a further reduction in the MK-High group compared with the MK-

Low group (Figure 4F). In layer V, no significant group differences were detected, although the number of PVINs in the MK-High group tended to be lower than that in the Control group (Figure 4G). In females, PVINs in the MK-Low and MK-High groups were significantly reduced compared with the Control group across cortical layers, although the difference between the Control and MK-Low groups in layer V remained at a trend level (Figure 4I, J).

These results suggest that neonatal MK-801 treatment induces persistent alterations in the mPFC inhibitory system, with prominent reductions in PVINs, particularly in layer II / III.

### PVIN-specific chemogenetic activation in the mPFC alleviates social affective impairments in MK-High rats

Based on the observed vulnerability of mPFC PVINs to neonatal MK-801 treatment, we next examined whether PVIN-specific activation in the mPFC could modulate socio-affective behaviors. To this end, we injected an excitatory DREADD virus selectively targeting PVINs into the mPFC (Figure 5A, B, Figure S9A).

**Figure 5.**
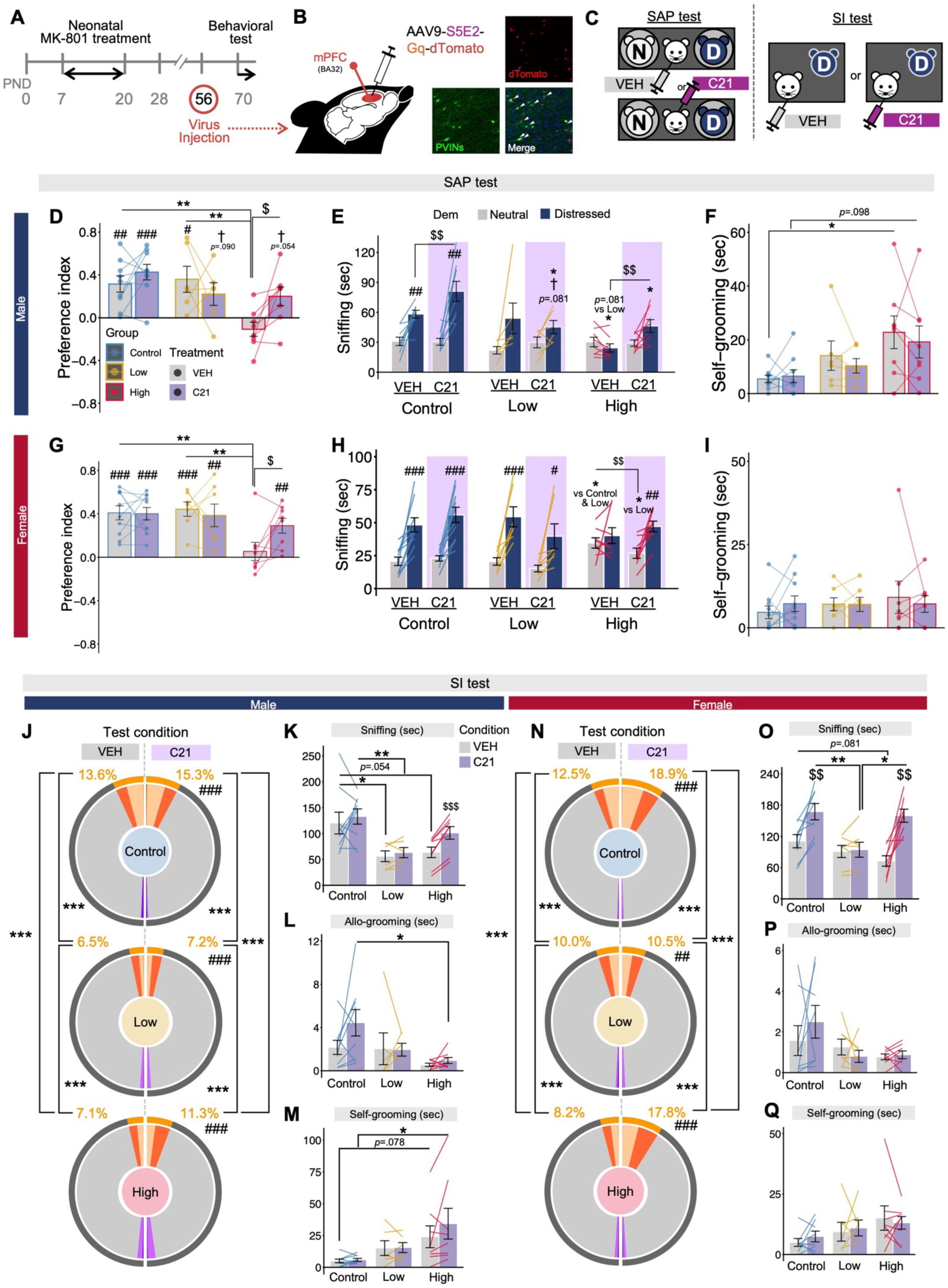
Chemogenetic activation targeting PVINs in the mPFC ameliorates social affective impairments in MK-High rats. **A** Experimental design for the SAP and SI tests with DREADD manipulation. **B** Schematic of viral vector injection. An excitatory DREADD selectively targeting PVINs was introduced into subjects’ mPFC. **C** Within-subject design for vehicle (VEH) and Compound 21 (C21) treatment conditions in the SAP test (left) and SI test (right). **D–I** Data were obtained from 48 subjects (Control: male n = 9, female n = 10; Low: male n = 6, female n = 7; High: male n = 8, female n = 8). **D–F** SAP test results in males: preference index (**D**), time spent sniffing each demonstrator (**E**), and self-grooming during the test (**F**). **G–I** SAP test results in females: preference index (**G**), time spent sniffing each demonstrator (**H**), and self-grooming during the test (**I**). **J–Q** Data were obtained from 46 subjects (Control: male n = 9, female n = 8; Low: male n = 6, female n = 7; High: male n = 8, female n = 8). **J–M**, SI test results in males: proportion of behavioral categories (**J**), time spent sniffing (**K**), allo-grooming (**L**), and self-grooming (**M**). **N–Q** SI test results in females: proportion of behavioral categories (**N**), time spent sniffing (**O**), allo-grooming (**P**), and self-grooming (**Q**). Bar and line graphs show the mean ± s.e.m. \**p* < 0.05, \*\**p* < 0.01, and \*\*\**p* < 0.001 indicate significant differences between groups. #*p* < 0.05, ##*p* < 0.01, ###*p* < 0.001, and †*p* < 0.10 indicate significant differences or trends between neutral and distressed demonstrators in panels **E**, **H**, and **J–Q**, or from chance level in panels **D** and **G**. $*p* < 0.05, $$*p* < 0.01, and $$$*p* < 0.001 indicate significant differences between vehicle-and C21-treated conditions.

**Figure 6.**
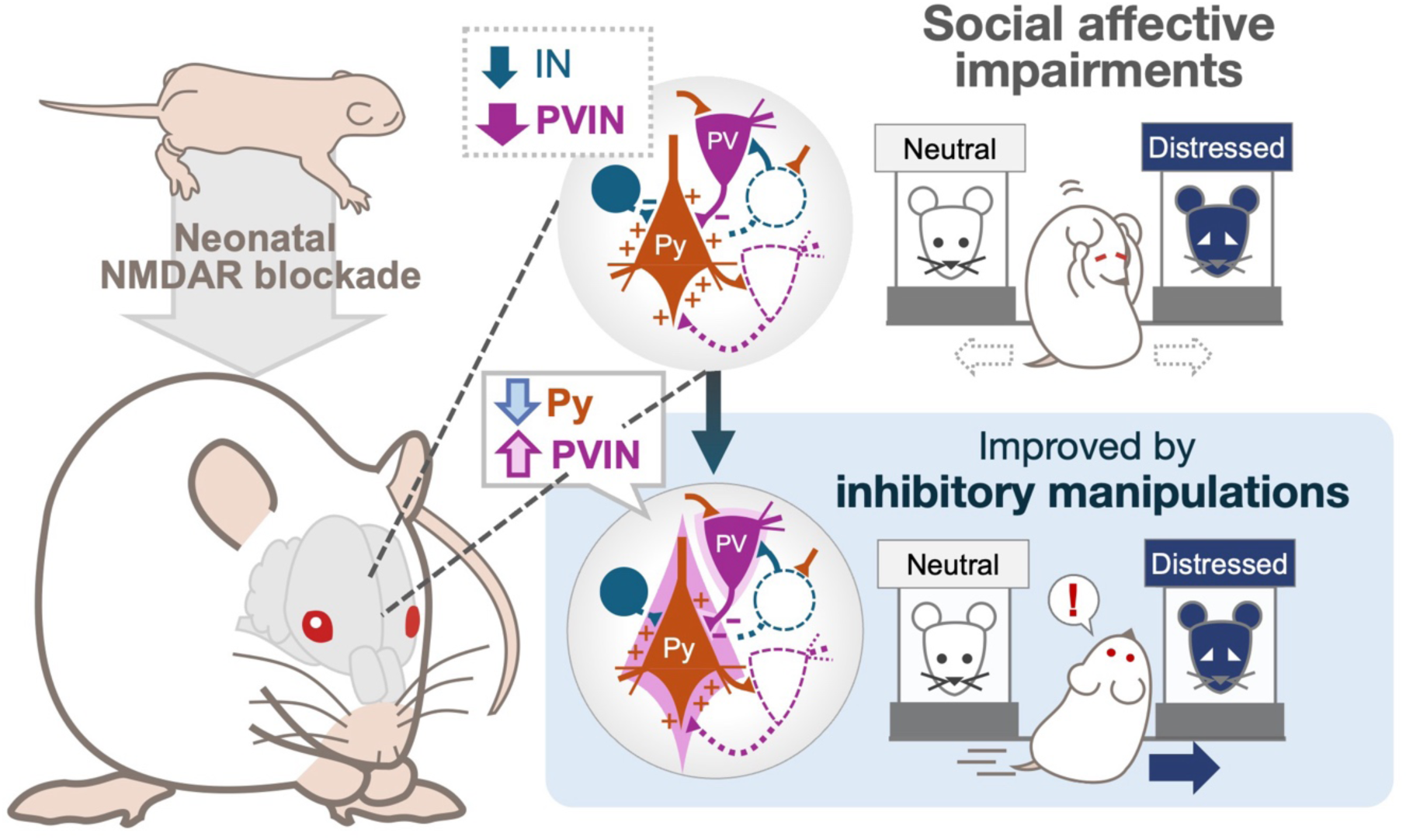
Graphical summary of this study. Chronic neonatal blockade of NMDArs contribute to social affective impairments in adulthood through disruption of inhibitory circuit development, particularly PVIN-mediated inhibitory circuits.

In the SAP test (Figure 5C), C21 treatment significantly increased the preference for distressed demonstrators in the MK-High group, whereas no significant effects were observed in the Control or MK-Low groups (Figure 5D, G, Figure S9C, E). These effects were reflected by increased sniffing toward distressed demonstrators in males (Figure 5E), whereas in females, they were attributable to decreased sniffing toward neutral demonstrators (Figure 5H). Apart from these changes, C21 treatment did not consistently alter sniffing time toward either demonstrator in the Control or MK-Low groups, except for an increase in sniffing toward distressed demonstrators in male Control rats (Figure 5E, H). C21 treatment did not affect self-grooming or locomotion behaviors (Figure 5F, I, Figure S9B, D).

In the SI test, chemogenetic activation of mPFC PVINs increased social behavior across all groups, although significant between-group differences remained (Figure 5J, N). C21 treatment significantly increased sniffing behavior in the MK-High group (Figure 5K, O, Figure S9G, I, J). This effect was also observed in Control females, but not in Control males or MK-Low rats (Figure 5K, O, Figure S9F, G, I, J). Other behaviors, including allogrooming and self-grooming, were not affected by C21 treatment (Figure 5L, M, P, Q, Figure S9H, K).

Together, these findings indicate that PVIN-specific chemogenetic activation in the mPFC ameliorates social affective impairments in MK-High rats.

## Discussion

In the present study, we found that rats neonatally treated with MK-801, a developmental model relevant to schizophrenia, exhibited impaired social affective responses characterized by reduced social exploration of distressed conspecifics. These behavioral deficits were ameliorated by chemogenetic inhibition of pyramidal cells or activation of PVINs in the mPFC. Together, these findings suggest that impaired inhibitory regulation within the mPFC contributes to social affective dysfunction and raise the possibility that PVIN-mediated inhibitory circuits represent a potential therapeutic target for alleviating social deficits associated with schizophrenia.

Specifically, chronic neonatal MK-801 treatment reduced social exploration of and preference for a distressed conspecific while increasing self-grooming behavior. These behavioral alterations resemble certain aspects of social dysfunction observed in patients with schizophrenia [3, 35]. Consistent with our findings, impaired social affective preference has been reported in mice with neonatal NMDAr subunit knockout [17] or chronic PCP treatment [36]. These convergent findings support the idea that disruption of NMDAr-mediated signaling during early development contributes to persistent social affective deficits in adulthood. In addition, MK-801-treated rats in the present study, exhibited excessive self-grooming, an anxiety-or stress-related behavior in rodents, in both the SAP and SI tests. This phenotype may reflect altered affective responses in social contexts, as observed in schizophrenia [37]. Thus, the reduced sniffing of distressed conspecifics and heightened self- grooming behavior observed in MK-801-treated rats indicate that chronic neonatal NMDAr blockade alters affective responses when animals encounter distressed conspecifics.

The DREADD-based circuit manipulations and histological analyses performed in this study consistently support the hypothesis that insufficient inhibitory control within the mPFC contributes to schizophrenia-related social affective deficits. Our findings suggest that appropriate regulation of mPFC pyramidal cell activity by inhibitory interneurons, particularly PVINs, is critical for the expression of social affective responses. We performed two manipulations designed to enhance inhibitory control over mPFC output: chemogenetic inhibition of pyramidal cells and activation of PVINs. Both manipulations restored social affective preference in the MK-High group. Among multiple classes of inhibitory interneurons in the mPFC, PVINs are particularly important for controlling pyramidal cell output because they preferentially target the perisomatic region of pyramidal cells [38]. In addition, chemogenetic activation of mPFC pyramidal cells in naïve rats reproduced impairments in social affective preference similar to those observed in MK-801-treated rats. Together, these findings suggest that social affective preference depends on maintaining mPFC pyramidal cell activity within an appropriate range through inhibitory regulation.

One possible explanation for the restorative effect of PVIN activation in the MK-High group is that direct activation of PVINs compensated for a broader reduction in inhibitory control within the mPFC. Our immunohistochemical data demonstrated that the MK-High group exhibited a reduction in GABA-positive cells within the mPFC, with a prominent decrease in PVINs. This finding is consistent with previous reports of PVIN abnormalities in schizophrenia [23] and in animal models of chronic NMDAr blockade [31], as well as with the present observation that circuit manipulations enhancing inhibitory regulation improved social affective deficits. PVINs constitute a major source of inhibitory input onto pyramidal cells within local mPFC circuits [39]. In contrast, other types of inhibitory interneurons, including SST-and VIP-positive cells [40], do not primarily inhibit pyramidal cell bodies; rather, their inhibitory effects on pyramidal cell activity may be mediated through PVIN-dependent circuit mechanisms. In the MK-High group, both PVINs and the overall population of GABAergic cells were reduced, indicating that inhibitory interneuron populations other than PVINs may also have been affected and could contribute to social affective deficits. Nevertheless, given the direct perisomatic inhibitory control exerted by PVINs over pyramidal cells [39], the restoration of social affective preference by PVIN-specific activation suggests that artificial PVIN activation rebalanced pyramidal cell activity by compensating for reduced inhibitory control within the mPFC.

Importantly, inhibition of mPFC pyramidal cells disrupted social affective preference and reduced exploration of a distressed conspecific in the Control and MK-Low groups. These findings indicate that excessive suppression of pyramidal cell activity, like excessive activation, can impair appropriate social affective responses. Thus, social affective behavior likely requires balanced regulation of mPFC pyramidal cell activity, rather than a simple increase or decrease in cortical excitability. In contrast, activation of mPFC PVINs did not impair social affective preference in the Control group; instead, it increased social exploration of distressed conspecifics in the SAP and SI tests. This result is consistent with a previous study showing that PVIN inhibition decreases social exploration without directly modulating social affective preference [41]. Indeed, PVIN activity contributes to perceptual, attentional, and cognitive functions [42]. Optogenetic and chemogenetic activation of PVINs enhances these functions not only in animal models of schizophrenia [33, 43] but also in healthy animals [44]. Therefore, activation of PVINs may enhance attention toward distressed conspecifics beyond baseline levels, which can explain why chemogenetic activation of PVINs in the Control group increased exploration of distressed conspecifics.

Notably, the reduction in PVINs was most evident in layer II/III of the mPFC. Layer II/III PVIN abnormalities have been reported in patients with schizophrenia [45] and in a PCP-treated animal model [43]. Layer II/III pyramidal cells give rise to subcortical projections, including those to the basolateral amygdala and nucleus accumbens [46, 47], regions that are crucial for social preference [48, 49]. Given this, the layer-specific abnormalities in the mPFC observed in both previous studies and the present work may provide important clues for elucidating the neural circuits underlying schizophrenia-associated social affective impairments.

Another interesting finding was the presence of sex-related differences in the SAP test, particularly in the effects of DREADD-based circuit manipulations. Sex differences in social cognition and social behavior are widely recognized across mammalian species, including humans [50]. In rodents, previous studies have reported sex-dependent differences in the quantity and quality of social behaviors [51], as well as in neural activity during social situations [52]. Recent work has further shown that behavioral responses toward distressed conspecifics differ between males and females, and that neural circuits involving the prefrontal cortex contribute to such sex-specific behavioral strategies [53].

In the present study, the overall pattern of social affective behavior and the general direction of the effects induced by circuit manipulations were broadly similar between males and females. However, closer inspection of the SAP test revealed subtle but consistent sex-related differences. During chemogenetic inhibition of mPFC pyramidal cells, males mainly showed changes in exploration time toward the neutral conspecific, whereas females showed changes primarily in exploration time toward the distressed conspecific. In contrast, during PVIN activation, this pattern appeared to be reversed: males showed recovery of exploration toward the distressed conspecific, whereas females showed changes in exploration toward the neutral conspecific. These findings raise the possibility that the behavioral components underlying social affective preference differ between males and females. In other words, preference for a distressed conspecific may be generated through distinct behavioral strategies, such as enhanced interest in the distressed conspecific or reduced interest in the neutral conspecific, depending on sex.

The relative contribution of excitatory and inhibitory circuit mechanisms to these behavioral components may also be sex-dependent. However, the exact mechanisms underlying these sex-related differences remain to be determined. In addition to previous studies suggesting that sex differences in neural circuit organization contribute to distinct strategies for social affective behaviors in mice [53], recent evidence indicates that PVINs themselves exhibit sex-dependent cellular and functional properties [54]. Thus, the sex-related differences observed in the present study may reflect differences at multiple levels, ranging from intrinsic cellular properties and synaptic organization to broader circuit-level dynamics. Future studies integrating molecular, cellular, circuit, and behavioral analyses will be necessary to clarify how sex influences mPFC-dependent social affective responses.

Although the present findings highlight the importance of mPFC inhibitory control in social affective behavior, manipulation of mPFC circuits did not fully normalize all behavioral abnormalities observed in neonatally MK-801-treated rats. Chemogenetic manipulation of mPFC pyramidal cells or PVINs improved abnormalities in social exploration of distressed conspecifics, but did not significantly alter self-grooming during the SAP test or allo-grooming behavior in the SI test. This dissociation suggests that mPFC inhibitory circuits preferentially regulate exploratory components of social affective behavior, whereas anxiety-like, stress-related, or more complex social behavioral abnormalities may involve partially distinct neural mechanisms.

The mPFC is a key cortical region involved in the regulation of social behavior [55], which may explain why mPFC circuit manipulations markedly influenced exploratory responses toward distressed conspecifics. However, broader behavioral improvements beyond social exploration may require modulation of distributed cortical and subcortical circuits. Increased self-grooming during exposure to a distressed conspecific may involve emotion-related regions such as the amygdala [53, 56], whereas complex social behaviors such as allo-grooming and helping-like behavior may involve cortical regions including the anterior cingulate cortex [57–59], both of which exhibit functional abnormalities in patients with schizophrenia [35, 60]. The habenula, a region involved in aversive processing and social avoidance [61], also shows structural and functional abnormalities in schizophrenia [62], suggesting its potential relevance to schizophrenia-related social dysfunction. Thus, while our findings support a central role for mPFC inhibitory control in social affective exploration, abnormalities in other behavioral domains may reflect dysfunction across broader neural systems. Further studies examining cortical and subcortical regions, as well as their anatomical and functional interactions with the mPFC, will be important for understanding the broader circuit mechanisms underlying schizophrenia-related social behavioral abnormalities.

Finally, neonatal NMDArs blockade may induce widespread alterations in brain development. NMDAr signaling during early postnatal periods is critical for the maturation of synaptic connections and neural circuits [63], suggesting that neonatal MK-801 treatment affects not only the mPFC and the regions discussed above, but also broader anatomical and functional networks. Although the present study focused on local mPFC circuits, schizophrenia-related social affective dysfunction may also reflect broader abnormalities across cortical and subcortical circuits, including the amygdala, retrosplenial cortex, and insular cortex, which have been implicated in social affective functions [34, 36, 64–67]. Future studies combining region-specific manipulations with analyses of anatomical and functional connectivity will be necessary to provide a more comprehensive understanding of how developmental NMDAr hypofunction gives rise to circuit abnormalities underlying schizophrenia-related social affective dysfunction.

Overall, our study highlights a link between developmental NMDAr hypofunction and schizophrenia-related social affective impairments in adulthood through alterations in prefrontal PVIN-mediated inhibitory circuits. The finding that activation of prefrontal PVINs improved social affective responses in MK-High rats supports a causal contribution of impaired PVIN-mediated inhibitory control to schizophrenia-related social abnormalities. These results further suggest that neural substrates underlying social dysfunction may, at least in part, overlap with those implicated in cognitive and attentional deficits. Together, these findings provide new insight into the circuit mechanisms of social affective dysfunction and identify prefrontal inhibitory circuits as a potential target for future therapeutic approaches.

## Acknowledgements

We thank Prof. Hiroyuki Hioki (Juntendo University Graduate School of Medicine), Prof. Megumu Takahashi (Juntendo University Graduate School of Medicine), and Prof. Yasuhiro R. Tanaka (Tamagawa University) for technical advice on the experimental procedures. We also thank Prof. Aki Takahashi (University of Tsukuba) and Dr. Koshiro Mitsui (Toyo University) for valuable advice and insightful discussions regarding the experimental design and interpretation of the results.

## Author contributions

R.T. conceptualized the study, developed the methodology, performed the experiments, conducted formal analyses, acquired funding, visualized the data, and wrote the original draft. N.H., C.G.d.R., T.X., and K.I. performed experiments and reviewed and edited the manuscript.

T.O. contributed to methodology and reviewed and edited the manuscript. M.H. acquired funding and reviewed and edited the manuscript. K.Y. supervised the study and contributed to reviewing and editing the manuscript. M.T. conceptualized and supervised the study, performed experiments, acquired funding, and reviewed and edited the manuscript. All authors read and approved the final manuscript.

## Ethics approval

This study was performed in accordance with the Guidelines for the Care and Use of Laboratory Animals at the University of Tsukuba. All animal experiments were approved by the University of Tsukuba Animal Ethics Committee (Authorization number: 24-392, 25-389, and 26-386) and the University of Tsukuba Safety Committee for Recombinant DNA Experiments (Authorization number: 240218 and 250227).

## References

1. American Psychiatric Association. Diagnostic and Statistical Manual of Mental Disorders 5th edn (American Psychiatric Publication, 2013).

2. Neill JC et al. Animal models of cognitive dysfunction and negative symptoms of schizophrenia: focus on NMDAr antagonism. Pharmacol. Ther. 128, 419–432 (2010).

3. Green MF, Horan WP & Lee J. Social cognition in schizophrenia. Nat. Rev. Neurosci. 16, 620–631 (2015).

4. Derntl B et al. Generalized deficit in all core components of empathy in schizophrenia. Schizophr. Res. 108, 197–206 (2009).

5. Yizhar O & Levy DR. The social dilemma: prefrontal control of mammalian sociability. Curr. Opin. Neurobiol. 68, 67–75 (2021).

6. Comer AL et al. Increased expression of schizophrenia-associated gene C4 leads to hypoconnectivity of prefrontal cortex and reduced social interaction. PLoS Biol. 18, e3000604 (2020).

7. Sánchez-González A et al. Increased thin-spine density in frontal cortex pyramidal neurons in a genetic rat model of schizophrenia-relevant features. Eur. Neuropsychopharmacol. 44, 79–91 (2021).

8. Tapias-Espinosa C et al. Decreased activation of parvalbumin interneurons in the medial prefrontal cortex in intact inbred Roman rats with schizophrenia-like reduced sensorimotor gating. Behav. Brain Res. 437, 114141 (2023).

9. Uno Y & Coyle JT. Glutamate hypothesis in schizophrenia. Psychiatry Clin. Neurosci. 73, 204–215 (2019).

10. Sommer IEC et al. The treatment of hallucinations in schizophrenia spectrum disorders. Schizophr. Bull. 38, 704–714 (2012).

11. Singh T et al. Rare coding variants in ten genes confer substantial risk for schizophrenia. Nature 604, 509–516 (2022).

12. Clapcote SJ et al. Behavioral phenotypes of Disc1 missense mutations in mice. Neuron 54, 387–402 (2007).

13. Mei L & Xiong WC. Neuregulin 1 in neural development, synaptic plasticity and schizophrenia. Nat. Rev. Neurosci. 9, 437–452 (2008).

14. Furuie H, Yamada K & Ichitani Y. Differential effects of N-methyl-D-aspartate receptor blockade during the second and third postnatal weeks on spatial working and reference memory in adult rats. Brain Res. 1721, 146339 (2019).

15. Kawabe K & Miyamoto E. Effects of early postnatal MK-801 treatment on behavioral properties in rats: differences according to treatment schedule. Behav. Brain Res. 370, 111926 (2019).

16. Brigman JL. Effects of subchronic phencyclidine treatment on social behaviors and operant discrimination and reversal learning in C57BL/6J mice. Front. Behav. Neurosci. 3, 49 (2009).

17. Uehara JM, Gomez Acosta M, Bello EP & Belforte JE. Early postnatal NMDAr ablation in cortical interneurons impairs affective state discrimination and social functioning. Neuropsychopharmacology 50, 1022–1033 (2025).

18. O’Tuathaigh CMP et al. Phenotypic characterization of spatial cognition and social behavior in mice with ‘knockout’ of the schizophrenia risk gene neuregulin 1. Neuroscience 147, 18–27 (2007).

19. Takagi A et al. Disrupted-in-Schizophrenia 1 regulates spines of the glutamate synapse via Rac1. Nat. Neurosci. 13, 327–332 (2010).

20. Nakazawa K et al. GABAergic interneuron origin of schizophrenia pathophysiology. Neuropharmacology 62, 1574–1583 (2012).

21. Nakazawa K, Jeevakumar V & Nakao K. Spatial and temporal boundaries of NMDA receptor hypofunction leading to schizophrenia. npj Schizophr. 3, 7 (2017).

22. De Jonge JC, Vinkers CH, Hulshoff Pol HE & Marsman A. GABAergic mechanisms in schizophrenia: linking postmortem and in vivo studies. Front. Psychiatry 8, 118 (2017).

23. Hashimoto T et al. Gene expression deficits in a subclass of GABA neurons in the prefrontal cortex of subjects with schizophrenia. J. Neurosci. 23, 6315–6326 (2003).

24. Gonzalez-Burgos G, Hashimoto T & Lewis DA. Alterations of cortical GABA neurons and network oscillations in schizophrenia. Curr. Psychiatry Rep. 12, 335–344 (2010).

25. Druga R, Salaj M & Al-Redouan A. Parvalbumin-positive neurons in the neocortex: a review. Physiol. Res. 72, S173–S191 (2023).

26. Sohal VS, Zhang F, Yizhar O & Deisseroth K. Parvalbumin neurons and gamma rhythms enhance cortical circuit performance. Nature 459, 698–702 (2009).

27. Senkowski D & Gallinat J. Dysfunctional prefrontal gamma-band oscillations reflect working memory and other cognitive deficits in schizophrenia. Biol. Psychiatry 77, 1010– 1019 (2015).

28. Basar-Eroglu C et al. Working memory related gamma oscillations in schizophrenia patients. Int. J. Psychophysiol. 64, 39–45 (2007).

29. Williams LM et al. Emotion-elicited gamma synchrony in patients with first-episode schizophrenia: a neural correlate of social cognition outcomes. J. Psychiatry Neurosci. 34, 303–313 (2009).

30. Plataki ME et al. Effect of neonatal treatment with the NMDAr antagonist, MK-801, during different temporal windows of postnatal period in adult prefrontal cortical and hippocampal function. Front. Behav. Neurosci. 15, 664972 (2021).

31. Shepard R, Heslin K, Hagerdorn P & Coutellier L. Downregulation of Npas4 in parvalbumin interneurons and cognitive deficits after neonatal NMDAr blockade: relevance for schizophrenia. Transl. Psychiatry 9, 99 (2019).

32. Belforte JE et al. Postnatal NMDAr ablation in corticolimbic interneurons confers schizophrenia-like phenotypes. Nat. Neurosci. 13, 76–83 (2010).

33. Patrono E, Hrůzova K, Svoboda J & Stuchlík A. The role of optogenetic stimulations of parvalbumin-positive interneurons in the prefrontal cortex and the ventral hippocampus on an acute MK-801 model of schizophrenia-like cognitive inflexibility. Schizophr. Res. 252, 198–205 (2023).

34. Rogers-Carter MM et al. Insular cortex mediates approach and avoidance responses to social affective stimuli. Nat. Neurosci. 21, 404–414 (2018).

35. Anticevic A et al. Amygdala recruitment in schizophrenia in response to aversive emotional material: a meta-analysis of neuroimaging studies. Schizophr. Bull. 38, 608–621 (2012).

36. Dautan D et al. Cortico-cortical transfer of socially derived information gates emotion recognition. Nat. Neurosci. 27, 1318–1332 (2024).

37. Birchwood M et al. Social anxiety and the shame of psychosis: a study in first episode psychosis. Behav. Res. Ther. 45, 1025–1037 (2007).

38. Nagy-Pál P et al. Structural organization of perisomatic inhibition in the mouse medial prefrontal cortex. J. Neurosci. 43, 6972–6987 (2023).

39. Ferguson BR & Gao WJ. PV interneurons: critical regulators of E/I balance for prefrontal cortex-dependent behavior and psychiatric disorders. Front. Neural Circuits 12, 37 (2018).

40. Boroujeni KB, Balaram P, Tiesinga P & Womelsdorf T. Prefrontal cortex interneurons and their contributions to attention, working memory, and adaptive behavior. Prog. Neurobiol. 258, 102873 (2026).

41. Scheggia D et al. Somatostatin interneurons in the prefrontal cortex control affective state discrimination in mice. Nat. Neurosci. 23, 47–60 (2020).

42. Lewis DA, Curley AA, Glausier JR & Volk DW. Cortical parvalbumin interneurons and cognitive dysfunction in schizophrenia. Trends Neurosci. 35, 57–67 (2012).

43. Arime Y et al. Activation of prefrontal parvalbumin interneurons ameliorates working memory deficit even under clinically comparable antipsychotic treatment in a mouse model of schizophrenia. Neuropsychopharmacology 49, 720–730 (2024).

44. Kim H, Ährlund-Richter S, Wang X, Deisseroth K & Carlén M. Prefrontal parvalbumin neurons in control of attention. Cell 164, 208–218 (2016).

45. Dienel SJ, Schoonover KE & Lewis DA. Cognitive dysfunction and prefrontal cortical circuit alterations in schizophrenia: developmental trajectories. Biol. Psychiatry 92, 450– 459 (2022).

46. Manoocheri K & Carter AG. Rostral and caudal basolateral amygdala engage distinct circuits in the prelimbic and infralimbic prefrontal cortex. eLife 11, e82688 (2022).

47. Green SM, Nathani S, Zimmerman J, Fireman D & Urs NM. Retrograde labeling illuminates distinct topographical organization of D1 and D2 receptor-positive pyramidal neurons in the prefrontal cortex of mice. eNeuro 7, ENEURO.0194-20.2020 (2020).

48. Huang WC et al. Social behavior is modulated by valence-encoding mPFC-amygdala sub-circuitry. Cell Rep. 32, 107899 (2020).

49. Dölen G, Darvishzadeh A, Huang KW & Malenka RC. Social reward requires coordinated activity of nucleus accumbens oxytocin and serotonin. Nature 501, 179–184 (2013).

50. Eagly AH. Sex Differences in Social Behavior: A Social-role Interpretation (Psychology Press, 1987).

51. An XL et al. Strain and sex differences in anxiety-like and social behaviors in C57BL/6J and BALB/cJ mice. Exp. Anim. 60, 111–123 (2011).

52. Mikosz M, Nowak A, Werka T & Knapska E. Sex differences in social modulation of learning in rats. Sci. Rep. 5, 18114 (2015).

53. Fang S et al. Sexually dimorphic control of affective state processing and empathic behaviors. Neuron 112, 1498–1517.e8 (2024).

54. Fabian CB et al. Parvalbumin interneuron mGlu5 receptors govern sex differences in prefrontal cortex physiology and binge drinking. Neuropsychopharmacology 49, 1861– 1871 (2024).

55. Ko J. Neuroanatomical substrates of rodent social behavior: the medial prefrontal cortex and its projection patterns. Front. Neural Circuits 11, 41 (2017).

56. Hong W, Kim DW & Anderson DJ. Antagonistic control of social versus repetitive self-grooming behaviors by separable amygdala neuronal subsets. Cell 158, 1348–1361 (2014).

57. Burkett JP et al. Oxytocin-dependent consolation behavior in rodents. Science 351, 375– 378 (2016).

58. Song D et al. Mediodorsal thalamus-projecting anterior cingulate cortex neurons modulate helping behavior in mice. Curr. Biol. 33, 4330–4342.e5 (2023).

59. Hernandez-Lallement J et al. Harm to others acts as a negative reinforcer in rats. Curr. Biol. 30, 949–961.e7 (2020).

60. Fujiwara H et al. Anterior cingulate pathology and social cognition in schizophrenia: a study of gray matter, white matter and sulcal morphometry. NeuroImage 36, 1236–1245 (2007).

61. Dodis G, Choi S & Flanigan M. The lateral habenula as a master regulator of innate and learned social behaviors. Psychopharmacology 243, 475–493 (2026).

62. Zhang L et al. Altered volume and functional connectivity of the habenula in schizophrenia. Front. Hum. Neurosci. 11, 636 (2017).

63. Ewald RC & Cline HT. NMDA receptors and brain development. In: Van Dongen AM, editor. Biology of the NMDA Receptor (CRC Press, 2008).

64. Ferretti V et al. Oxytocin signaling in the central amygdala modulates emotion discrimination in mice. Curr. Biol. 29, 1938–1953.e6 (2019).

65. Djerdjaj A et al. Social affective behaviors among female rats involve the basolateral amygdala and insular cortex. PLoS One 18, e0281794 (2023).

66. Fujima S, Sato M, Nakai N & Takumi T. Parvalbumin interneurons in the insular cortex control social familiarity and emotion recognition. Cell Rep. 44, 116085 (2025).

67. Chun EK et al. Insular input to the prelimbic cortex underlies social affective behavior in rats. J. Neurosci. 46, e1747252026 (2026).

