## Supplemental Material 1 for "Neonatal NMDA receptor blockade disrupts prefrontal cortical regulation of social affective behavior: implications for schizophrenia"

### Methods

#### Behavioral experiments.

*Open-field test.* The test was conducted in a room with a light intensity of 65 lux using an open-field apparatus (90 × 90 × 45 cm) made of acrylic. The subject was placed in a corner of the apparatus and allowed to explore freely for 15 minutes. After each test, the apparatus was cleaned with a cloth soaked in a 0.1% sodium hypochlorite solution. The subjects' behaviors during the test were recorded using a camera mounted above the apparatus. Total distance traveled and time spent in the corner area were automatically measured using ANY-maze video tracking software (Version 6.16, Stoelting Co.).

*Elevated plus-maze test.* A grey elevated plus-maze apparatus (open arm: 10 × 50 cm, closed arm: 10 × 50 cm, center platform: 10 × 10 cm) made of plastic was used. The subjects were placed in the center area of the maze and allowed to explore freely for 10 minutes. The apparatus was cleaned with a cloth soaked in a 70 % ethanol solution after each trial. The subjects' behaviors were recorded using a camera mounted above the maze, and the time spent in the open arms and the number of entries into the open arm were manually measured using ANY-maze software.

*Social novelty preference test.* The test was conducted in an open-field apparatus (90 × 90 × 45 cm) under a light intensity of 65 lux. The day before the test, the subjects were habituated to the arena containing two empty cylinders for 20 minutes (arena habituation). On the test day, they were presented with two cylinders containing age- and sex-matched unfamiliar conspecifics. These cylinders were placed diagonally in the arena, and the subjects were allowed to explore the environment freely for 5 minutes (sample phase). After a 5-minute interval, they were placed back into the open-field arena and allowed to explore two cylinders again for 5 minutes (test phase). In the test phase, one of the cylinders was replaced with a new one containing another unfamiliar rat (novel demonstrator). The time spent sniffing each cylinder during both the sample and test phases was measured. Based on the sniffing time during the test phase, the novelty preference index was calculated by the formula below:

$$Novelty\ preference\ index = \frac{Sniffing\ time_{novel} - Sniffing\ time_{familiar}}{Sniffing\ time_{novel} + Sniffing\ time_{familiar}}$$

*Odor preference test.* A 3-chambered box made of acryl and two cylinders were used. Prior to the test, the subjects underwent arena habituation for 20 minutes on Day 1. On Day2, they were presented with two cylinders containing a piece of cotton with a social odor for 20 minutes (social odor habituation). The social odor was collected by wiping the entire body of a novel stimulus rat with a piece of cotton, avoiding excrement. On the test day, the subjects were exposed to two cylinders containing pieces of cotton with different types of social odors: one from a neutral rat (injected with SAL) and the other from a distressed rat (injected with LiCl 150 mg/kg). The subjects were allowed to explore the apparatus freely for 10 minutes, and the time spent sniffing each cylinder was measured.

*SAP test (moderate distressed condition).* Basic procedures were the same as the SAP test procedure in the main contents. On the test day, the subjects were exposed to two cylinders containing conspecifics in different affective states: one was in a neutral state (injected with saline (SAL)), and the other was in a state of moderate distress (injected with a Lithium Chloride (LiCl) solution at 65 mg/kg). The subjects were allowed to explore the apparatus freely for 10 min, and sniffing behavior toward each cylinder was measured.

*SAP test (Stressed condition).* Basic procedures were the same as the SAP test procedure in the main contents. On the test day, the subjects were exposed to two cylinders containing conspecifics in different affective states: one was in a neutral state (no treatment), and the other was in a stressed state induced by 1 hour of restraint immediately before the test (Refer to Figure S4). The subjects were allowed to explore the apparatus freely for 10 min, and sniffing behavior toward each cylinder was measured.

**Statistics.**

Statistical analyses were performed using R (ver. 4.2.2). Changes in body weight during the neonatal treatment were analyzed using a two-way analysis of variance (ANOVA) with group and day after birth as factors, followed by multiple comparisons with Bonferroni correction.

The behavioral measurements in the open-field and elevated plus-maze tests were analyzed using a one-way ANOVA with group as the factor, followed by pairwise t-tests with Bonferroni correction.

In the social novelty preference test, a one-way ANOVA with demonstrator (novel or familiar) as the factor, followed by Bonferroni post-hoc tests, was conducted. To compare the novelty preference index to the chance level, one-sample t-tests were used.

For the behavioral analysis of the SAP (moderate-distressed condition and stressed condition) and Odor preference tests, the time spent sniffing behavior was analyzed using a two-way ANOVA with group and affective state of the demonstrator (neutral or distressed/stressed) as factors, followed by Bonferroni post-hoc tests. One-sample t-tests were used to compare the preference index for each group to the chance level. Additionally, following a one-way ANOVA, pairwise t-tests were performed to examine between-group differences in the SAP/Odor preference tests.

**Supplemental Figures**

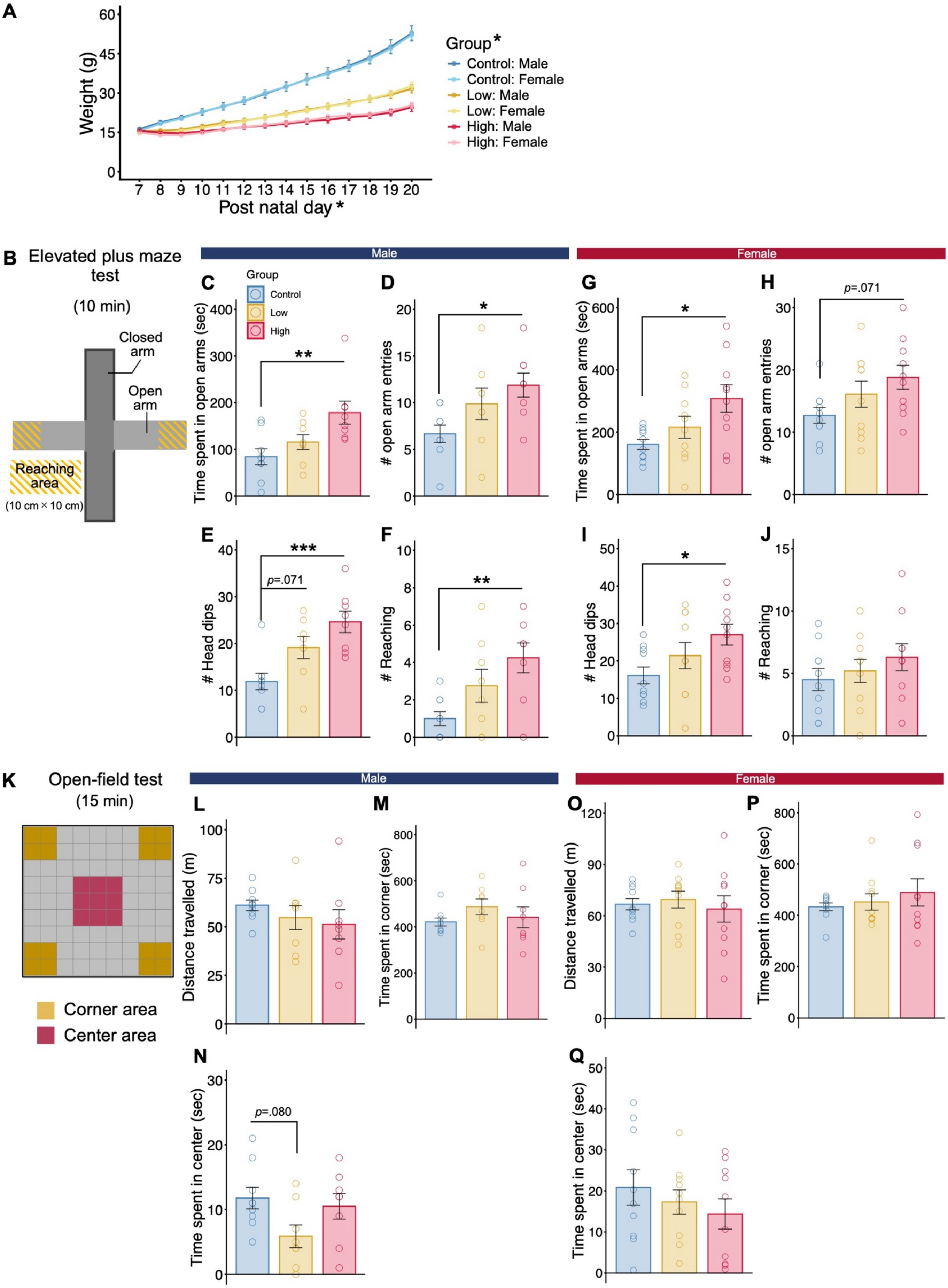

**Figure S1. Assessment of general behaviors in the neonatal MK-801-treated rats. A−Q** Data obtained from a total of 55 subjects (Control: male n = 9, female n = 10; Low: male n = 8, female n = 10; High: male n = 8, female n = 10). **A** Changes in body weight during neonatal MK-801 treatment. **B** Schematic illustration of the EPM test arena. **C−F** Behavioral measurements during the EPM test in males (**C** time spent in open arms; **D** the number of open arm entries; **E** the number of head dips; **F** the number of reaching area entries). **G−J** Behavioral measurements during the EPM test in females (**G** time spent in open arms; **H** the number of open arm entries; **I** the number of head dips; **J** the number of reaching area entries). **K** Schematic illustration of the open-field test arena. **L−N** Behavioral measurements during the open-field test in males (**L** distance travelled; **M** time spent in corner; **N** time spent in center). **O−Q** Behavioral measurements during the open-field test in females (**O** distance travelled; **P** time spent in corner; **Q** time spent in center). Bar and line graphs show the mean ± s.e.m. *$p < 0.05$, **$p < 0.01$, and ***$p < 0.001$ indicate significant differences between groups.

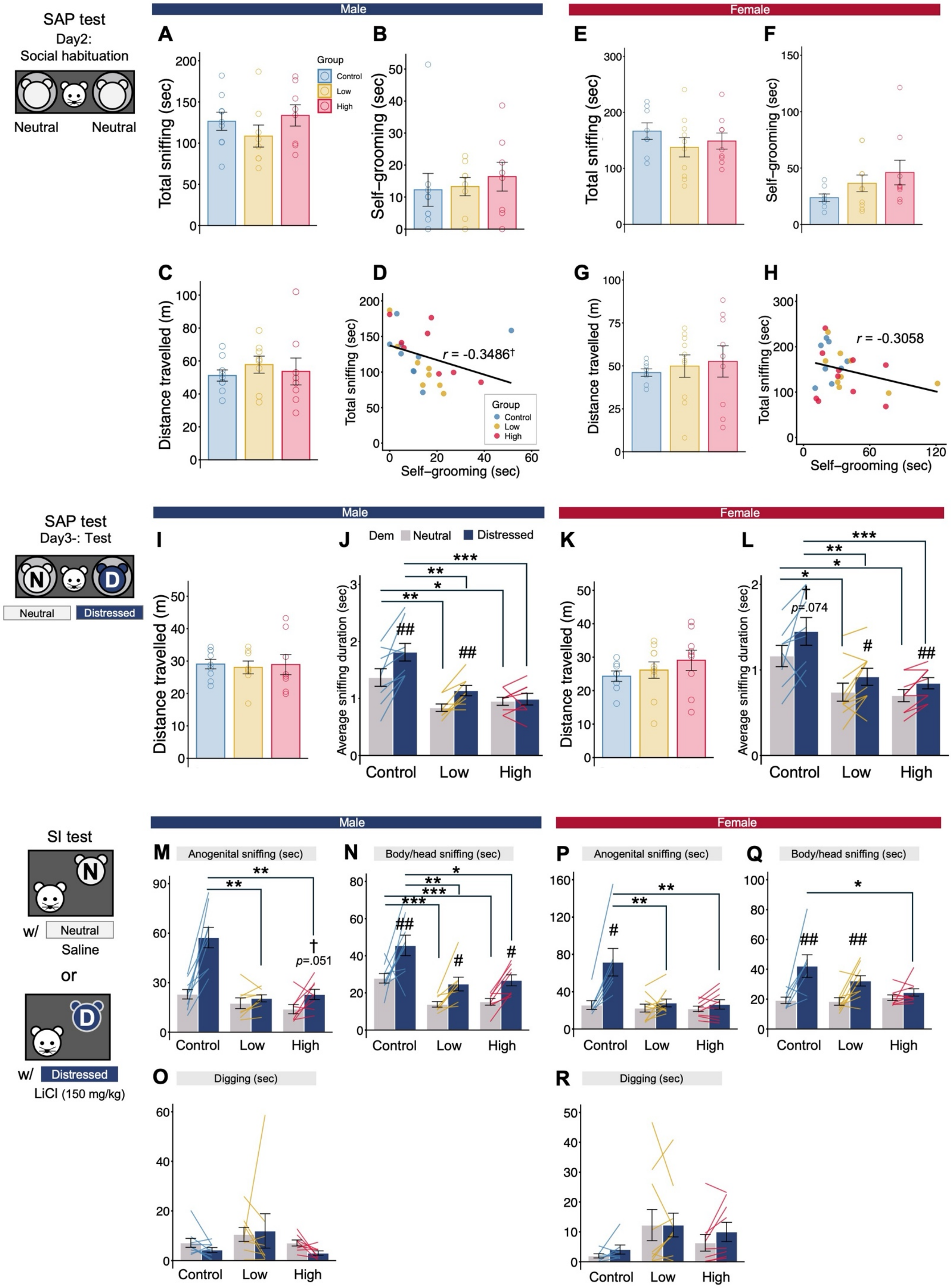

SAP test
Day2:
Social habituation
Neutral
Neutral
Male
Female
A
B
C
D
E
F
G
H
Total sniffing (sec)
Self−grooming (sec)
Distance travelled (m)
Group
Control
Low
High
r = -0.3486†
r = -0.3058
SAP test
Day3-: Test
N
D
Neutral
Distressed
I
J
K
L
Dem
Neutral
Distressed
Average sniffing duration (sec)
p=.074
Control
Low
High
SI test
w/
Neutral
Saline
or
w/
Distressed
LiCl (150 mg/kg)
M
N
O
P
Q
R
Anogenital sniffing (sec)
Body/head sniffing (sec)
Digging (sec)
p=.051

**Figure S2. Additional behavioral data during the SAP and SI tests.**

**A–L**, Data were obtained from 55 subjects (Control: male n = 9, female n = 8; Low: male n = 8, female n = 10; High: male n = 8, female n = 9). **A–D**, Social habituation measurements in males: time spent sniffing cylinders containing demonstrators (**A**), time spent self-grooming (**B**), distance travelled (**C**), and correlation between sniffing and self-grooming (**D**). **E–H**, Social habituation measurements in females: time spent sniffing cylinders containing demonstrators (**E**), time spent self-grooming (**F**), distance travelled (**G**), and correlation between sniffing and self-grooming (**H**). **I–L**, Additional SAP test measurements in males (**I–J**) and females (**K–L**): distance travelled (**I**, **K**) and average duration of a single sniffing event toward each cylinder (**J**, **L**). **M–R**, Additional SI test measurements were obtained from 51 subjects (Control: male n = 9, female n = 7; Low: male n = 8, female n = 10; High: male n = 8, female n = 9) in males (**M–O**) and females (**P–R**): time spent sniffing the anogenital area (**M**, **P**), time spent sniffing the head/body area (**N**, **Q**), and digging behavior (**O**, **R**). Bar and line graphs show the mean ± s.e.m. *$p < 0.05$, **$p < 0.01$, and ***$p < 0.001$ indicate significant differences between groups. #$p < 0.05$, ##$p < 0.01$, ###$p < 0.001$, and †$p < 0.10$ indicate significant differences or trends between neutral and distressed demonstrators.

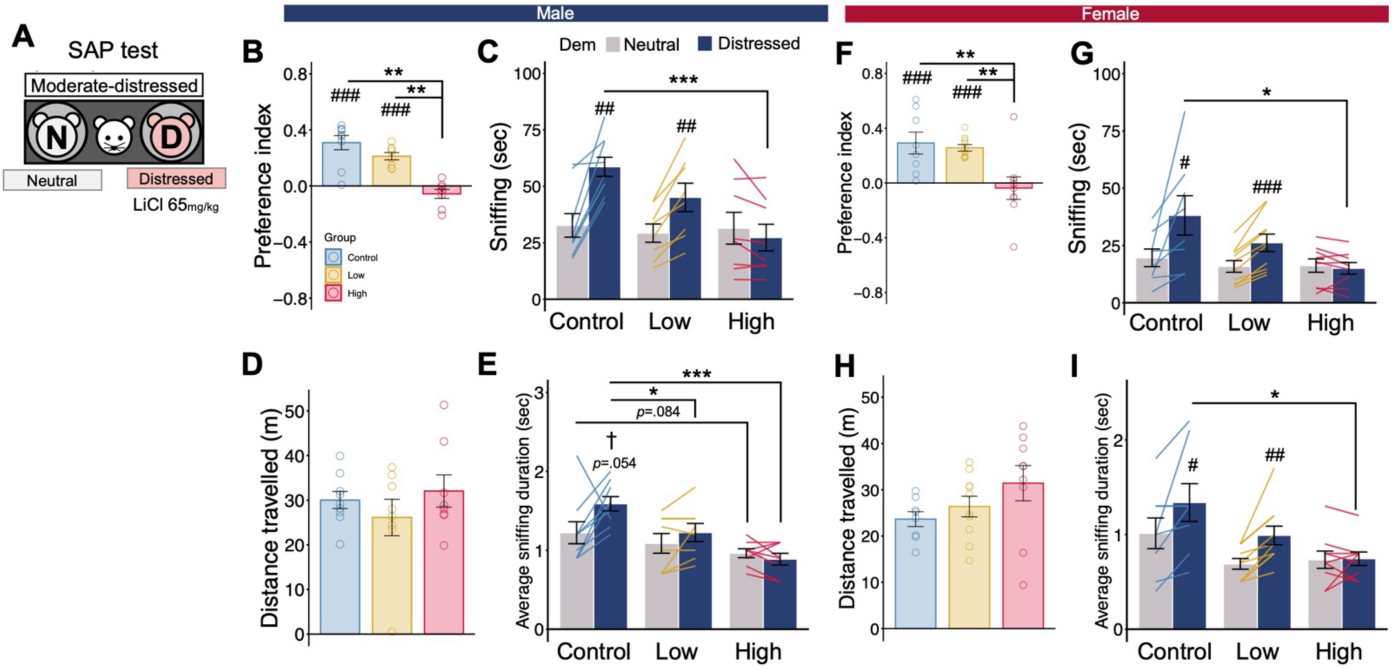

Male
Female
A
SAP test
Moderate-distressed
N
D
Neutral
Distressed
LiCl 65mg/kg
B
Preference index
Group
Control
Low
High
C
Dem
Neutral
Distressed
Sniffing (sec)
Control
Low
High
D
Distance travelled (m)
E
Average sniffing duration (sec)
p=.054
p=.084
F
Preference index
G
Sniffing (sec)
H
Distance travelled (m)
I
Average sniffing duration (sec)

**Figure S3. Social affective responses toward a moderately distressed demonstrator were also impaired in MK-801-treated rats.**

**A** Schematic illustration of the social affective preference (SAP) test using a moderately distressed demonstrator induced by a low-dose LiCl administration. **B–I**, Data were obtained from 52 subjects (Control: male n = 9, female n = 8; Low: male n = 8, female n = 10; High: male n = 8, female n = 9). **B–E** Behavioral measurements during the SAP test in males: preference index (**B**), time spent sniffing each demonstrator (**C**), distance travelled (**D**), and mean duration of individual sniffing events toward each cylinder (**E**). **F–I** Behavioral measurements during the SAP test in females: preference index (**F**), time spent sniffing each demonstrator (**G**), distance travelled (**H**), and mean duration of individual sniffing events toward each cylinder (**I**). Bar and line graphs show the mean ± s.e.m. *$p < 0.05$, **$p < 0.01$, and ***$p < 0.001$ indicate significant differences between groups. #$p < 0.05$, ##$p < 0.01$, ###$p < 0.001$, and †$p < 0.10$ indicate significant differences or trends between neutral and distressed demonstrators in panels **C**, **E**, **G**, **I**, or significant differences or trends from chance level in the preference index in panels **B** and **F**.

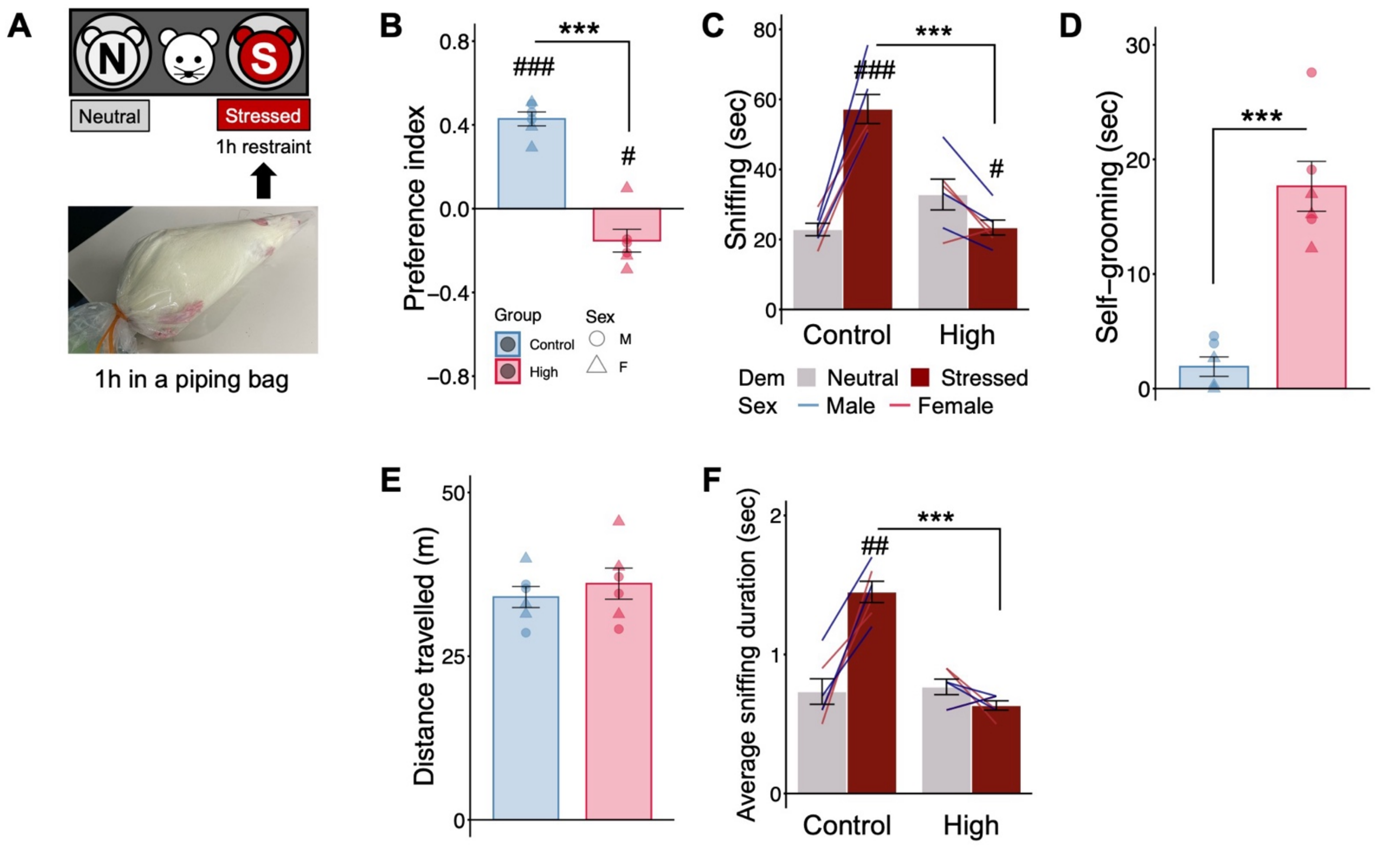
A
Neutral
Stressed
1h restraint
1h in a piping bag
B
***
###
#
Preference index
Group
Control
High
Sex
M
F
C
***
###
#
Sniffing (sec)
Control
High
Dem Neutral Stressed
Sex Male Female
D
***
Self−grooming (sec)
E
Distance travelled (m)
F
***
##
Average sniffing duration (sec)
Control
High

**Figure S4. Behavioral responses toward a restraint-stressed demonstrator were also disturbed in MK-801-treated rats.**

**A** Schematic illustration of the social affective preference (SAP) test using a demonstrator exposed to 1-h restraint stress. **B–F** Behavioral measures during the test were obtained from 12 subjects (Control: male n = 3, female n = 3; High: male n = 3, female n = 3): preference index (**B**), time spent sniffing each demonstrator (**C**), time spent self-grooming (**D**), distance traveled (**E**), and mean duration of individual sniffing events toward each cylinder (**F**). Bar and line graphs show the mean ± s.e.m. ***$p < 0.001$ indicates a significant difference between groups. #$p < 0.05$, ##$p < 0.01$, and ###$p < 0.001$ indicate significant differences between neutral and stressed demonstrators in panels **C** and **F**, or from chance level in the preference index in panel **B**.

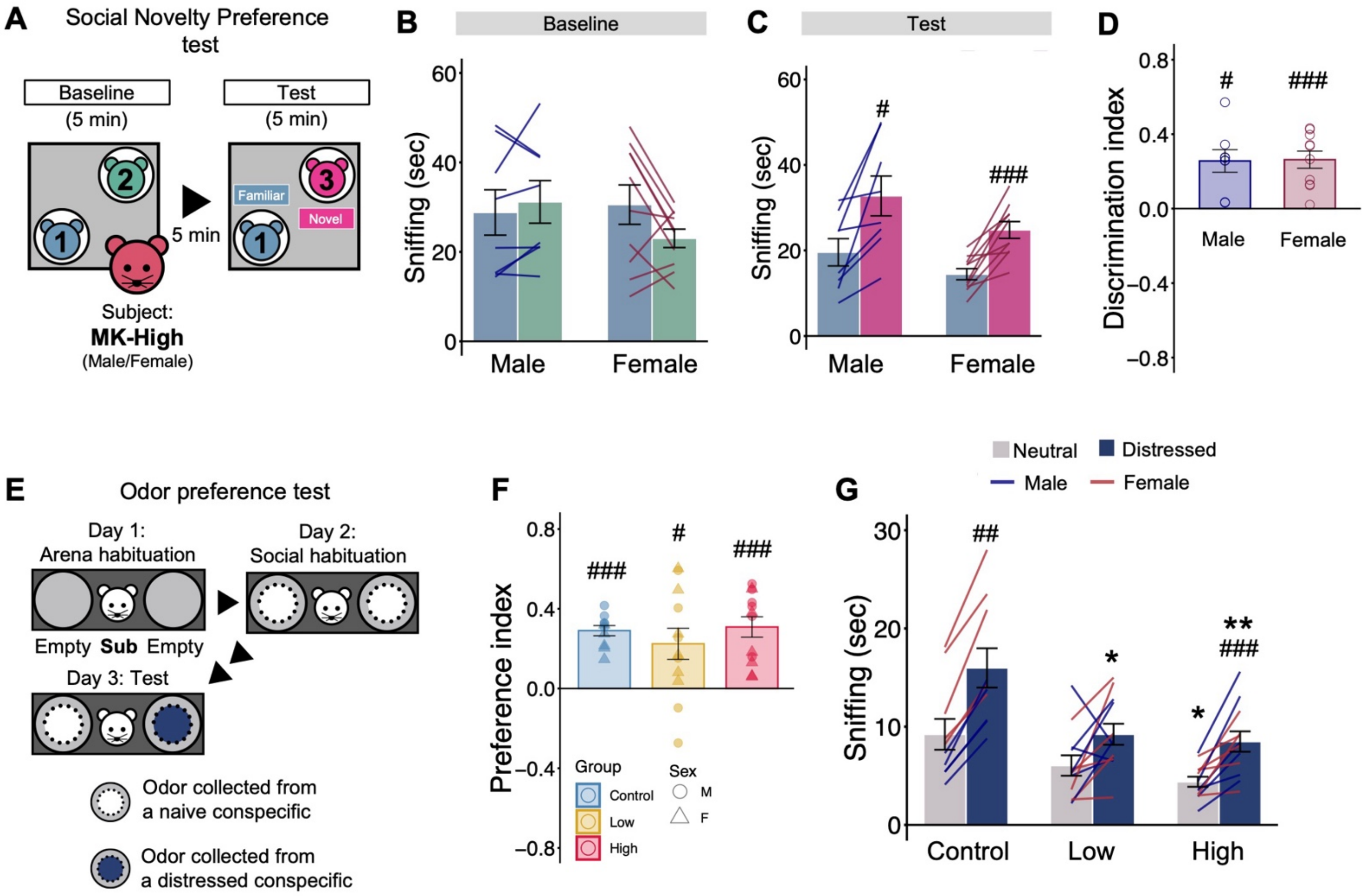

A
Social Novelty Preference test
Baseline (5 min)
Test (5 min)
Familiar
Novel
5 min
Subject:
MK-High
(Male/Female)
B
Baseline
Sniffing (sec)
Male
Female
C
Test
#
###
D
Discrimination index
E
Odor preference test
Day 1: Arena habituation
Day 2: Social habituation
Empty Sub Empty
Day 3: Test
Odor collected from a naive conspecific
Odor collected from a distressed conspecific
F
Preference index
Group
Control
Low
High
Sex
M
F
G
Neutral
Distressed
Male
Female
##
*
**
Control
Low
High

**Figure S5. Social novelty preference and odor preference were intact in MK-801-treated rats.**

**A** Schematic illustration of the social novelty preference test. **B–D** Behavioral measures during the test were obtained from 18 subjects (High: male n = 8, female n = 10): time spent sniffing each demonstrator during the baseline phase (**B**) and test phase (**C**), and discrimination index (**D**). **E** Schematic illustration of the odor preference test. **F** and **G** Behavioral measures during the test were obtained from 33 subjects (Control: male n = 4, female n = 5; Low: male n = 6, female n = 6; High: male n = 6, female n = 6): preference index (**F**) and time spent sniffing each cylinder containing odor samples collected from a neutral or distressed demonstrator (**G**). Bar and line graphs show the mean ± s.e.m. $^{*}p < 0.05$ and $^{**}p < 0.01$ indicate significant differences between groups. $\#p < 0.05$, $\#\#p < 0.01$, and $\#\#\#p < 0.001$ indicate significant differences between familiar and novel demonstrators in panel **C**, between neutral and distressed odors in panel **G**, or from chance level in the index values in panels **D** and **F**.

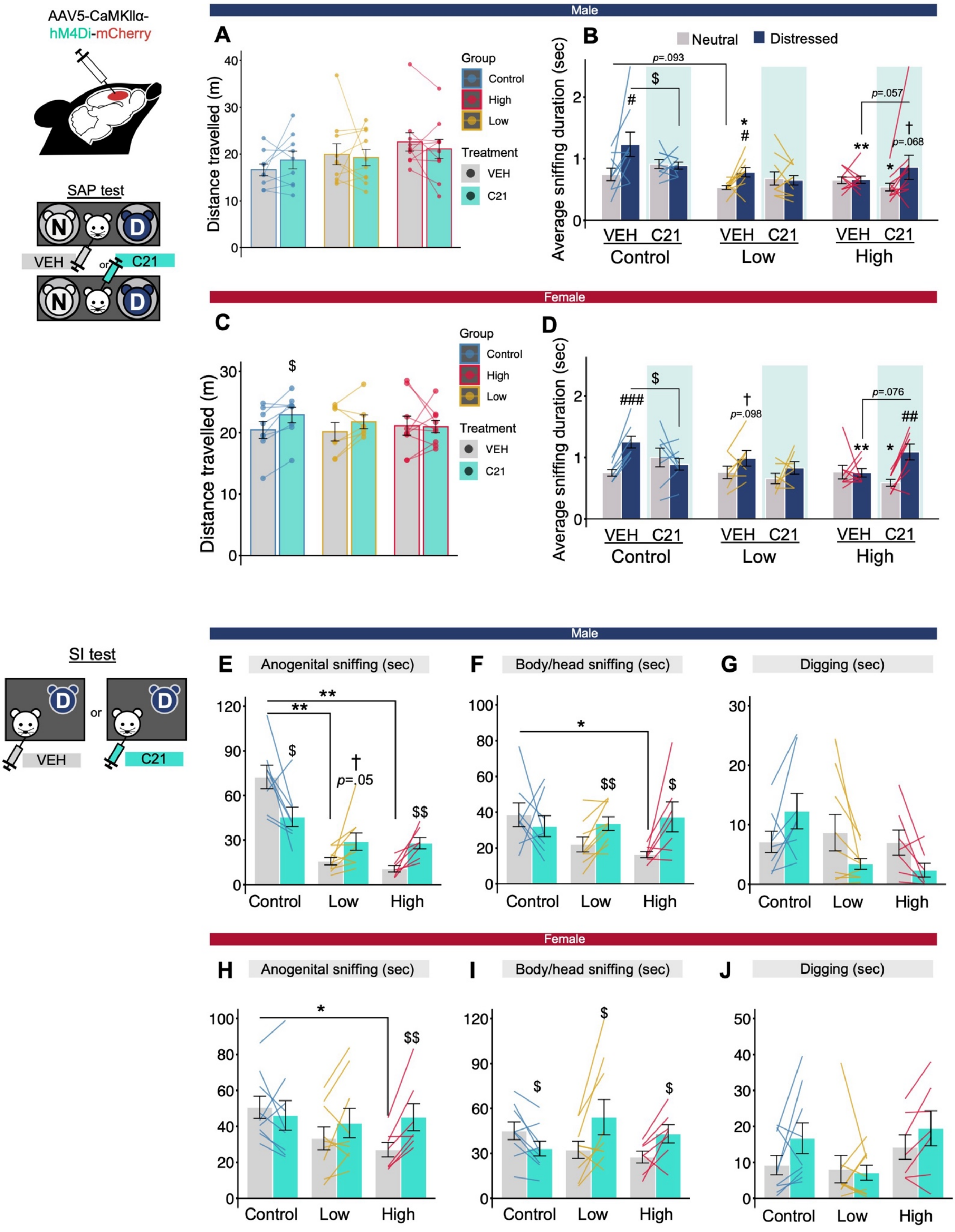

AAV5-CaMKIIα-
hM4Di-mCherry
SAP test
N
D
VEH
or
C21
Male
A
Distance travelled (m)
Group
Control
High
Low
Treatment
VEH
C21
B
Neutral
Distressed
Average sniffing duration (sec)
p=.093
$
#
*
#
p=.057
**
*
†
p=.068
VEH C21
Control
Low
High
Female
C
$
D
###
†
p=.098
p=.076
##
SI test
or
E
Anogenital sniffing (sec)
**
**
†
p=.05
$$
F
Body/head sniffing (sec)
*
G
Digging (sec)
Control
Low
High
H
I
J

**Figure S6. DREADD-mediated inhibition of mPFC pyramidal cells did not alter locomotor activity or general exploratory behavior but modulated social investigation.**

**A–D** Data were obtained from 53 subjects (Control: male n = 9, female n = 8; Low: male n = 10, female n = 7; High: male n = 10, female n = 9). **A** and **B,** Additional behavioral measures during the SAP test in males: distance traveled (**A**) and mean duration of individual sniffing events directed toward each demonstrator (**B**). **C** and **D**, Additional behavioral measures during the SAP test in females: distance traveled (**C**) and mean duration of individual sniffing events directed toward each demonstrator (**D**). **E–J,** Data were obtained from 50 subjects (Control: male n = 9, female n = 9; Low: male n = 9, female n = 9; High: male n = 7, female n = 7). **E–G,** Additional behavioral measures during the SI test in males: time spent sniffing the anogenital area (**E**), sniffing the body/head area (**F**), and digging (**G**). **H–J,** Additional behavioral measures during the SI test in females: time spent sniffing the anogenital area (**H**), sniffing the body/head area (**I**), and digging (**J**). Bar and line graphs show the mean ± s.e.m. *$p < 0.05$, **$p < 0.01$, and ***$p < 0.001$ indicate significant differences between groups. #$p < 0.05$, ##$p < 0.01$, ###$p < 0.001$, and †$p < 0.10$ indicate significant differences or trends between neutral and distressed demonstrators. \$$p < 0.05$ and \$\$$p < 0.01$ indicate significant differences between vehicle and C21 conditions.

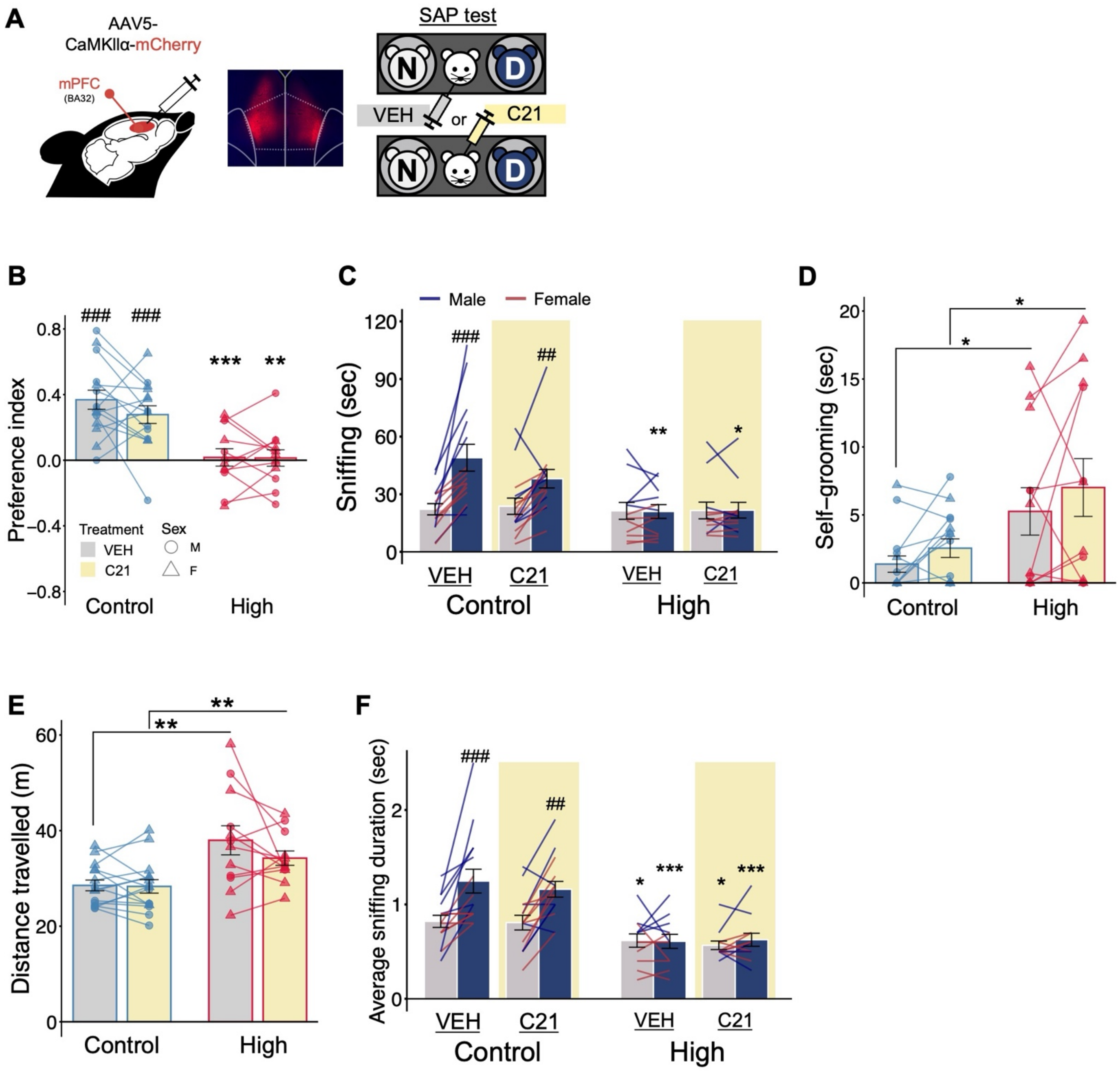
A
AAV5-
CaMKIIα-mCherry
mPFC
(BA32)
SAP test
N
D
VEH
or
C21
B
###
###
***
**
Preference index
Treatment
VEH
C21
Sex
M
F
Control
High
C
Male
Female
Sniffing (sec)
###
##
**
*
VEH
C21
VEH
C21
Control
High
D
Self−grooming (sec)
*
*
Control
High
E
**
**
Distance travelled (m)
Control
High
F
Average sniffing duration (sec)
###
##
*
***
*
***
VEH
C21
VEH
C21
Control
High

**Figure S7. Control viral expression in the mPFC does not affect social affective behavior.**

**A−F** Data derived from a total of 27 subjects (Control: male n = 7, female n = 8, High: male n = 5, female n = 7). **A** Schematic of viral vector injection. Control DREADD targeting mPFC pyramidal cells was introduced into the mPFC. **B–F** Results obtained from 27 subjects: preference index (**B**), time spent sniffing each demonstrator (**C**), self-grooming during the test (**D**), distance traveled (**E**), and mean duration of individual sniffing events directed toward each demonstrator (**F**).

Bar and line graphs show the mean ± s.e.m. *$p < 0.05$, **$p < 0.01$, and ***$p < 0.001$ indicate significant differences between groups. #$p < 0.05$, ##$p < 0.01$, ###$p < 0.001$, and †$p < 0.10$ indicate significant differences or trends between neutral and distressed demonstrators in panels **C**, **F**, or from chance level in panel **B**.

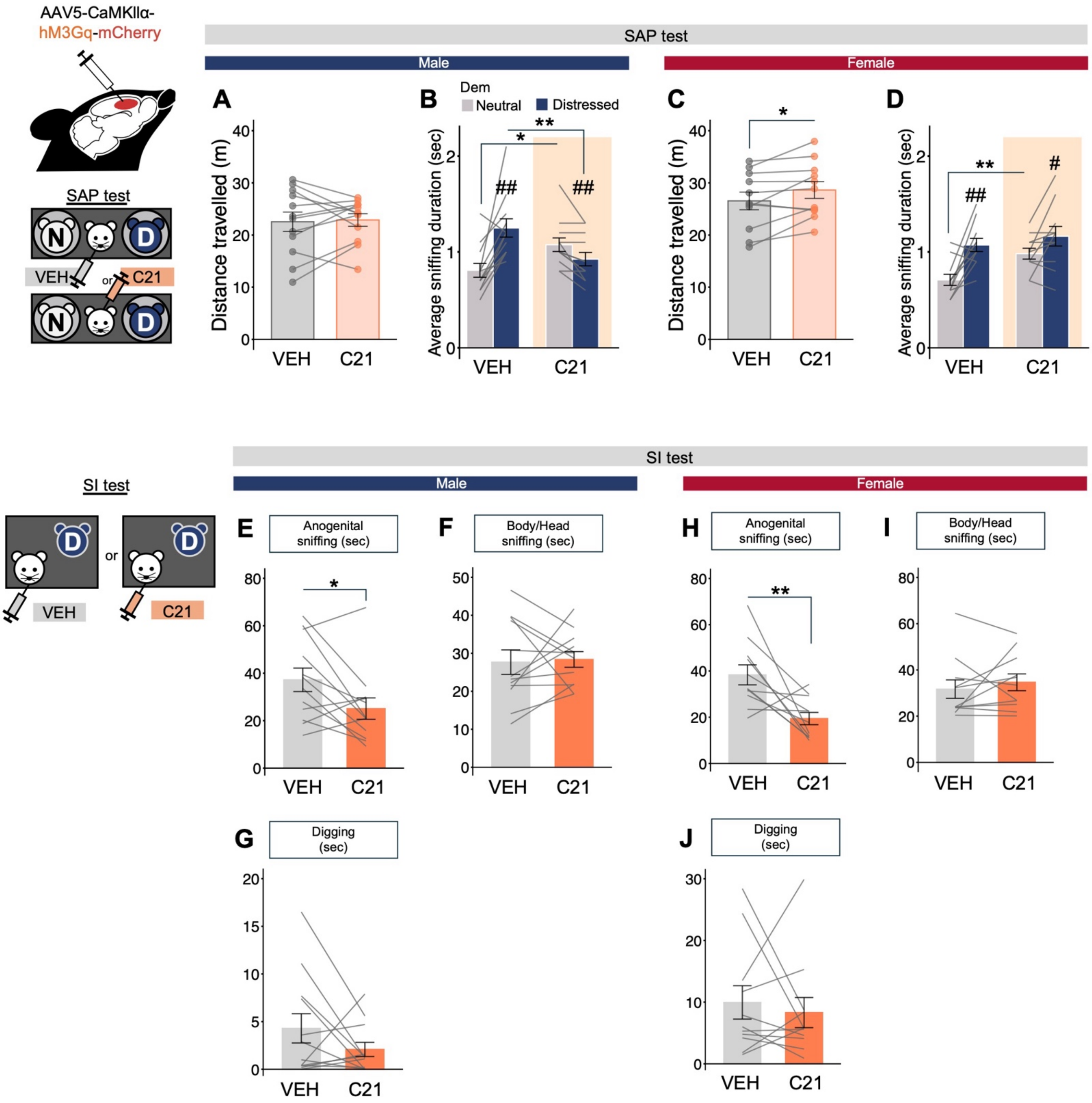

AAV5-CaMKIIα-
hM3Gq-mCherry
SAP test
N
D
VEH
or
C21
SAP test
Male
Female
A
B
C
D
Dem
Neutral
Distressed
Distance travelled (m)
Average sniffing duration (sec)
VEH
C21
*
**
##
#
SI test
Male
Female
E
F
G
H
I
J
Anogenital sniffing (sec)
Body/Head sniffing (sec)
Digging (sec)

**Figure S8. Activation of mPFC pyramidal cells replicated the social exploratory style of MK-801-treated rats.**

**A−J** Data derived from a total of 21 subjects (Male: n = 12, Female: n = 11). **A** and **B**, Additional behavioral measures during the SAP test in males: distance traveled (**A**) and mean duration of individual sniffing events directed toward each demonstrator (**B**). **C** and **D**, Additional behavioral measures during the SAP test in females: distance traveled (**C**) and mean duration of individual sniffing events directed toward each demonstrator (**D**). **E–G** Additional behavioral measures during the SI test in males: time spent sniffing the anogenital area (**E**), sniffing the body/head area (**F**), and digging (**G**). **H–J** Additional behavioral measures during the SI test in females: time spent sniffing the anogenital area (**H**), sniffing the body/head area (**I**), and digging (**J**). Bar and line graphs show the mean ± s.e.m. *$p < 0.05$ and **$p < 0.01$ indicate significant differences between vehicle- and C21-treated conditions. #$p < 0.05$, ##$p < 0.01$, and ###$p < 0.001$ indicate significant differences between neutral and distressed demonstrators.

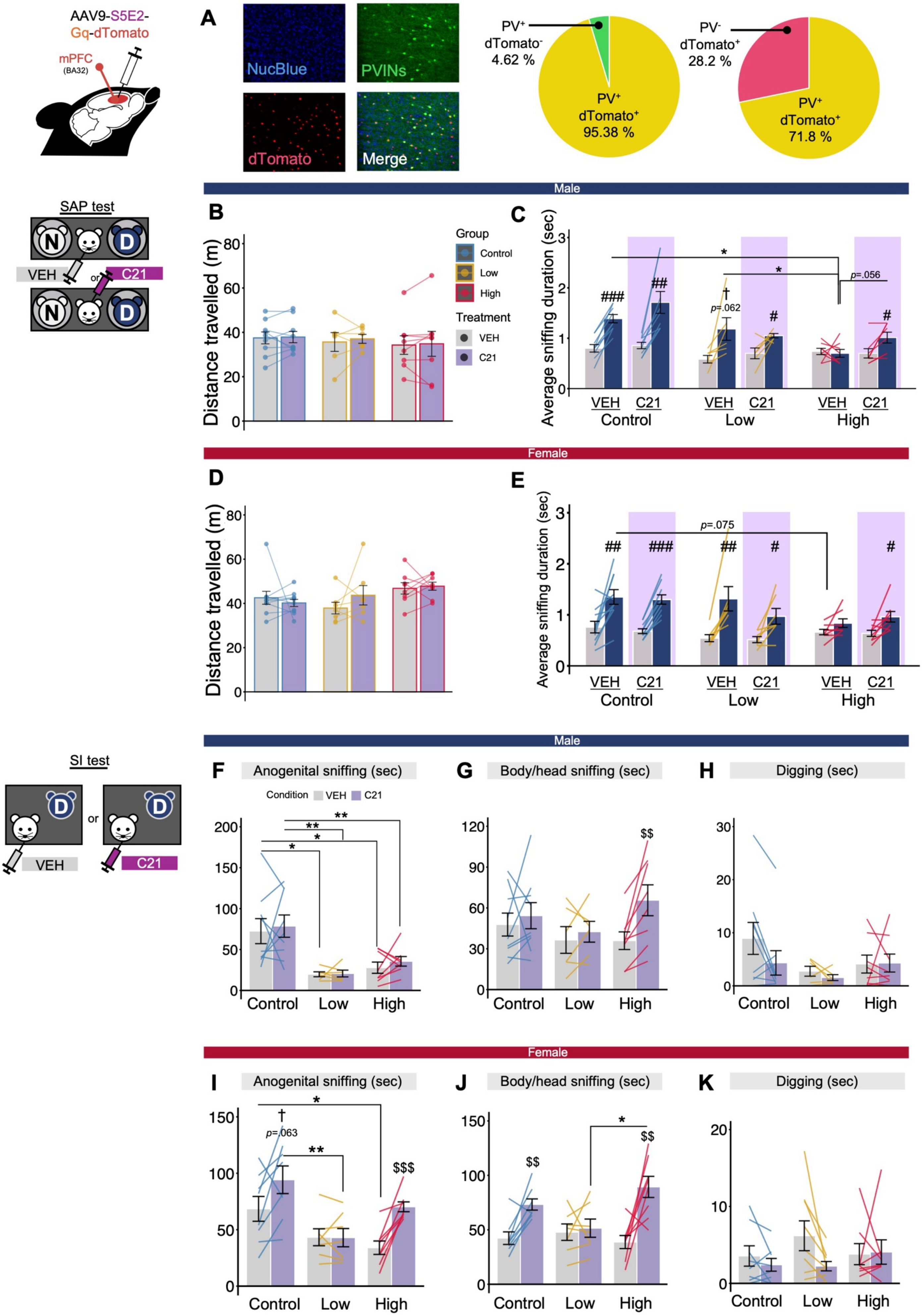
AAV9-S5E2-
Gq-dTomato
mPFC
(BA32)
A
NucBlue
PVINs
dTomato
Merge
PV+ dTomato- 4.62 %
PV+ dTomato+ 95.38 %
PV- dTomato+ 28.2 %
PV+ dTomato+ 71.8 %
Male
SAP test
N
D
VEH
or
C21
B
Distance travelled (m)
Group
Control
Low
High
Treatment
VEH
C21
C
Average sniffing duration (sec)
###
##
†
p=.062
#
p=.056
VEH C21
Control
Low
High
Female
D
E
p=.075
Male
SI test
F
Anogenital sniffing (sec)
Condition VEH C21
G
Body/head sniffing (sec)
$$
H
Digging (sec)
Female
I
p=.063
$$$
J
K

**Figure S9. Chemogenetic activation of mPFC PVINs did not affect locomotor activity or general exploratory behavior but altered social investigation in the Control and MK-High groups.**

**A,** Assessment of S5E2 promoter-driven viral targeting of PVINs in the mPFC. Representative merged images show the overlap between S5E2-Gq-mCherry expression and dTomato-labeled PVINs. Pie charts show the specificity of viral expression among S5E2-Gq-mCherry-positive cells and the infection efficiency among dTomato-positive PVINs. **B–E** Data were obtained from 48 subjects (Control: male n = 9, female n = 10; Low: male n = 6, female n = 7; High: male n =8, female n = 8). **B** and **C**, Additional behavioral measures during the SAP test in males: distance traveled (**B**) and mean duration of individual sniffing events directed toward each demonstrator (**C**). **D** and **E**, Additional behavioral measures during the SAP test in females: distance traveled (**D**) and mean duration of individual sniffing events directed toward each demonstrator (**E**). **F–K** Data were obtained from 46 subjects (Control: male n = 9, female n = 8; Low: male n = 6, female n = 7; High: male n = 8, female n = 8). **F–H** Additional behavioral measures during the SI test in males: time spent sniffing the anogenital area (**F**), sniffing the body/head area (**G**), and digging (**H**). **I–K** Additional behavioral measures during the SI test in females: time spent sniffing the anogenital area (**I**), sniffing the body/head area (**J**), and digging (**K**). Bar and line graphs show the mean ± s.e.m. *$p < 0.05$, **$p < 0.01$, and ***$p < 0.001$ indicate significant differences between groups. #$p < 0.05$, ##$p < 0.01$, ###$p < 0.001$, and †$p < 0.10$ indicate significant differences or trends between neutral and distressed demonstrators. \$$p < 0.05$ and \$\$$p < 0.01$ indicate significant differences between vehicle and C21 conditions.
