## Supplemental Material 2 for "Neonatal NMDA receptor blockade disrupts prefrontal cortical regulation of social affective behavior: implications for schizophrenia"

### Supplemental tables

| Figure | Analysis | Factor | Statistical value | p-value | |
|---|---|---|---|---|---|
| 1E | One-sample t-test | Control vs chance level | $t_{8} = 6.308$ | $p < .001$ | *** |
| | | Low vs chance level | $t_{7} = 8.511$ | $p < .001$ | *** |
| | | High vs chance level | $t_{7} = 1.415$ | $p = .2$ | n.s. |
| | Pairwise t-test | Control vs Low | | $p = 1$ | n.s. |
| | | Control vs High | | $p < .01$ | ** |
| | | Low vs High | | $p < .01$ | ** |
| 1F | Two-way ANOVA | Group | $F_{2,22} = 20.399$ | $p < .001$ | *** |
| | | Demonstrator | $F_{1,22} = 66.367$ | $p < .001$ | *** |
| | | Group: Demonstrator | $F_{2,22} = 16.215$ | $p < .001$ | *** |
| | Pairwise t-test | Control: Neutral vs Distressed | | $p < .001$ | *** |
| | (Group) | Low: Neutral vs Distressed | | $p < .001$ | *** |
| | | High: Neutral vs Distressed | | $p = .721$ | n.s. |
| | (Demonstrator) | Neutral: Control vs Low | | $p < .01$ | ** |
| | | Neutral: Control vs High | | $p < .05$ | * |
| | | Neutral: Low vs High | | $p = .691$ | n.s. |
| | | Distressed: Control vs Low | | $p < .01$ | ** |
| | | Distressed: Control vs High | | $p < .001$ | *** |
| | | Distressed: Low vs High | | $p < .05$ | * |
| 1G | One-way ANOVA | Group | $F_{2,22} = 6.446$ | $p < .01$ | ** |
| | Pairwise t-test | Control vs Low | | $p < .05$ | * |
| | | Control vs High | | $p < .05$ | * |
| | | Low vs High | | $p = 1$ | n.s. |
| 1H | Pearson's product-moment | | $t_{23} = -3.438$ | $p < .01$ | ** |
| | correlation coefficient | | $r = -0.583$ | | |
| 1I | One-sample t-test | Control vs chance level | $t_{7} = 3.9652$ | $p < .01$ | ** |
| | | Low vs chance level | $t_{9} = 2.900$ | $p < .05$ | * |
| | | High vs chance level | $t_{8} = -0.539$ | $p = .605$ | n.s. |
| | Pairwise t-test | Control vs Low | | $p = .688$ | n.s. |
| | | Control vs High | | $p < .01$ | ** |
| | | Low vs High | | $p = .067$ | † |
| 1J | Two-way ANOVA | Group | $F_{2,24} = 1.629$ | $p = .217$ | n.s. |
| | | Demonstrator | $F_{1,24} = 17.482$ | $p < .001$ | *** |
| | | Group: Demonstrator | $F_{2,24} = 5.425$ | $p < .05$ | * |
| | Pairwise t-test | Control: Neutral vs Distressed | | $p < .01$ | ** |
| | (Group) | Low: Neutral vs Distressed | | $p = .09$ | † |
| | | High: Neutral vs Distressed | | $p = .695$ | n.s. |
| | (Demonstrator) | Neutral: Control vs Low | | $p = 1$ | n.s. |

| | | | | | | |
|---|---|---|---|---|---|---|
| | | Neutral: Control vs High | | | $p = 1$ | n.s. |
| | | Neutral: Low vs High | | | $p = 1$ | n.s. |
| | | Distressed: Control vs Low | | | $p = .217$ | n.s. |
| | | Distressed: Control vs High | | | $p = .074$ | † |
| | | Distressed: Low vs High | | | $p = 1$ | n.s. |
| 1K | One-way ANOVA | Group | $F_{2,24}$ | $= 4.825$ | $p < .05$ | * |
| | Pairwise t-test | Control vs Low | | | $p < .05$ | * |
| | | Control vs High | | | $p = .166$ | n.s. |
| | | Low vs High | | | $p = .913$ | n.s. |
| 1L | Pearson's product-moment | | $t_{25}$ | $= -3.438$ | $p = .189$ | n.s. |
| | correlation coefficient | | $r$ | $= -0.261$ | | |
| 1N | Type III Likelihood Ratio | Group | $X^2_{2}$ | $= 3221.9$ | $p < .001$ | *** |
| | Chi-Square Test | Demonstrator | $X^2_{1}$ | $= 2018.0$ | $p < .001$ | *** |
| | | Group: Demonstrator | $X^2_{2}$ | $= 187.5$ | $p < .001$ | *** |
| | Tukey-adjusted EMM | Control: Neutral vs Distressed | | | $p < .001$ | *** |
| | comparisons (Group) | Low: Neutral vs Distressed | | | $p < .001$ | *** |
| | | High: Neutral vs Distressed | | | $p < .001$ | *** |
| | (Demonstrator) | Neutral: Control vs Low | | | $p < .001$ | *** |
| | | Neutral: Control vs High | | | $p < .001$ | *** |
| | | Neutral: Low vs High | | | $p = .093$ | † |
| | | Distressed: Control vs Low | | | $p < .001$ | *** |
| | | Distressed: Control vs High | | | $p < .001$ | *** |
| | | Distressed: Low vs High | | | $p < .001$ | *** |
| 1O | Two-way ANOVA | Group | $F_{2,22}$ | $= 23.032$ | $p < .001$ | *** |
| | | Demonstrator | $F_{1,22}$ | $= 45.774$ | $p < .001$ | *** |
| | | Group: Demonstrator | $F_{2,22}$ | $= 7.906$ | $p < .01$ | ** |
| | Pairwise t-test | Control: Neutral vs Distressed | | | $p < .001$ | *** |
| | (Group) | Low: Neutral vs Distressed | | | $p = .059$ | † |
| | | High: Neutral vs Distressed | | | $p < .05$ | * |
| | (Demonstrator) | Neutral: Control vs Low | | | $p < .01$ | ** |
| | | Neutral: Control vs High | | | $p < .01$ | ** |
| | | Neutral: Low vs High | | | $p = 1$ | n.s. |
| | | Distressed: Control vs Low | | | $p < .001$ | *** |
| | | Distressed: Control vs High | | | $p < .001$ | *** |
| | | Distressed: Low vs High | | | $p = 1$ | n.s. |
| 1P | Two-way ANOVA | Group | $F_{2,22}$ | $= 31.712$ | $p < .001$ | *** |
| | | Demonstrator | $F_{1,22}$ | $= 13.528$ | $p < .01$ | ** |
| | | Group: Demonstrator | $F_{2,22}$ | $= 14.586$ | $p < .001$ | *** |
| | Pairwise t-test | Control: Neutral vs Distressed | | | $p < .001$ | *** |
| | (Group) | Low: Neutral vs Distressed | | | $p = .492$ | n.s. |

| | | | | | |
|---|---|---|---|---|---|
| | | High: Neutral vs Distressed | | $p$ =.548 | n.s. |
| | (Demonstrator) | Neutral: Control vs Low | | $p$ <.001 | *** |
| | | Neutral: Control vs High | | $p$ <.001 | *** |
| | | Neutral: Low vs High | | $p$ =1 | n.s. |
| | | Distressed: Control vs Low | | $p$ <.05 | * |
| | | Distressed: Control vs High | | $p$ =.147 | n.s. |
| | | Distressed: Low vs High | | $p$ =1 | n.s. |
| 1Q | Two-way ANOVA | Group | $F_{2,22}$ = 8.956 | $p$ <.01 | ** |
| | | Demonstrator | $F_{1,22}$ = 0.125 | $p$ =.728 | n.s. |
| | | Group: Demonstrator | $F_{2,22}$ = 1.248 | $p$ =.307 | n.s. |
| | Pairwise t-test (Group) | Control: Neutral vs Distressed | | $p$ =.362 | n.s. |
| | | Low: Neutral vs Distressed | | $p$ =.477 | n.s. |
| | | High: Neutral vs Distressed | | $p$ =.254 | n.s. |
| | (Demonstrator) | Neutral: Control vs Low | | $p$ =.071 | † |
| | | Neutral: Control vs High | | $p$ <.05 | * |
| | | Neutral: Low vs High | | $p$ =1 | n.s. |
| | | Distressed: Control vs Low | | $p$ <.01 | ** |
| | | Distressed: Control vs High | | $p$ =.074 | † |
| | | Distressed: Low vs High | | $p$ =.622 | n.s. |
| 1R | Type III Likelihood Ratio Chi-Square Test | Group | $X^2$ 2 = 2678.33 | $p$ <.001 | *** |
| | | Demonstrator | $X^2$ 1 = 2485.95 | $p$ <.001 | *** |
| | | Group: Demonstrator | $X^2$ 2 = 814.05 | $p$ <.001 | *** |
| | Tukey-adjusted EMM comparisons (Group) | Control: Neutral vs Distressed | | $p$ <.001 | *** |
| | | Low: Neutral vs Distressed | | $p$ <.001 | *** |
| | | High: Neutral vs Distressed | | $p$ <.001 | *** |
| | (Demonstrator) | Neutral: Control vs Low | | $p$ <.001 | *** |
| | | Neutral: Control vs High | | $p$ <.05 | * |
| | | Neutral: Low vs High | | $p$ =.165 | n.s. |
| | | Distressed: Control vs Low | | $p$ <.001 | *** |
| | | Distressed: Control vs High | | $p$ <.001 | *** |
| | | Distressed: Low vs High | | $p$ <.001 | *** |
| 1S | Two-way ANOVA | Group | $F_{2,23}$ = 8.408 | $p$ <.01 | ** |
| | | Demonstrator | $F_{1,23}$ = 46.038 | $p$ <.001 | *** |
| | | Group: Demonstrator | $F_{2,23}$ = 13.981 | $p$ <.001 | *** |
| | Pairwise t-test (Group) | Control: Neutral vs Distressed | | $p$ <.01 | ** |
| | | Low: Neutral vs Distressed | | $p$ <.05 | * |
| | | High: Neutral vs Distressed | | $p$ =.06 | † |
| | (Demonstrator) | Neutral: Control vs Low | | $p$ =1 | n.s. |
| | | Neutral: Control vs High | | $p$ =1 | n.s. |
| | | Neutral: Low vs High | | $p$ =1 | n.s. |

| | | | | | |
|---|---|---|---|---|---|
| | | Distressed: Control vs Low | | $p < .01$ | ** |
| | | Distressed: Control vs High | | $p < .001$ | *** |
| | | Distressed: Low vs High | | $p = 1$ | n.s. |
| 1T | Two-way ANOVA | Group | $F_{2,23} = 10.980$ | $p < .001$ | *** |
| | | Demonstrator | $F_{1,23} = 10.461$ | $p < .01$ | ** |
| | | Group: Demonstrator | $F_{2,23} = 7.135$ | $p < .01$ | ** |
| | Pairwise t-test | Control: Neutral vs Distressed | | $p < .05$ | * |
| | (Group) | Low: Neutral vs Distressed | | $p = 1$ | n.s. |
| | | High: Neutral vs Distressed | | $p = .104$ | n.s. |
| | (Demonstrator) | Neutral: Control vs Low | | $p = .084$ | † |
| | | Neutral: Control vs High | | $p < .05$ | * |
| | | Neutral: Low vs High | | $p = 1$ | n.s. |
| | | Distressed: Control vs Low | | $p < .01$ | ** |
| | | Distressed: Control vs High | | $p < .01$ | ** |
| | | Distressed: Low vs High | | $p = 1$ | n.s. |
| 1U | Two-way ANOVA | Group | $F_{2,23} = 5.046$ | $p < .05$ | * |
| | | Demonstrator | $F_{1,23} = 2.311$ | $p = .142$ | n.s. |
| | | Group: Demonstrator | $F_{2,23} = 0.010$ | $p = .990$ | n.s. |
| | Pairwise t-test | Control: Neutral vs Distressed | | $p < .05$ | * |
| | (Group) | Low: Neutral vs Distressed | | $p = .458$ | n.s. |
| | | High: Neutral vs Distressed | | $p = .352$ | n.s. |
| | (Demonstrator) | Neutral: Control vs Low | | $p < .05$ | * |
| | | Neutral: Control vs High | | $p = .183$ | n.s. |
| | | Neutral: Low vs High | | $p = .926$ | n.s. |
| | | Distressed: Control vs Low | | $p = .17$ | n.s. |
| | | Distressed: Control vs High | | $p = .706$ | n.s. |
| | | Distressed: Low vs High | | $p = 1$ | n.s. |

**Table S1. Detailed statistical results corresponding to Figure 1.**

| Figure | Analysis | Factor | Statistical value | p-value |
|---|---|---|---|---|
| 2D | One-sample t-test | Control (Vehicle) vs chance level | $t_{8} = 3.533$ | $p < .01$ ** |
| | | Control (C21) vs chance level | $t_{8} = 1.094$ | $p = .306$ n.s. |
| | | Low (Vehicle) vs chance level | $t_{9} = 3.662$ | $p < .01$ ** |
| | | Low(C21) vs chance level | $t_{9} = -0.001$ | $p = .999$ n.s. |
| | | High (Vehicle) vs chance level | $t_{9} = 0.215$ | $p = .836$ n.s. |
| | | High (C21) vs chance level | $t_{9} = 2.253$ | $p = .051$ † |
| | Two-way ANOVA | Group | $F_{2,26} = 0.164$ | $p = .849$ n.s. |
| | | Condition | $F_{1,26} = 1.556$ | $p = .223$ n.s. |
| | | Group: Condition | $F_{2,26} = 4.786$ | $p < .05$ * |
| | Pairwise t-test (Group) | Control: Vehicle vs C21 | | $p = .05$ † |
| | | Low: Vehicle vs C21 | | $p = .06$ † |
| | | High: Vehicle vs C21 | | $p = .163$ n.s. |
| | (Condition) | Vehicle: Control vs Low | | $p = .988$ n.s. |
| | | Vehicle: Control vs High | | $p = .069$ † |
| | | Vehicle: Low vs High | | $p = .069$ † |
| | | C21: Control vs Low | | $p = .628$ n.s. |
| | | C21: Control vs High | | $p = .251$ n.s. |
| | | C21: Low vs High | | $p = .128$ n.s. |
| 2E | Three-way ANOVA | Group | $F_{2,26} = 4.446$ | $p < .05$ * |
| | | Demonstrator | $F_{1,26} = 39.731$ | $p < .001$ *** |
| | | Condition | $F_{1,26} = 0.498$ | $p = .487$ n.s. |
| | | Group: Demonstrator | $F_{2,26} = 0.719$ | $p = .497$ n.s. |
| | | Group: Condition | $F_{2,26} = 0.078$ | $p = .926$ n.s. |
| | | Demonstrator: Condition | $F_{1,26} = 0.758$ | $p = .392$ n.s. |
| | | Group: Demonstrator: Condition | $F_{2,26} = 6.151$ | $p < .01$ ** |
| | Two-way ANOVA (Group: Control) | Demonstrator | $F_{1,8} = 23.7$ | $p < .01$ ** |
| | | Condition | $F_{1,8} = 0.351$ | $p = .57$ n.s. |
| | | Demonstrator: Condition | $F_{1,8} = 4.98$ | $p = .056$ † |
| | (Group: Low) | Demonstrator | $F_{1,9} = 9.14$ | $p < .05$ * |
| | | Condition | $F_{1,9} = 0.029$ | $p = .868$ n.s. |
| | | Demonstrator: Condition | $F_{1,9} = 3.97$ | $p = .077$ † |
| | (Group: High) | Demonstrator | $F_{1,9} = 10.0$ | $p < .05$ * |
| | | Condition | $F_{1,9} = 0.212$ | $p = .656$ n.s. |
| | | Demonstrator: Condition | $F_{1,9} = 4.18$ | $p = .071$ † |
| | Two-way ANOVA (Condition: Vehicle) | Group | $F_{2,26} = 2.31$ | $p = .12$ n.s. |
| | | Demonstrator | $F_{1,26} = 19.1$ | $p < .001$ *** |
| | | Group: Demonstrator | $F_{2,26} = 6.21$ | $p < .01$ ** |
| | (Condition: C21) | Group | $F_{2,26} = 3.35$ | $p = .051$ † |
| | | Demonstrator | $F_{1,26} = 6.1$ | $p < .05$ * |

| | | | | |
|---|---|---|---|---|
| | | Group: Demonstrator | $F_{2,26} = 3.37$ | $p = .05$ † |
| | Two-way ANOVA | Group | $F_{2,26} = 3.08$ | $p = .063$ † |
| | (Demonstrator: Neutral) | Condition | $F_{1,26} = 2.03$ | $p = .166$ n.s. |
| | | Group: Condition | $F_{2,26} = 4.98$ | $p < .05$ * |
| | (Demonstrator:Distressed) | Group | $F_{2,26} = 4.41$ | $p < .05$ * |
| | | Condition | $F_{1,26} = 0.002$ | $p = .961$ n.s. |
| | | Group: Condition | $F_{2,26} = 1.86$ | $p = .177$ n.s. |
| | Pairwise t-test | Vehicle: Neutral vs Distressed | | $p < .01$ ** |
| | (Group: Control) | C21: Neutral vs Distressed | | $p = .432$ n.s. |
| | | Neutral: Vehicle vs C21 | | $p = .053$ † |
| | | Distressed: Vehicle vs C21 | | $p = .137$ n.s. |
| | (Group: Low) | Vehicle: Neutral vs Distressed | | $p < .01$ ** |
| | | C21: Neutral vs Distressed | | $p = .903$ n.s. |
| | | Neutral: Vehicle vs C21 | | $p = .122$ n.s. |
| | | Distressed: Vehicle vs C21 | | $p = .384$ n.s. |
| | (Group: High) | Vehicle: Neutral vs Distressed | | $p = .783$ n.s. |
| | | C21: Neutral vs Distressed | | $p < .05$ * |
| | | Neutral: Vehicle vs C21 | | $p = .089$ † |
| | | Distressed: Vehicle vs C21 | | $p = .137$ n.s. |
| | Pairwise t-test | Neutral: Control vs Low | | $p = .232$ n.s. |
| | (Condition: Vehicle) | Neutral: Control vs High | | $p = .632$ n.s. |
| | | Neutral: High vs Low | | $p = .115$ n.s. |
| | | Distressed: Control vs Low | | $p = .131$ n.s. |
| | | Distressed: Control vs High | | $p < .05$ * |
| | | Distressed: High vs Low | | $p = .443$ n.s. |
| | (Condition: C21) | Neutral: Control vs Low | | $p = .085$ † |
| | | Neutral: Control vs High | | $p < .05$ * |
| | | Neutral: High vs Low | | $p = .351$ n.s. |
| | | Distressed: Control vs Low | | $p = .698$ n.s. |
| | | Distressed: Control vs High | | $p = .198$ n.s. |
| | | Distressed: High vs Low | | $p = .15$ n.s. |
| 2F | Two-way ANOVA | Group | $F_{2,26} = 0.164$ | $p = .052$ † |
| | | Condition | $F_{1,26} = 1.556$ | $p = .988$ n.s. |
| | | Group: Condition | $F_{2,26} = 4.786$ | $p = .493$ n.s. |
| | Pairwise t-test | Control: Vehicle vs C21 | | $p = .19$ n.s. |
| | (Group) | Low: Vehicle vs C21 | | $p = .689$ n.s. |
| | | High: Vehicle vs C21 | | $p = .628$ n.s. |
| | (Condition) | Vehicle: Control vs Low | | $p = 1$ n.s. |
| | | Vehicle: Control vs High | | $p = .281$ n.s. |
| | | Vehicle: Low vs High | | $p = .693$ n.s. |

| | | | | | |
|---|---|---|---|---|---|
| | | C21: Control vs Low | | | $p$ =.565 n.s. |
| | | C21: Control vs High | | | $p$ =.065 † |
| | | C21: Low vs High | | | $p$ =.815 n.s. |
| 2G | One-sample t-test | Control (Vehicle) vs chance level | $t_{7}$ | =17.771 | $p$ <.001 *** |
| | | Control (C21) vs chance level | $t_{7}$ | =-1.853 | $p$ =.106 n.s. |
| | | Low (Vehicle) vs chance level | $t_{6}$ | =3.605 | $p$ <.05 * |
| | | Low(C21) vs chance level | $t_{6}$ | =0.990 | $p$ =.360 n.s. |
| | | High (Vehicle) vs chance level | $t_{8}$ | =0.184 | $p$ =.859 n.s. |
| | | High (C21) vs chance level | $t_{8}$ | =4.943 | $p$ <.01 ** |
| | Two-way ANOVA | Group | $F_{2,21}$ | =0.119 | $p$ =.507 n.s. |
| | | Condition | $F_{1,21}$ | =4.597 | $p$ <.05 * |
| | | Group: Condition | $F_{2,21}$ | =16.298 | $p$ <.001 *** |
| | Pairwise t-test | Control: Vehicle vs C21 | | | $p$ <.001 *** |
| | (Group) | Low: Vehicle vs C21 | | | $p$ =.236 n.s. |
| | | High: Vehicle vs C21 | | | $p$ <.05 * |
| | (Condition) | Vehicle: Control vs Low | | | $p$ =.576 n.s. |
| | | Vehicle: Control vs High | | | $p$ <.01 ** |
| | | Vehicle: Low vs High | | | $p$ =.071 † |
| | | C21: Control vs Low | | | $p$ =.235 n.s. |
| | | C21: Control vs High | | | $p$ <.01 ** |
| | | C21: Low vs High | | | $p$ =.335 n.s. |
| 2H | Three-way ANOVA | Group | $F_{2,21}$ | =0.842 | $p$ =.445 n.s. |
| | | Demonstrator | $F_{1,21}$ | =49.911 | $p$ <.001 *** |
| | | Condition | $F_{1,21}$ | =0.140 | $p$ =.712 n.s. |
| | | Group: Demonstrator | $F_{2,21}$ | =0.093 | $p$ =.911 n.s. |
| | | Group: Condition | $F_{2,21}$ | =2.865 | $p$ =.079 † |
| | | Demonstrator: Condition | $F_{1,21}$ | =4.827 | $p$ <.05 * |
| | | Group: Demonstrator: Condition | $F_{2,21}$ | =12.462 | $p$ <.001 *** |
| | Two-way ANOVA | Demonstrator | $F_{1,7}$ | =46.7 | $p$ <.001 *** |
| | (Group: Control) | Condition | $F_{1,7}$ | =2.07 | $p$ =.193 n.s. |
| | | Demonstrator: Condition | $F_{1,7}$ | =33.5 | $p$ <.001 *** |
| | (Group: Low) | Demonstrator | $F_{1,6}$ | =14.5 | $p$ <.01 ** |
| | | Condition | $F_{1,6}$ | =1.01 | $p$ =.353 n.s. |
| | | Demonstrator: Condition | $F_{1,6}$ | =1.13 | $p$ =.328 n.s. |
| | (Group: High) | Demonstrator | $F_{1,8}$ | =11.9 | $p$ <.05 * |
| | | Condition | $F_{1,8}$ | =2.29 | $p$ =.174 n.s. |
| | | Demonstrator: Condition | $F_{1,8}$ | =5.88 | $p$ <.05 * |
| | Two-way ANOVA | Group | $F_{2,21}$ | =2.59 | $p$ =.099 † |
| | (Condition: Vehicle) | Demonstrator | $F_{1,21}$ | =36.1 | $p$ <.001 *** |
| | | Group: Demonstrator | $F_{2,21}$ | =8.90 | $p$ <.01 ** |

| | | | | | |
|---|---|---|---|---|---|
| | (Condition: C21) | Group | $F_{2,21}$ = 0.41 | $p$ = .669 | n.s. |
| | | Demonstrator | $F_{1,21}$ = 5.36 | $p$ < .05 | * |
| | | Group: Demonstrator | $F_{2,21}$ = 7.60 | $p$ < .01 | ** |
| | Two-way ANOVA | Group | $F_{2,21}$ = 0.507 | $p$ = .61 | n.s. |
| | (Demonstrator: Neutral) | Condition | $F_{1,21}$ = 2.16 | $p$ = .158 | n.s. |
| | | Group: Condition | $F_{2,21}$ = 2.36 | $p$ = .121 | n.s. |
| | (Demonstrator:Distressed) | Group | $F_{2,21}$ = 0.7 | $p$ = .508 | n.s. |
| | | Condition | $F_{1,21}$ = 1.59 | $p$ = .221 | n.s. |
| | | Group: Condition | $F_{2,21}$ = 7.11 | $p$ < .01 | ** |
| | Pairwise t-test | Vehicle: Neutral vs Distressed | | $p$ < .001 | *** |
| | (Group: Control) | C21: Neutral vs Distressed | | $p$ = .175 | n.s. |
| | | Neutral: Vehicle vs C21 | | $p$ = .056 | † |
| | | Distressed: Vehicle vs C21 | | $p$ < .01 | ** |
| | (Group: Low) | Vehicle: Neutral vs Distressed | | $p$ < .05 | * |
| | | C21: Neutral vs Distressed | | $p$ = .456 | n.s. |
| | | Neutral: Vehicle vs C21 | | $p$ = .879 | n.s. |
| | | Distressed: Vehicle vs C21 | | $p$ = .333 | n.s. |
| | (Group: High) | Vehicle: Neutral vs Distressed | | $p$ = .573 | n.s. |
| | | C21: Neutral vs Distressed | | $p$ < .01 | ** |
| | | Neutral: Vehicle vs C21 | | $p$ = .936 | n.s. |
| | | Distressed: Vehicle vs C21 | | $p$ = .062 | † |
| | Pairwise t-test | Neutral: Control vs Low | | $p$ = 1 | n.s. |
| | (Condition: Vehicle) | Neutral: Control vs High | | $p$ = 1 | n.s. |
| | | Neutral: High vs Low | | $p$ = 1 | n.s. |
| | | Distressed: Control vs Low | | $p$ = .676 | n.s. |
| | | Distressed: Control vs High | | $p$ < .05 | * |
| | | Distressed: High vs Low | | $p$ = .294 | n.s. |
| | (Condition: C21) | Neutral: Control vs Low | | $p$ = .506 | n.s. |
| | | Neutral: Control vs High | | $p$ = .23 | n.s. |
| | | Neutral: High vs Low | | $p$ = 1 | n.s. |
| | | Distressed: Control vs Low | | $p$ = 1 | n.s. |
| | | Distressed: Control vs High | | $p$ = .2 | n.s. |
| | | Distressed: High vs Low | | $p$ = .315 | n.s. |
| 2I | Two-way ANOVA | Group | $F_{2,21}$ = 8.305 | $p$ < .01 | ** |
| | | Condition | $F_{1,21}$ = 0.250 | $p$ = .623 | n.s. |
| | | Group: Condition | $F_{2,21}$ = 1.022 | $p$ = .377 | n.s. |
| | Pairwise t-test | Control: Vehicle vs C21 | | $p$ = .268 | n.s. |
| | (Group) | Low: Vehicle vs C21 | | $p$ = .261 | n.s. |
| | | High: Vehicle vs C21 | | $p$ = .43 | n.s. |
| | (Condition) | Vehicle: Control vs Low | | $p$ = 1 | n.s. |

| | | | | | | |
|---|---|---|---|---|---|---|
| | | Vehicle: Control vs High | | | $p <.01$ | ** |
| | | Vehicle: Low vs High | | | $p <.05$ | * |
| | | C21: Control vs Low | | | $p = 1$ | n.s. |
| | | C21: Control vs High | | | $p = .149$ | n.s. |
| | | C21: Low vs High | | | $p = .131$ | n.s. |
| 2J | Type III Likelihood Ratio Chi-Square Test | Group | $X^2$ 2 | $= 257.58$ | $p < .001$ | *** |
| | | Condition | $X^2$ 1 | $= 504.98$ | $p < .001$ | *** |
| | | Group: Condition | $X^2$ 2 | $= 2076.39$ | $p < .001$ | *** |
| | Tukey-adjusted EMM comparisons (Group) | Control: Neutral vs Distressed | | | $p < .001$ | *** |
| | | Low: Neutral vs Distressed | | | $p < .001$ | *** |
| | | High: Neutral vs Distressed | | | $p < .001$ | *** |
| | (Condition) | Vehicle: Control vs Low | | | $p < .001$ | *** |
| | | Vehicle: Control vs High | | | $p < .001$ | *** |
| | | Vehicle: Low vs High | | | $p < .001$ | *** |
| | | C21: Control vs Low | | | $p < .001$ | *** |
| | | C21: Control vs High | | | $p < .001$ | *** |
| | | C21: Low vs High | | | $p = .451$ | n.s. |
| 2K | Two-way ANOVA | Group | $F_{2,21}$ | $= 30.336$ | $p < .001$ | *** |
| | | Condition | $F_{1,21}$ | $= 3.500$ | $p = .075$ | † |
| | | Group: Condition | $F_{2,21}$ | $= 16.709$ | $p < .001$ | *** |
| | Pairwise t-test (Group) | Control: Vehicle vs C21 | | | $p < .05$ | * |
| | | Low: Vehicle vs C21 | | | $p < .001$ | *** |
| | | High: Vehicle vs C21 | | | $p < .01$ | ** |
| | (Condition) | Vehicle: Control vs Low | | | $p < .001$ | *** |
| | | Vehicle: Control vs High | | | $p < .001$ | *** |
| | | Vehicle: Low vs High | | | $p = .88$ | n.s. |
| | | C21: Control vs Low | | | $p = .27$ | n.s. |
| | | C21: Control vs High | | | $p = .556$ | n.s. |
| | | C21: Low vs High | | | $p = 1$ | n.s. |
| 2L | Two-way ANOVA | Group | $F_{2,21}$ | $= 3.361$ | $p = .054$ | † |
| | | Condition | $F_{1,21}$ | $= 0.092$ | $p = .765$ | n.s. |
| | | Group: Condition | $F_{2,21}$ | $= 0.723$ | $p = .497$ | n.s. |
| | Pairwise t-test (Group) | Control: Vehicle vs C21 | | | $p = .775$ | n.s. |
| | | Low: Vehicle vs C21 | | | $p = .052$ | † |
| | | High: Vehicle vs C21 | | | $p = .845$ | n.s. |
| | (Condition) | Vehicle: Control vs Low | | | $p < .05$ | * |
| | | Vehicle: Control vs High | | | $p = .155$ | n.s. |
| | | Vehicle: Low vs High | | | $p = 1$ | n.s. |
| | | C21: Control vs Low | | | $p = .341$ | n.s. |
| | | C21: Control vs High | | | $p = .616$ | n.s. |

| | | | | | | |
|---|---|---|---|---|---|---|
| | | C21: Low vs High | | | $p = 1$ | n.s. |
| 2M | Two-way ANOVA | Group | $F_{2,21}$ | $= 5.166$ | $p < .05$ | * |
| | | Condition | $F_{1,21}$ | $= 0.012$ | $p = .914$ | n.s. |
| | | Group: Condition | $F_{2,21}$ | $= 2.072$ | $p = .151$ | n.s. |
| | Pairwise t-test | Control: Vehicle vs C21 | | | $p = .607$ | n.s. |
| | (Group) | Low: Vehicle vs C21 | | | $p = .186$ | n.s. |
| | | High: Vehicle vs C21 | | | $p = .302$ | n.s. |
| | (Condition) | Vehicle: Control vs Low | | | $p = .092$ | † |
| | | Vehicle: Control vs High | | | $p < .01$ | ** |
| | | Vehicle: Low vs High | | | $p = .664$ | n.s. |
| | | C21: Control vs Low | | | $p = .071$ | † |
| | | C21: Control vs High | | | $p = .626$ | n.s. |
| | | C21: Low vs High | | | $p = .958$ | n.s. |
| 2N | Type III Likelihood Ratio | Group | $X^2_{2}$ | $= 127.39$ | $p < .001$ | *** |
| | Chi-Square Test | Condition | $X^2_{1}$ | $= 127.88$ | $p < .001$ | *** |
| | | Group: Condition | $X^2_{2}$ | $= 980.59$ | $p < .001$ | *** |
| | Tukey-adjusted EMM | Control: Neutral vs Distressed | | | $p < .001$ | *** |
| | comparisons (Group) | Low: Neutral vs Distressed | | | $p < .001$ | *** |
| | | High: Neutral vs Distressed | | | $p < .001$ | *** |
| | (Condition) | Vehicle: Control vs Low | | | $p < .001$ | *** |
| | | Vehicle: Control vs High | | | $p < .001$ | *** |
| | | Vehicle: Low vs High | | | $p < .001$ | *** |
| | | C21: Control vs Low | | | $p < .001$ | *** |
| | | C21: Control vs High | | | $p < .001$ | *** |
| | | C21: Low vs High | | | $p < .001$ | *** |
| 2O | Two-way ANOVA | Group | $F_{2,22}$ | $= 0.476$ | $p = .633$ | n.s. |
| | | Condition | $F_{1,22}$ | $= 8.323$ | $p < .01$ | ** |
| | | Group: Condition | $F_{2,22}$ | $= 8.977$ | $p < .01$ | ** |
| | Pairwise t-test | Control: Vehicle vs C21 | | | $p < .05$ | * |
| | (Group) | Low: Vehicle vs C21 | | | $p < .05$ | * |
| | | High: Vehicle vs C21 | | | $p < .01$ | ** |
| | (Condition) | Vehicle: Control vs Low | | | $p = .109$ | n.s. |
| | | Vehicle: Control vs High | | | $p < .05$ | * |
| | | Vehicle: Low vs High | | | $p = 1$ | n.s. |
| | | C21: Control vs Low | | | $p = 1$ | n.s. |
| | | C21: Control vs High | | | $p = 1$ | n.s. |
| | | C21: Low vs High | | | $p = 1$ | n.s. |
| 2P | Two-way ANOVA | Group | $F_{2,22}$ | $= 1.474$ | $p = .251$ | n.s. |
| | | Condition | $F_{1,22}$ | $= 5.477$ | $p < .05$ | * |
| | | Group: Condition | $F_{2,22}$ | $= 0.693$ | $p = .511$ | n.s. |

| | | | | |
|---|---|---|---|---|
| | Pairwise t-test | Control: Vehicle vs C21 | | $p$ =.493 n.s. |
| | (Group) | Low: Vehicle vs C21 | | $p$ =.202 n.s. |
| | | High: Vehicle vs C21 | | $p$ =.068 † |
| | (Condition) | Vehicle: Control vs Low | | $p$ =.325 n.s. |
| | | Vehicle: Control vs High | | $p$ =.129 n.s. |
| | | Vehicle: Low vs High | | $p$ =1 n.s. |
| | | C21: Control vs Low | | $p$ =1 n.s. |
| | | C21: Control vs High | | $p$ =.961 n.s. |
| | | C21: Low vs High | | $p$ =1 n.s. |
| 2Q | Two-way ANOVA | Group | $F_{2,22}$ =2.376 | $p$ =.116 n.s. |
| | | Condition | $F_{1,22}$ =1.860 | $p$ =.186 n.s. |
| | | Group: Condition | $F_{2,22}$ =0.389 | $p$ =.682 n.s. |

**Table S2. Detailed statistical results corresponding to Figure 2.**

| Figure | Analysis | Factor | Statistical value | p-value |
|---|---|---|---|---|
| 3D | One-sample t-test | Vehicle vs chance level | $t_{11} = 5.861$ | $p < .001$ *** |
| | | C21 vs chance level | $t_{11} = -2.382$ | $p < .05$ * |
| | One-way ANOVA | Condition | $F_{1,11} = 29.877$ | $p < .001$ *** |
| | Pairwise t-test | Vehicle vs C21 | | $p < .001$ *** |
| 3E | Two-way ANOVA | Condition | $F_{1,11} = 0.080$ | $p = .782$ n.s. |
| | | Demonstrator | $F_{1,11} = 4.594$ | $p = .055$ † |
| | | Condition:Demonstrator | $F_{1,11} = 36.650$ | $p < .001$ *** |
| | Pairwise t-test (Condition) | Vehicle: Neutral vs Distressed | | $p < .001$ *** |
| | | C21: Neutral vs Distressed | | $p = .083$ † |
| | (Demonstrator) | Neutral: Vehicle vs C21 | | $p < .01$ ** |
| | | Distressed: Vehicle vs C21 | | $p < .05$ * |
| 3F | One-way ANOVA | Condition | $F_{1,11} = 0.019$ | $p = .892$ n.s. |
| 3G | One-sample t-test | Vehicle vs chance level | $t_{10} = 6.170$ | $p < .001$ *** |
| | | C21 vs chance level | $t_{10} = 0.632$ | $p = .542$ n.s. |
| | One-way ANOVA | Condition | $F_{1,10} = 19.516$ | $p < .01$ ** |
| | Pairwise t-test | Vehicle vs C21 | | $p < .01$ ** |
| 3H | Two-way ANOVA | Condition | $F_{1,10} = 1.583$ | $p = .782$ n.s. |
| | | Demonstrator | $F_{1,10} = 10.775$ | $p < .01$ ** |
| | | Condition:Demonstrator | $F_{1,10} = 13.191$ | $p < .01$ ** |
| | Pairwise t-test (Condition) | Vehicle: Neutral vs Distressed | | $p < .01$ ** |
| | | C21: Neutral vs Distressed | | $p = .681$ n.s. |
| | (Demonstrator) | Neutral: Vehicle vs C21 | | $p < .01$ ** |
| | | Distressed: Vehicle vs C21 | | $p = .416$ n.s. |
| 3I | One-way ANOVA | Condition | $F_{1,10} = 0.109$ | $p = .748$ n.s. |
| 3J | Type III Likelihood Ratio Chi-Square Test | Condition | $X^2_{1} = 257.58$ | $p < .001$ *** |
| | Tukey-adjusted EMM comparisons | Vehicle vs C21 | | $p < .001$ *** |
| 3K | One-way ANOVA | Condition | $F_{1,11} = 2.962$ | $p = .113$ n.s. |
| 3L | One-way ANOVA | Condition | $F_{1,11} = 1.476$ | $p = .25$ n.s. |
| 3M | One-way ANOVA | Condition | $F_{1,11} = 0.062$ | $p = .808$ n.s. |
| 3N | Type III Likelihood Ratio Chi-Square Test | Condition | $X^2_{1} = 267.53$ | $p < .001$ *** |
| | Tukey-adjusted EMM comparisons | Vehicle vs C21 | | $p < .001$ *** |
| 3O | One-way ANOVA | Condition | $F_{1,10} = 5.623$ | $p < .05$ * |
| | Pairwise t-test | Vehicle vs C21 | | $p < .05$ * |
| 3P | One-way ANOVA | Condition | $F_{1,10} = 2.047$ | $p = .183$ n.s. |

| | | | | |
|---|---|---|---|---|
| 3Q | One-way ANOVA | Condition | $F_{1,10} = 0.121$ | $p = .735$ n.s. |

**Table S3. Detailed statistical results corresponding to Figure 3.**

| Figure | Analysis | Factor | Statistical value | p-value | |
|---|---|---|---|---|---|
| 4B | One-way ANOVA | Group | $F_{2,13} = 4.063$ | $p < .05$ | * |
| | Pairwise t-test | Control vs Low | | $p = 1$ | n.s. |
| | | Control vs High | | $p = .063$ | † |
| | | Low vs High | | $p = .085$ | † |
| 4C | One-way ANOVA | Group | $F_{2,15} = 13.734$ | $p < .001$ | *** |
| | Pairwise t-test | Control vs Low | | $p = .152$ | n.s. |
| | | Control vs High | | $p < .001$ | *** |
| | | Low vs High | | $p < .05$ | * |
| 4E | One-way ANOVA | Group | $F_{2,13} = 17.052$ | $p < .001$ | *** |
| | Pairwise t-test | Control vs Low | | $p < .05$ | * |
| | | Control vs High | | $p < .001$ | *** |
| | | Low vs High | | $p = .156$ | n.s. |
| 4F | One-way ANOVA | Group | $F_{2,13} = 43.507$ | $p < .001$ | *** |
| | Pairwise t-test | Control vs Low | | $p < .001$ | *** |
| | | Control vs High | | $p < .001$ | *** |
| | | Low vs High | | $p = .089$ | † |
| 4G | One-way ANOVA | Group | $F_{2,13} = 3.531$ | $p = .06$ | † |
| | Pairwise t-test | Control vs Low | | $p = .826$ | n.s. |
| | | Control vs High | | $p = .059$ | † |
| | | Low vs High | | $p = .51$ | n.s. |
| 4H | One-way ANOVA | Group | $F_{2,13} = 17.103$ | $p < .001$ | *** |
| | Pairwise t-test | Control vs Low | | $p < .01$ | ** |
| | | Control vs High | | $p < .001$ | *** |
| | | Low vs High | | $p = .337$ | n.s. |
| 4I | One-way ANOVA | Group | $F_{2,13} = 17.654$ | $p < .001$ | *** |
| | Pairwise t-test | Control vs Low | | $p < .01$ | ** |
| | | Control vs High | | $p < .001$ | *** |
| | | Low vs High | | $p = .637$ | n.s. |
| 4I | One-way ANOVA | Group | $F_{2,13} = 9.339$ | $p < .01$ | *** |
| | Pairwise t-test | Control vs Low | | $p = .096$ | † |
| | | Control vs High | | $p < .01$ | ** |
| | | Low vs High | | $p = .281$ | n.s. |

**Table S4. Detailed statistical results corresponding to Figure 4.**

| Figure | Analysis | Factor | Statistical value | p-value |
|---|---|---|---|---|
| 5D | One-sample t-test | Control (Vehicle) vs chance level | $t_{8}$ = 4.172 | $p$ <.01 ** |
| | | Control (C21) vs chance level | $t_{8}$ 5.876 | $p$ <.001 *** |
| | | Low (Vehicle) vs chance level | $t_{5}$ = 2.966 | $p$ <.05 * |
| | | Low(C21) vs chance level | $t_{5}$ = 2.100 | $p$ =.090 † |
| | | High (Vehicle) vs chance level | $t_{7}$ = -1.603 | $p$ =.153 n.s. |
| | | High (C21) vs chance level | $t_{7}$ = 2.304 | $p$ =.055 † |
| | Two-way ANOVA | Group | $F_{2,20}$ = 7.945 | $p$ <.01 ** |
| | | Condition | $F_{1,20}$ = 1.834 | $p$ =.191 n.s. |
| | | Group: Condition | $F_{2,20}$ = 3.181 | $p$ =.063 † |
| | Pairwise t-test | Control: Vehicle vs C21 | | $p$ =.251 n.s. |
| | (Group) | Low: Vehicle vs C21 | | $p$ =.475 n.s. |
| | | High: Vehicle vs C21 | | $p$ <.05 * |
| | (Condition) | Vehicle: Control vs Low | | $p$ =1 n.s. |
| | | Vehicle: Control vs High | | $p$ <.01 ** |
| | | Vehicle: Low vs High | | $p$ <.01 ** |
| | | C21: Control vs Low | | $p$ =.36 n.s. |
| | | C21: Control vs High | | $p$ =.197 n.s. |
| | | C21: Low vs High | | $p$ =1 n.s. |
| 5E | Three-way ANOVA | Group | $F_{2,20}$ = 5.920 | $p$ <.05 * |
| | | Demonstrator | $F_{1,20}$ = 32.384 | $p$ <.001 *** |
| | | Condition | $F_{1,20}$ = 6.659 | $p$ <.05 * |
| | | Group: Demonstrator | $F_{2,20}$ = 6.616 | $p$ <.01 ** |
| | | Group: Condition | $F_{2,20}$ = 1.685 | $p$ =.211 n.s. |
| | | Demonstrator: Condition | $F_{1,20}$ = 2.118 | $p$ =.161 n.s. |
| | | Group: Demonstrator: Condition | $F_{2,20}$ = 3.390 | $p$ =.054 † |
| | Two-way ANOVA | Demonstrator | $F_{1,8}$ = 31.2 | $p$ <.001 *** |
| | (Group: Control) | Condition | $F_{1,8}$ = 6.01 | $p$ <.05 * |
| | | Demonstrator: Condition | $F_{1,8}$ = 4.45 | $p$ =.068 † |
| | (Group: Low) | Demonstrator | $F$ = 5.96 | $p$ =.058 † |
| | | Condition | $F$ = 0.004 | $p$ =.95 n.s. |
| | | Demonstrator: Condition | $F$ = 1.10 | $p$ =.343 n.s. |
| | (Group: High) | Demonstrator | $F$ = 2.15 | $p$ =.186 n.s. |
| | | Condition | $F$ = 9.74 | $p$ <.05 * |
| | | Demonstrator: Condition | $F$ = 6.74 | $p$ <.05 * |
| | Two-way ANOVA | Group | $F_{2,20}$ = 3.94 | $p$ <.05 * |
| | (Condition: Vehicle) | Demonstrator | $F_{1,20}$ = 12.8 | $p$ <.01 ** |
| | | Group: Demonstrator | $F_{2,20}$ = 5.90 | $p$ <.05 * |
| | (Condition: C21) | Group | $F_{2,20}$ = 5.70 | $p$ <.05 * |
| | | Demonstrator | $F_{1,20}$ = 25.3 | $p$ <.001 *** |

| | | | | | |
|---|---|---|---|---|---|
| | | Group: Demonstrator | $F_{2,20}$ = 4.73 | $p$ <.05 | * |
| | Two-way ANOVA | Group | $F_{2,20}$ = 0.634 | $p$ =.541 | n.s. |
| | (Demonstrator: Neutral) | Condition | $F_{1,20}$ = 0.404 | $p$ =.532 | n.s. |
| | | Group: Condition | $F_{2,20}$ = 0.677 | $p$ =.519 | n.s. |
| | (Demonstrator:Distressed) | Group | $F_{2,20}$ = 7.35 | $p$ <.01 | ** |
| | | Condition | $F_{1,20}$ = 5.32 | $p$ <.05 | * |
| | | Group: Condition | $F_{2,20}$ = 3.55 | $p$ <.05 | * |
| | Pairwise t-test | Vehicle: Neutral vs Distressed | | $p$ <.01 | ** |
| | (Group: Control) | C21: Neutral vs Distressed | | $p$ <.01 | ** |
| | | Neutral: Vehicle vs C21 | | $p$ =.928 | n.s. |
| | | Distressed: Vehicle vs C21 | | $p$ <.05 | * |
| | (Group: Low) | Vehicle: Neutral vs Distressed | | $p$ =.106 | n.s. |
| | | C21: Neutral vs Distressed | | $p$ =.081 | † |
| | | Neutral: Vehicle vs C21 | | $p$ =.282 | n.s. |
| | | Distressed: Vehicle vs C21 | | $p$ =.526 | n.s. |
| | (Group: High) | Vehicle: Neutral vs Distressed | | $p$ =.15 | n.s. |
| | | C21: Neutral vs Distressed | | $p$ =.054 | † |
| | | Neutral: Vehicle vs C21 | | $p$ =.858 | n.s. |
| | | Distressed: Vehicle vs C21 | | $p$ <.01 | ** |
| | Pairwise t-test | Neutral: Control vs Low | | $p$ =.528 | n.s. |
| | (Condition: Vehicle) | Neutral: Control vs High | | $p$ =1 | n.s. |
| | | Neutral: High vs Low | | $p$ =.652 | n.s. |
| | | Distressed: Control vs Low | | $p$ =1 | n.s. |
| | | Distressed: Control vs High | | $p$ <.05 | * |
| | | Distressed: High vs Low | | $p$ =.053 | † |
| | (Condition: C21) | Neutral: Control vs Low | | $p$ =1 | n.s. |
| | | Neutral: Control vs High | | $p$ =1 | n.s. |
| | | Neutral: High vs Low | | $p$ =1 | n.s. |
| | | Distressed: Control vs Low | | $p$ <.05 | * |
| | | Distressed: Control vs High | | $p$ <.05 | * |
| | | Distressed: High vs Low | | $p$ =1 | n.s. |
| 5F | Two-way ANOVA | Group | $F_{2,20}$ = 4.536 | $p$ <.05 | * |
| | | Condition | $F_{1,20}$ = 0.683 | $p$ =.418 | n.s. |
| | | Group: Condition | $F_{2,20}$ = 0.415 | $p$ =.666 | n.s. |
| | Pairwise t-test | Control: Vehicle vs C21 | | $p$ =.702 | n.s. |
| | (Group) | Low: Vehicle vs C21 | | $p$ =.388 | n.s. |
| | | High: Vehicle vs C21 | | $p$ =.564 | n.s. |
| | (Condition) | Vehicle: Control vs Low | | $p$ =.602 | n.s. |
| | | Vehicle: Control vs High | | $p$ <.05 | * |
| | | Vehicle: Low vs High | | $p$ =.634 | n.s. |

| | | | | | |
|---|---|---|---|---|---|
| | | C21: Control vs Low | | $p = 1$ | n.s. |
| | | C21: Control vs High | | $p = .098$ | † |
| | | C21: Low vs High | | $p = .493$ | n.s. |
| 5G | One-sample t-test | Control (Vehicle) vs chance level | $t_{9} = 6.320$ | $p < .001$ | *** |
| | | Control (C21) vs chance level | $t_{9} = 7.037$ | $p < .001$ | *** |
| | | Low (Vehicle) vs chance level | $t_{6} = 6.742$ | $p < .001$ | *** |
| | | Low(C21) vs chance level | $t_{6} = 3.712$ | $p < .001$ | *** |
| | | High (Vehicle) vs chance level | $t_{7} = 0.639$ | $p = .543$ | n.s. |
| | | High (C21) vs chance level | $t_{7} = 4.243$ | $p < .01$ | ** |
| | Two-way ANOVA | Group | $F_{2,22} = 5.658$ | $p < .05$ | * |
| | | Condition | $F_{1,22} = 1.166$ | $p = .292$ | n.s. |
| | | Group: Condition | $F_{2,22} = 2.822$ | $p = .081$ | † |
| | Pairwise t-test | Control: Vehicle vs C21 | | $p = .92$ | n.s. |
| | (Group) | Low: Vehicle vs C21 | | $p = .646$ | n.s. |
| | | High: Vehicle vs C21 | | $p < .05$ | * |
| | (Condition) | Vehicle: Control vs Low | | $p = 1$ | n.s. |
| | | Vehicle: Control vs High | | $p < .01$ | ** |
| | | Vehicle: Low vs High | | $p < .01$ | ** |
| | | C21: Control vs Low | | $p = 1$ | n.s. |
| | | C21: Control vs High | | $p = .863$ | n.s. |
| | | C21: Low vs High | | $p = 1$ | n.s. |
| 5H | Three-way ANOVA | Group | $F_{2,22} = 0.756$ | $p = .481$ | n.s. |
| | | Demonstrator | $F_{1,22} = 65.659$ | $p < .001$ | *** |
| | | Condition | $F_{1,22} = 0.444$ | $p = .512$ | n.s. |
| | | Group: Demonstrator | $F_{2,22} = 3.594$ | $p < .05$ | * |
| | | Group: Condition | $F_{2,22} = 2.444$ | $p = .110$ | n.s. |
| | | Demonstrator: Condition | $F_{1,22} = 0.516$ | $p = .480$ | n.s. |
| | | Group: Demonstrator: Condition | $F_{2,22} = 1.823$ | $p = .185$ | n.s. |
| | Two-way ANOVA | Demonstrator | $F_{1,9} = 40.8$ | $p < .001$ | *** |
| | (Group: Control) | Condition | $F_{1,9} = 1.40$ | $p = .268$ | n.s. |
| | | Demonstrator: Condition | $F_{1,9} = 0.706$ | $p = .423$ | n.s. |
| | (Group: Low) | Demonstrator | $F_{1,6} = 19.1$ | $p < .01$ | ** |
| | | Condition | $F_{1,6} = 2.90$ | $p = .14$ | n.s. |
| | | Demonstrator: Condition | $F_{1,6} = 0.768$ | $p = .414$ | n.s. |
| | (Group: High) | Demonstrator | $F_{1,7} = 10.7$ | $p < .05$ | * |
| | | Condition | $F_{1,7} = 0.008$ | $p = .931$ | n.s. |
| | | Demonstrator: Condition | $F_{1,7} = 2.58$ | $p = .152$ | n.s. |
| | Two-way ANOVA | Group | $F_{2,22} = 0.216$ | $p = .808$ | n.s. |
| | (Condition: Vehicle) | Demonstrator | $F_{1,22} = 35.6$ | $p < .001$ | *** |
| | | Group: Demonstrator | $F_{2,22} = 5.03$ | $p < .05$ | * |

| | | | | | |
|---|---|---|---|---|---|
| | (Condition: C21) | Group | $F_{2,22}$ = 2.79 | $p$ = .083 | † |
| | | Demonstrator | $F_{1,22}$ = 42.1 | $p$ < .001 | *** |
| | | Group: Demonstrator | $F_{2,22}$ = 0.971 | $p$ = .394 | n.s. |
| | Two-way ANOVA | Group | $F_{2,22}$ = 5.42 | $p$ < .05 | * |
| | (Demonstrator: Neutral) | Condition | $F_{1,22}$ = 3.92 | $p$ = .06 | † |
| | | Group: Condition | $F_{2,22}$ = 2.98 | $p$ = .071 | † |
| | (Demonstrator:Distressed) | Group | $F_{2,22}$ = 0.871 | $p$ = .433 | n.s. |
| | | Condition | $F_{1,22}$ = 0.0003 | $p$ = .987 | n.s. |
| | | Group: Condition | $F_{2,22}$ = 2.60 | $p$ = .152 | n.s. |
| | Pairwise t-test | Vehicle: Neutral vs Distressed | | $p$ < .001 | *** |
| | (Group: Control) | C21: Neutral vs Distressed | | $p$ < .001 | *** |
| | | Neutral: Vehicle vs C21 | | $p$ = .563 | n.s. |
| | | Distressed: Vehicle vs C21 | | $p$ = .268 | n.s. |
| | (Group: Low) | Vehicle: Neutral vs Distressed | | $p$ < .001 | *** |
| | | C21: Neutral vs Distressed | | $p$ < .05 | * |
| | | Neutral: Vehicle vs C21 | | $p$ = .152 | n.s. |
| | | Distressed: Vehicle vs C21 | | $p$ = .221 | n.s. |
| | (Group: High) | Vehicle: Neutral vs Distressed | | $p$ = .439 | n.s. |
| | | C21: Neutral vs Distressed | | $p$ < .01 | ** |
| | | Neutral: Vehicle vs C21 | | $p$ < .01 | ** |
| | | Distressed: Vehicle vs C21 | | $p$ = .465 | n.s. |
| | Pairwise t-test | Neutral: Control vs Low | | $p$ = 1 | n.s. |
| | (Condition: Vehicle) | Neutral: Control vs High | | $p$ < .05 | * |
| | | Neutral: High vs Low | | $p$ < .05 | * |
| | | Distressed: Control vs Low | | $p$ = 1 | n.s. |
| | | Distressed: Control vs High | | $p$ = 1 | n.s. |
| | | Distressed: High vs Low | | $p$ = .413 | n.s. |
| | (Condition: C21) | Neutral: Control vs Low | | $p$ = .216 | n.s. |
| | | Neutral: Control vs High | | $p$ = .979 | n.s. |
| | | Neutral: High vs Low | | $p$ < .05 | * |
| | | Distressed: Control vs Low | | $p$ = .279 | n.s. |
| | | Distressed: Control vs High | | $p$ = 1 | n.s. |
| | | Distressed: High vs Low | | $p$ = 1 | n.s. |
| 5I | Two-way ANOVA | Group | $F_{2,22}$ = 0.251 | $p$ = .781 | n.s. |
| | | Condition | $F_{1,22}$ = 0.010 | $p$ = .923 | n.s. |
| | | Group: Condition | $F_{2,22}$ = 0.566 | $p$ = .576 | n.s. |
| 5J | Type III Likelihood Ratio | Group | $X^2_{2}$ = 2112.42 | $p$ < .001 | *** |
| | Chi-Square Test | Condition | $X^2_{1}$ = 90.33 | $p$ < .001 | *** |
| | | Group: Condition | $X^2_{2}$ = 292.87 | $p$ < .001 | *** |
| | | Control: Neutral vs Distressed | | $p$ < .001 | *** |

| | | | | |
|---|---|---|---|---|
| | Tukey-adjusted EMM comparisons (Group) | Low: Neutral vs Distressed | | $p$ <.001 *** |
| | | High: Neutral vs Distressed | | $p$ <.001 *** |
| | (Condition) | Vehicle: Control vs Low | | $p$ <.001 *** |
| | | Vehicle: Control vs High | | $p$ <.001 *** |
| | | Vehicle: Low vs High | | $p$ <.001 *** |
| | | C21: Control vs Low | | $p$ <.001 *** |
| | | C21: Control vs High | | $p$ <.001 *** |
| | | C21: Low vs High | | $p$ <.001 *** |
| 5K | Two-way ANOVA | Group | $F_{2,20}$ =6.802 | $p$ <.01 ** |
| | | Condition | $F_{1,20}$ =5.238 | $p$ <.05 * |
| | | Group: Condition | $F_{2,20}$ =1.278 | $p$ =.300 n.s. |
| | Pairwise t-test (Group) | Control: Vehicle vs C21 | | $p$ =.541 n.s. |
| | | Low: Vehicle vs C21 | | $p$ =.297 n.s. |
| | | High: Vehicle vs C21 | | $p$ <.001 *** |
| | (Condition) | Vehicle: Control vs Low | | $p$ <.05 * |
| | | Vehicle: Control vs High | | $p$ =.054 † |
| | | Vehicle: Low vs High | | $p$ =1 n.s. |
| | | C21: Control vs Low | | $p$ <.01 ** |
| | | C21: Control vs High | | $p$ =.268 n.s. |
| | | C21: Low vs High | | $p$ =.206 n.s. |
| 5L | Two-way ANOVA | Group | $F_{2,20}$ =4.345 | $p$ <.05 * |
| | | Condition | $F_{1,20}$ =1.818 | $p$ =.193 n.s. |
| | | Group: Condition | $F_{2,20}$ =1.339 | $p$ =.285 n.s. |
| | Pairwise t-test (Group) | Control: Vehicle vs C21 | | $p$ =.119 n.s. |
| | | Low: Vehicle vs C21 | | $p$ =.953 n.s. |
| | | High: Vehicle vs C21 | | $p$ =.314 n.s. |
| | (Condition) | Vehicle: Control vs Low | | $p$ =1 n.s. |
| | | Vehicle: Control vs High | | $p$ =.44 n.s. |
| | | Vehicle: Low vs High | | $p$ =.681 n.s. |
| | | C21: Control vs Low | | $p$ =.211 n.s. |
| | | C21: Control vs High | | $p$ <.05 * |
| | | C21: Low vs High | | $p$ =1 n.s. |
| 5M | Two-way ANOVA | Group | $F_{2,20}$ =3.886 | $p$ <.05 * |
| | | Condition | $F_{1,20}$ =1.868 | $p$ =.187 n.s. |
| | | Group: Condition | $F_{2,20}$ =1.384 | $p$ =.274 n.s. |
| | Pairwise t-test (Group) | Control: Vehicle vs C21 | | $p$ =.545 n.s. |
| | | Low: Vehicle vs C21 | | $p$ =.918 n.s. |
| | | High: Vehicle vs C21 | | $p$ =.204 n.s. |
| | (Condition) | Vehicle: Control vs Low | | $p$ =.757 n.s. |

| | | | | | |
|---|---|---|---|---|---|
| | | Vehicle: Control vs High | | | $p$ =.078 † |
| | | Vehicle: Low vs High | | | $p$ =.972 n.s. |
| | | C21: Control vs Low | | | $p$ =1 n.s. |
| | | C21: Control vs High | | | $p$ <.05 * |
| | | C21: Low vs High | | | $p$ =.337 n.s. |
| 5N | Type III Likelihood Ratio Chi-Square Test | Group | $X^2_{2}$ | =2587.6 | $p$ <.001 *** |
| | | Condition | $X^2_{1}$ | =1120.3 | $p$ <.001 *** |
| | | Group: Condition | $X^2_{2}$ | =1303.9 | $p$ <.001 *** |
| | Tukey-adjusted EMM comparisons (Group) | Control: Neutral vs Distressed | | | $p$ <.001 *** |
| | | Low: Neutral vs Distressed | | | $p$ <.01 ** |
| | | High: Neutral vs Distressed | | | $p$ <.001 *** |
| | (Condition) | Vehicle: Control vs Low | | | $p$ <.001 *** |
| | | Vehicle: Control vs High | | | $p$ <.001 *** |
| | | Vehicle: Low vs High | | | $p$ <.001 *** |
| | | C21: Control vs Low | | | $p$ <.001 *** |
| | | C21: Control vs High | | | $p$ <.001 *** |
| | | C21: Low vs High | | | $p$ <.001 *** |
| 5O | Two-way ANOVA | Group | $F_{2,20}$ | =4.696 | $p$ <.05 * |
| | | Condition | $F_{1,20}$ | =31.661 | $p$ <.001 *** |
| | | Group: Condition | $F_{2,20}$ | =7.639 | $p$ <.01 ** |
| | Pairwise t-test (Group) | Control: Vehicle vs C21 | | | $p$ <.01 ** |
| | | Low: Vehicle vs C21 | | | $p$ =.676 n.s. |
| | | High: Vehicle vs C21 | | | $p$ <.01 ** |
| | (Condition) | Vehicle: Control vs Low | | | $p$ =.739 n.s. |
| | | Vehicle: Control vs High | | | $p$ =.081 † |
| | | Vehicle: Low vs High | | | $p$ =.839 n.s. |
| | | C21: Control vs Low | | | $p$ <.01 ** |
| | | C21: Control vs High | | | $p$ =1 n.s. |
| | | C21: Low vs High | | | $p$ <.05 * |
| 5P | Two-way ANOVA | Group | $F_{2,20}$ | =3.474 | $p$ =.051 † |
| | | Condition | $F_{1,20}$ | =0.199 | $p$ =.660 n.s. |
| | | Group: Condition | $F_{2,20}$ | =0.885 | $p$ =.428 n.s. |
| 5Q | Two-way ANOVA | Group | $F_{2,20}$ | =2.249 | $p$ =.132 n.s. |
| | | Condition | $F_{1,20}$ | =0.099 | $p$ =.756 n.s. |
| | | Group: Condition | $F_{2,20}$ | =0.399 | $p$ =.676 n.s. |

**Table S5. Detailed statistical results corresponding to Figure 5.**

| Figure | Analysis | Factor | Statistical value | p-value |
|---|---|---|---|---|
| S1A | Two-way ANOVA | Group | $F_{2,31}$ = 48.056 | $p$ <.001 *** |
| | | Postnatal day (PND) | $F_{27,837}$ = 438.843 | $p$ <.001 *** |
| | | Group: PND | $F_{54,837}$ = 78.906 | $p$ <.001 *** |
| S1C | One-way ANOVA | Group | $F_{2,22}$ = 6.176 | $p$ <.01 ** |
| | Pairwise t-test | Control vs Low | | $p$ =.785 n.s. |
| | | Control vs High | | $p$ <.01 ** |
| | | Low vs High | | $p$ =.103 n.s. |
| S1D | One-way ANOVA | Group | $F_{2,22}$ = 4.164 | $p$ <.05 * |
| | Pairwise t-test | Control vs Low | | $p$ =.28 n.s. |
| | | Control vs High | | $p$ <.05 * |
| | | Low vs High | | $p$ =.898 n.s. |
| S1E | One-way ANOVA | Group | $F_{2,22}$ = 9.265 | $p$ <.01 ** |
| | Pairwise t-test | Control vs Low | | $p$ =.071 † |
| | | Control vs High | | $p$ <.001 *** |
| | | Low vs High | | $p$ =.259 n.s. |
| S1F | One-way ANOVA | Group | $F_{2,22}$ = 5.562 | $p$ <.05 * |
| | Pairwise t-test | Control vs Low | | $p$ =.261 n.s. |
| | | Control vs High | | $p$ <.01 ** |
| | | Low vs High | | $p$ =.45 n.s. |
| S1G | One-way ANOVA | Group | $F_{2,27}$ = 4.805 | $p$ <.05 * |
| | Pairwise t-test | Control vs Low | | $p$ =.782 n.s. |
| | | Control vs High | | $p$ <.05 * |
| | | Low vs High | | $p$ =.197 n.s. |
| S1H | One-way ANOVA | Group | $F_{2,27}$ = 2.891 | $p$ =.073 † |
| | Pairwise t-test | Control vs Low | | $p$ =.577 n.s. |
| | | Control vs High | | $p$ =.071 † |
| | | Low vs High | | $p$ =.893 n.s. |
| S1I | One-way ANOVA | Group | $F_{2,27}$ = 3.56 | $p$ <.05 * |
| | Pairwise t-test | Control vs Low | | $p$ =.617 n.s. |
| | | Control vs High | | $p$ <.05 * |
| | | Low vs High | | $p$ =.545 n.s. |
| S1J | One-way ANOVA | Group | $F_{2,27}$ = 0.881 | $p$ =.426 n.s. |
| S1L | One-way ANOVA | Group | $F_{2,22}$ = 1.057 | $p$ =.364 n.s. |
| S1M | One-way ANOVA | Group | $F_{2,22}$ = 3.276 | $p$ =.057 † |
| | Pairwise t-test | Control vs Low | | $p$ =.080 † |
| | | Control vs High | | $p$ =1 n.s. |
| | | Low vs High | | $p$ =.161 n.s. |
| S1N | One-way ANOVA | Group | $F_{2,22}$ = 0.793 | $p$ =.465 n.s. |

| | | | | |
|---|---|---|---|---|
| S1O | One-way ANOVA | Group | $F_{2,27}$ = 0.599 | $p$ = .556 n.s. |
| S1P | One-way ANOVA | Group | $F_{2,27}$ = 0.754 | $p$ = .48 n.s. |
| S1Q | One-way ANOVA | Group | $F_{2,27}$ = 0.242 | $p$ = .787 n.s. |

**Table S6. Detailed statistical results corresponding to Figure S1.**

| Figure | Analysis | Factor | Statistical value | p-value |
|---|---|---|---|---|
| S2A | One-way ANOVA | Group | $F_{2,22} = 1.048$ | $p = .367$ n.s. |
| S2B | One-way ANOVA | Group | $F_{2,22} = 0.241$ | $p = .788$ n.s. |
| S2C | One-way ANOVA | Group | $F_{2,22} = 0.341$ | $p = .715$ n.s. |
| S2D | Pearson's product-moment | | $t_{23} = -1.784$ | $p = .088$ † |
| | correlation coefficient | | $r = -0.349$ | |
| S2E | One-way ANOVA | Group | $F_{2,24} = 0.703$ | $p = .505$ n.s. |
| S2F | One-way ANOVA | Group | $F_{2,24} = 1.882$ | $p = .174$ n.s. |
| S2G | One-way ANOVA | Group | $F_{2,24} = 0.316$ | $p = .732$ n.s. |
| S2H | Pearson's product-moment | | $t_{25} = -1.606$ | $p = .121$ n.s. |
| | correlation coefficient | | $r = -0.306$ | |
| S2J | One-way ANOVA | Group | $F_{2,22} = 0.06$ | $p = .942$ n.s. |
| S2K | Two-way ANOVA | Group | $F_{2,22} = 11.413$ | $p < .001$ *** |
| | | Demonstrator | $F_{1,22} = 21.900$ | $p < .001$ *** |
| | | Group: Demonstrator | $F_{2,22} = 4.614$ | $p < .05$ * |
| | Pairwise t-test | Control: Neutral vs Distressed | | $p < .01$ ** |
| | (Group) | Low: Neutral vs Distressed | | $p < .05$ * |
| | | High: Neutral vs Distressed | | $p = 1$ n.s. |
| | (Demonstrator) | Neutral: Control vs Low | | $p < .01$ ** |
| | | Neutral: Control vs High | | $p < .05$ * |
| | | Neutral: Low vs High | | $p = 1$ n.s. |
| | | Distressed: Control vs Low | | $p < .01$ ** |
| | | Distressed: Control vs High | | $p < .001$ *** |
| | | Distressed: Low vs High | | $p = 1$ n.s. |
| S2L | One-way ANOVA | Group | $F_{2,24} = 0.887$ | $p = .425$ n.s. |
| S2M | Two-way ANOVA | Group | $F_{2,22} = 11.413$ | $p < .001$ *** |
| | | Demonstrator | $F_{1,22} = 21.900$ | $p < .001$ *** |
| | | Group: Demonstrator | $F_{2,22} = 4.614$ | $p < .05$ * |
| | Pairwise t-test | Control: Neutral vs Distressed | | $p = .074$ † |
| | (Group) | Low: Neutral vs Distressed | | $p < .05$ * |
| | | High: Neutral vs Distressed | | $p < .01$ ** |
| | (Demonstrator) | Neutral: Control vs Low | | $p < .05$ * |
| | | Neutral: Control vs High | | $p < .05$ * |
| | | Neutral: Low vs High | | $p = 1$ n.s. |
| | | Distressed: Control vs Low | | $p < .01$ ** |
| | | Distressed: Control vs High | | $p < .01$ ** |
| | | Distressed: Low vs High | | $p = 1$ n.s. |
| S2N | Two-way ANOVA | Group | $F_{2,22} = 17.934$ | $p < .001$ *** |
| | | Demonstrator | $F_{1,22} = 34.390$ | $p < .001$ *** |
| | | Group: Demonstrator | $F_{2,22} = 13.884$ | $p < .001$ *** |

| | | | | |
|---|---|---|---|---|
| | Pairwise t-test | Control: Neutral vs Distressed | | $p < .001$ *** |
| | (Group) | Low: Neutral vs Distressed | | $p = .457$ n.s. |
| | | High: Neutral vs Distressed | | $p = .051$ † |
| | (Demonstrator) | Neutral: Control vs Low | | $p = .609$ n.s. |
| | | Neutral: Control vs High | | $p = .125$ n.s. |
| | | Neutral: Low vs High | | $p < .01$ ** |
| | | Distressed: Control vs Low | | $p < .001$ *** |
| | | Distressed: Control vs High | | $p < .001$ *** |
| | | Distressed: Low vs High | | $p = 1$ n.s. |
| S2O | Two-way ANOVA | Group | $F_{2,22} = 16.885$ | $p < .001$ *** |
| | | Demonstrator | $F_{1,22} = 22.719$ | $p < .001$ *** |
| | | Group: Demonstrator | $F_{2,22} = 0.607$ | $p = .554$ n.s. |
| | Pairwise t-test | Control: Neutral vs Distressed | | $p < .05$ * |
| | (Group) | Low: Neutral vs Distressed | | $p < .05$ * |
| | | High: Neutral vs Distressed | | $p < .01$ ** |
| | (Demonstrator) | Neutral: Control vs Low | | $p < .001$ *** |
| | | Neutral: Control vs High | | $p < .001$ *** |
| | | Neutral: Low vs High | | $p = 1$ n.s. |
| | | Distressed: Control vs Low | | $p < .01$ ** |
| | | Distressed: Control vs High | | $p < .05$ * |
| | | Distressed: Low vs High | | $p = 1$ n.s. |
| S2P | Two-way ANOVA | Group | $F_{2,22} = 2.219$ | $p = .132$ n.s. |
| | | Demonstrator | $F_{1,22} = 0.010$ | $p = .922$ n.s. |
| | | Group: Demonstrator | $F_{2,22} = 0.218$ | $p = .806$ n.s. |
| S2Q | Two-way ANOVA | Group | $F_{2,25} = 6.200$ | $p < .001$ *** |
| | | Demonstrator | $F_{1,25} = 15.964$ | $p < .001$ *** |
| | | Group: Demonstrator | $F_{2,25} = 6.959$ | $p < .001$ *** |
| | Pairwise t-test | Control: Neutral vs Distressed | | $p < .05$ * |
| | (Group) | Low: Neutral vs Distressed | | $p = .366$ n.s. |
| | | High: Neutral vs Distressed | | $p = .418$ n.s. |
| | (Demonstrator) | Neutral: Control vs Low | | $p = .947$ n.s. |
| | | Neutral: Control vs High | | $p = .56$ n.s. |
| | | Neutral: Low vs High | | $p = 1$ n.s. |
| | | Distressed: Control vs Low | | $p < .01$ ** |
| | | Distressed: Control vs High | | $p < .01$ ** |
| | | Distressed: Low vs High | | $p = 1$ n.s. |
| S2R | Two-way ANOVA | Group | $F_{2,25} = 3.254$ | $p = .055$ † |
| | | Demonstrator | $F_{1,25} = 27.470$ | $p < .001$ *** |
| | | Group: Demonstrator | $F_{2,25} = 4.936$ | $p < .05$ * |
| | Pairwise t-test | Control: Neutral vs Distressed | | $p < .01$ ** |

| | | | | | |
|---|---|---|---|---|---|
| | (Group) | Low: Neutral vs Distressed | | $p < .01$ | ** |
| | | High: Neutral vs Distressed | | $p = .232$ | n.s. |
| | (Demonstrator) | Neutral: Control vs Low | | $p = 1$ | n.s. |
| | | Neutral: Control vs High | | $p = 1$ | n.s. |
| | | Neutral: Low vs High | | $p = 1$ | n.s. |
| | | Distressed: Control vs Low | | $p = .499$ | n.s. |
| | | Distressed: Control vs High | | $p < .05$ | * |
| | | Distressed: Low vs High | | $p = .243$ | n.s. |
| S2S | Two-way ANOVA | Group | $F_{2,25} = 1.766$ | $p = .193$ | n.s. |
| | | Demonstrator | $F_{1,25} = 1.614$ | $p = .217$ | n.s. |
| | | Group: Demonstrator | $F_{2,25} = 0.538$ | $p = .591$ | n.s. |

**Table S7. Detailed statistical results corresponding to Figure S2.**

| Figure | Analysis | Factor | Statistical value | p-value |
|---|---|---|---|---|
| S3B | One-sample t-test | Control vs chance level | $t_{8} = 6.131$ | $p < .001$ *** |
| | | Low vs chance level | $t_{7} = 8.147$ | $p < .001$ *** |
| | | High vs chance level | $t_{7} = -1.774$ | $p = .119$ n.s. |
| | Pairwise t-test | Control vs Low | | $p = .266$ n.s. |
| | | Control vs High | | $p < .001$ *** |
| | | Low vs High | | $p < .001$ *** |
| S3C | Two-way ANOVA | Group | $F_{2,22} = 2.492$ | $p = .106$ n.s. |
| | | Demonstrator | $F_{1,22} = 37.786$ | $p < .001$ *** |
| | | Group: Demonstrator | $F_{2,22} = 18.892$ | $p < .001$ *** |
| | Pairwise t-test | Control: Neutral vs Distressed | | $p < .001$ *** |
| | (Group) | Low: Neutral vs Distressed | | $p < .01$ ** |
| | | High: Neutral vs Distressed | | $p = .121$ n.s. |
| | (Demonstrator) | Neutral: Control vs Low | | $p = 1$ n.s. |
| | | Neutral: Control vs High | | $p = 1$ n.s. |
| | | Neutral: Low vs High | | $p = 1$ n.s. |
| | | Distressed: Control vs Low | | $p = .272$ n.s. |
| | | Distressed: Control vs High | | $p < .01$ ** |
| | | Distressed: Low vs High | | $p = .102$ n.s. |
| S3D | One-way ANOVA | Group | $F_{2,22} = 0.834$ | $p = .448$ n.s. |
| S3E | Two-way ANOVA | Group | $F_{2,22} = 7.576$ | $p < .01$ ** |
| | | Demonstrator | $F_{1,22} = 4.573$ | $p < .05$ * |
| | | Group: Demonstrator | $F_{2,22} = 3.715$ | $p < .05$ * |
| | Pairwise t-test | Control: Neutral vs Distressed | | $p = .054$ † |
| | (Group) | Low: Neutral vs Distressed | | $p = .13$ n.s. |
| | | High: Neutral vs Distressed | | $p = .244$ n.s. |
| | (Demonstrator) | Neutral: Control vs Low | | $p = 1$ n.s. |
| | | Neutral: Control vs High | | $p = .369$ n.s. |
| | | Neutral: Low vs High | | $p = 1$ n.s. |
| | | Distressed: Control vs Low | | $p < .05$ * |
| | | Distressed: Control vs High | | $p < .001$ *** |
| | | Distressed: Low vs High | | $p = .641$ n.s. |
| S3F | One-sample t-test | Control vs chance level | $t_{7} = 3.639$ | $p < .01$ ** |
| | | Low vs chance level | $t_{9} = 10.715$ | $p < .001$ *** |
| | | High vs chance level | $t_{8} = -0.459$ | $p = .658$ n.s. |
| | Pairwise t-test | Control vs Low | | $p = 1$ n.s. |
| | | Control vs High | | $p < .01$ ** |
| | | Low vs High | | $p < .01$ ** |
| S3G | Two-way ANOVA | Group | $F_{2,24} = 2.858$ | $p = .077$ † |
| | | Demonstrator | $F_{1,24} = 16.789$ | $p < .001$ *** |

| | | | | |
|---|---|---|---|---|
| | | Group: Demonstrator | $F_{2,24}$ = 6.219 | *p* <.001 *** |
| | Pairwise t-test | Control: Neutral vs Distressed | | *p* <.05 * |
| | (Group) | Low: Neutral vs Distressed | | *p* <.001 *** |
| | | High: Neutral vs Distressed | | *p* =.397 n.s. |
| | (Demonstrator) | Neutral: Control vs Low | | *p* =1 n.s. |
| | | Neutral: Control vs High | | *p* =1 n.s. |
| | | Neutral: Low vs High | | *p* =1 n.s. |
| | | Distressed: Control vs Low | | *p* =.36 n.s. |
| | | Distressed: Control vs High | | *p* <.05 * |
| | | Distressed: Low vs High | | *p* =.404 n.s. |
| S3H | One-way ANOVA | Group | $F_{2,24}$ = 1.961 | *p* =.163 n.s. |
| S3I | Two-way ANOVA | Group | $F_{2,24}$ = 4.198 | *p* <.05 * |
| | | Demonstrator | $F_{1,24}$ = 17.793 | *p* <.001 *** |
| | | Group: Demonstrator | $F_{2,24}$ = 4.011 | *p* <.05 * |
| | Pairwise t-test | Control: Neutral vs Distressed | | *p* <.05 * |
| | (Group) | Low: Neutral vs Distressed | | *p* <.01 ** |
| | | High: Neutral vs Distressed | | *p* =.865 n.s. |
| | (Demonstrator) | Neutral: Control vs Low | | *p* =1 n.s. |
| | | Neutral: Control vs High | | *p* =.238 n.s. |
| | | Neutral: Low vs High | | *p* =.121 n.s. |
| | | Distressed: Control vs Low | | *p* =.196 n.s. |
| | | Distressed: Control vs High | | *p* <.05 * |
| | | Distressed: Low vs High | | *p* =.514 n.s. |

**Table S8. Detailed statistical results corresponding to Figure S3.**

| Figure | Analysis | Factor | Statistical value | p-value |
|---|---|---|---|---|
| S4B | One-sample t-test | Control vs chance level | $t_{5} = 12.887$ | $p < .001$ *** |
| | | High vs chance level | $t_{7} = -2.81$ | $p < .05$ * |
| | One-way ANOVA | Group | $F_{1,10} = 83.17$ | $p < .001$ *** |
| | Pairwise t-test | Control vs High | | $p < .001$ *** |
| S4C | Two-way ANOVA | Group | $F_{1,10} = 8.989$ | $p < .05$ * |
| | | Demonstrator | $F_{1,10} = 24.965$ | $p < .001$ *** |
| | | Group: Demonstrator | $F_{1,10} = 76.909$ | $p < .001$ *** |
| | Pairwise t-test | Control: Neutral vs Stressed | | $p < .001$ *** |
| | (Group) | High: Neutral vs Stressed | | $p < .05$ * |
| | (Demonstrator) | Neutral: Control vs High | | $p = .062$ † |
| | | Stressed: Control vs High | | $p < .001$ *** |
| S4D | One-way ANOVA | Group | $F_{1,10} = 45.43$ | $p < .001$ *** |
| | Pairwise t-test | Control vs High | | $p < .001$ *** |
| S4E | One-way ANOVA | Group | $F_{1,10} = 0.506$ | $p = .493$ n.s. |
| S4F | Two-way ANOVA | Group | $F_{1,10} = 33.571$ | $p < .001$ *** |
| | | Demonstrator | $F_{1,10} = 18.175$ | $p < .01$ ** |
| | | Group: Demonstrator | $F_{1,10} = 38.591$ | $p < .001$ *** |
| | Pairwise t-test | Control: Neutral vs Stressed | | $p < .01$ ** |
| | (Group) | High: Neutral vs Stressed | | $p = .175$ n.s. |
| | (Demonstrator) | Neutral: Control vs High | | $p = .763$ n.s. |
| | | Stressed: Control vs High | | $p < .001$ *** |

**Table S9. Detailed statistical results corresponding to Figure S4.**

| Figure | Analysis | Factor | Statistical value | p-value |
|---|---|---|---|---|
| S5B | One-way ANOVA (Male) | Demonstrator | $F_{1,7}$ = 0.809 | $p$ = .398 n.s. |
| | One-way ANOVA (Female) | Demonstrator | $F_{1,9}$ = 4.549 | $p$ = .062 n.s. |
| S5C | One-way ANOVA (Male) | Demonstrator | $F_{1,7}$ = 12.145 | $p$ < .05 * |
| | Pairwise t-test | Novel vs Familiar | | $p$ < .05 * |
| | One-way ANOVA (Female) | Demonstrator | $F_{1,9}$ = 29.102 | $p$ < .001 *** |
| | Pairwise t-test | Novel vs Familiar | | $p$ < .001 *** |
| S5D | One-sample t-test | Male vs chance level | $t_{7}$ = 4.214 | $p$ < .01 ** |
| | | Female vs chance level | $t_{9}$ = 5.702 | $p$ < .001 *** |
| S5E | One-sample t-test | Control vs chance level | $t_{9}$ = 11.291 | $p$ < .001 *** |
| | | Low vs chance level | $t_{9}$ = 2.874 | $p$ < .05 * |
| | | High vs chance level | $t_{11}$ = 6.026 | $p$ < .001 *** |
| | One-way ANOVA | Group | $F_{2,31}$ = 0.599 | $p$ = .555 n.s. |
| S5G | Two-way ANOVA | Group | $F_{2,31}$ = 8.017 | $p$ < .01 ** |
| | | Demonstrator | $F_{1,31}$ = 65.491 | $p$ < .001 *** |
| | | Group: Demonstrator | $F_{2,31}$ = 3.270 | $p$ = .051 † |
| | Pairwise t-test | Control: Neutral vs Distressed | | $p$ < .001 *** |
| | (Group) | Low: Neutral vs Distressed | | $p$ < .05 * |
| | | High: Neutral vs Distressed | | $p$ < .001 *** |
| | (Demonstrator) | Neutral: Control vs Low | | $p$ = .141 n.s. |
| | | Neutral: Control vs High | | $p$ < .05 * |
| | | Neutral: Low vs High | | $p$ = .806 n.s. |
| | | Distressed: Control vs Low | | $p$ < .01 ** |
| | | Distressed: Control vs High | | $p$ < .01 ** |
| | | Distressed: Low vs High | | $p$ = 1 n.s. |

**Table S10. Detailed statistical results corresponding to Figure S5.**

| Figure | Analysis | Factor | Statistical value | p-value |
|---|---|---|---|---|
| S6A | Two-way ANOVA | Group | $F_{2,26} = 1.609$ | $p = .219$ n.s. |
| | | Demonstrator | $F_{1,26} = 0.002$ | $p = .963$ n.s. |
| | | Group: Demonstrator | $F_{2,26} = 0.899$ | $p = .419$ n.s. |
| S6B | Three-way ANOVA | Group | $F_{2,26} = 6.721$ | $p < .01$ ** |
| | | Demonstrator | $F_{1,26} = 13.054$ | $p < .01$ ** |
| | | Condition | $F_{1,26} = 0.029$ | $p = .865$ n.s. |
| | | Group: Demonstrator | $F_{2,26} = 0.567$ | $p = .574$ n.s. |
| | | Group: Condition | $F_{2,26} = 0.360$ | $p = .701$ n.s. |
| | | Demonstrator: Condition | $F_{1,26} = 2.175$ | $p = .152$ n.s. |
| | | Group: Demonstrator: Condition | $F_{2,26} = 5.055$ | $p < .05$ * |
| | Two-way ANOVA | Demonstrator | $F_{1,8} = 4.61$ | $p = .064$ † |
| | (Group: Control) | Condition | $F_{1,8} = 0.471$ | $p = .512$ n.s. |
| | | Demonstrator: Condition | $F_{1,8} = 4.94$ | $p = .057$ † |
| | (Group: Low) | Demonstrator | $F_{1,9} = 3.70$ | $p = .086$ † |
| | | Condition | $F_{1,9} = 0.01$ | $p = .924$ n.s. |
| | | Demonstrator: Condition | $F_{1,9} = 4.11$ | $p = .073$ † |
| | (Group: High) | Demonstrator | $F_{1,9} = 4.80$ | $p = .056$ † |
| | | Condition | $F_{1,9} = 0.162$ | $p = .696$ n.s. |
| | | Demonstrator: Condition | $F_{1,9} = 2.57$ | $p = .053$ † |
| | Two-way ANOVA | Group | $F_{2,26} = 6.96$ | $p < .01$ ** |
| | (Condition: Vehicle) | Demonstrator | $F_{1,26} = 10.9$ | $p < .01$ ** |
| | | Group: Demonstrator | $F_{2,26} = 3.30$ | $p < .05$ * |
| | (Condition: C21) | Group | $F_{2,26} = 1.8$ | $p = .185$ n.s. |
| | | Demonstrator | $F_{1,26} = 1.73$ | $p = .2$ n.s. |
| | | Group: Demonstrator | $F_{2,26} = 2.94$ | $p = .07$ † |
| | Two-way ANOVA | Group | $F_{2,26} = 5.63$ | $p < .01$ ** |
| | (Demonstrator: Neutral) | Condition | $F_{1,26} = 1.29$ | $p = .267$ n.s. |
| | | Group: Condition | $F_{2,26} = 2.21$ | $p = .13$ n.s. |
| | (Demonstrator:Distressed) | Group | $F_{2,26} = 4.49$ | $p < .05$ * |
| | | Condition | $F_{1,26} = 0.769$ | $p = .388$ n.s. |
| | | Group: Condition | $F_{2,26} = 2.28$ | $p = .122$ n.s. |
| | Pairwise t-test | Vehicle: Neutral vs Distressed | | $p < .05$ * |
| | (Group: Control) | C21: Neutral vs Distressed | | $p = .813$ n.s. |
| | | Neutral: Vehicle vs C21 | | $p = .174$ n.s. |
| | | Distressed: Vehicle vs C21 | | $p = .153$ n.s. |
| | (Group: Low) | Vehicle: Neutral vs Distressed | | $p < .05$ * |
| | | C21: Neutral vs Distressed | | $p = .748$ n.s. |
| | | Neutral: Vehicle vs C21 | | $p = .296$ n.s. |
| | | Distressed: Vehicle vs C21 | | $p = .262$ n.s. |

| | | | | |
|---|---|---|---|---|
| | (Group: High) | Vehicle: Neutral vs Distressed | | $p$ =.903 n.s. |
| | | C21: Neutral vs Distressed | | $p$ =.068 † |
| | | Neutral: Vehicle vs C21 | | $p$ =.057 † |
| | | Distressed: Vehicle vs C21 | | $p$ =.35 n.s. |
| | Pairwise t-test | Neutral: Control vs Low | | $p$ =.093 † |
| | (Condition: Vehicle) | Neutral: Control vs High | | $p$ =.973 n.s. |
| | | Neutral: High vs Low | | $p$ =.603 n.s. |
| | | Distressed: Control vs Low | | $p$ <.05 * |
| | | Distressed: Control vs High | | $p$ <.01 ** |
| | | Distressed: High vs Low | | $p$ =1 n.s. |
| | (Condition: C21) | Neutral: Control vs Low | | $p$ =.199 n.s. |
| | | Neutral: Control vs High | | $p$ <.05 * |
| | | Neutral: High vs Low | | $p$ =.73 n.s. |
| | | Distressed: Control vs Low | | $p$ =1 n.s. |
| | | Distressed: Control vs High | | $p$ =.634 n.s. |
| | | Distressed: High vs Low | | $p$ =.773 n.s. |
| S6C | Two-way ANOVA | Group | $F_{2,21}$ =0.104 | $p$ =.902 n.s. |
| | | Condition | $F_{1,21}$ =3.769 | $p$ =.066 † |
| | | Group: Condition | $F_{2,21}$ =1.405 | $p$ =.268 n.s. |
| | Pairwise t-test | Control: Vehicle vs C21 | | $p$ <.05 * |
| | (Group) | Low: Vehicle vs C21 | | $p$ =.176 n.s. |
| | | High: Vehicle vs C21 | | $p$ =.918 n.s. |
| | (Condition) | Vehicle: Control vs Low | | $p$ =1 n.s. |
| | | Vehicle: Control vs High | | $p$ =1 n.s. |
| | | Vehicle: Low vs High | | $p$ =1 n.s. |
| | | C21: Control vs Low | | $p$ =1 n.s. |
| | | C21: Control vs High | | $p$ =.699 n.s. |
| | | C21: Low vs High | | $p$ =1 n.s. |
| S6D | Three-way ANOVA | Group | $F_{2,21}$ =2.937 | $p$ =.075 † |
| | | Demonstrator | $F_{1,21}$ =24.527 | $p$ <.001 *** |
| | | Condition | $F_{1,21}$ =0.285 | $p$ =.599 n.s. |
| | | Group: Demonstrator | $F_{2,21}$ =0.067 | $p$ =.935 n.s. |
| | | Group: Condition | $F_{2,21}$ =1.394 | $p$ =.270 n.s. |
| | | Demonstrator: Condition | $F_{1,21}$ =0.439 | $p$ =.515 n.s. |
| | | Group: Demonstrator: Condition | $F_{2,21}$ =8.765 | $p$ <.01 ** |
| | Two-way ANOVA | Demonstrator | $F_{1,7}$ =10.1 | $p$ <.05 * |
| | (Group: Control) | Condition | $F_{1,7}$ =0.294 | $p$ =.604 n.s. |
| | | Demonstrator: Condition | $F_{1,7}$ =12.9 | $p$ <.01 ** |
| | (Group: Low) | Demonstrator | $F_{1,6}$ =4.94 | $p$ =.068 † |
| | | Condition | $F_{1,6}$ =1.38 | $p$ =.285 n.s. |

| | | | |
|---|---|---|---|
| | Demonstrator: Condition | $F_{1,6} = 0.087$ | $p = .778$ n.s. |
| (Group: High) | Demonstrator | $F_{1,8} = 11.5$ | $p < .05$ * |
| | Condition | $F_{1,8} = 1.33$ | $p = .282$ n.s. |
| | Demonstrator: Condition | $F_{1,8} = 5.79$ | $p < .05$ * |
| Two-way ANOVA | Group | $F_{2,21} = 3.69$ | $p < .05$ * |
| (Condition: Vehicle) | Demonstrator | $F_{1,21} = 14.5$ | $p < .01$ ** |
| | Group: Demonstrator | $F_{2,21} = 5.47$ | $p < .05$ * |
| (Condition: C21) | Group | $F_{2,21} = 1.35$ | $p = .3$ n.s. |
| | Demonstrator | $F_{1,21} = 5.63$ | $p < .05$ * |
| | Group: Demonstrator | $F_{2,21} = 5.51$ | $p < .05$ * |
| Two-way ANOVA | Group | $F_{2,21} = 2.21$ | $p = .135$ n.s. |
| (Demonstrator: Neutral) | Condition | $F_{1,21} = 0.008$ | $p = .931$ n.s. |
| | Group: Condition | $F_{2,21} = 3.83$ | $p < .05$ * |
| (Demonstrator:Distressed) | Group | $F_{2,21} = 2.38$ | $p = .117$ n.s. |
| | Condition | $F_{1,21} = 0.53$ | $p = .474$ n.s. |
| | Group: Condition | $F_{2,21} = 5.44$ | $p < .05$ * |
| Pairwise t-test | Vehicle: Neutral vs Distressed | | $p < .001$ *** |
| (Group: Control) | C21: Neutral vs Distressed | | $p = .43$ n.s. |
| | Neutral: Vehicle vs C21 | | $p = .101$ n.s. |
| | Distressed: Vehicle vs C21 | | $p < .05$ * |
| (Group: Low) | Vehicle: Neutral vs Distressed | | $p = .285$ n.s. |
| | C21: Neutral vs Distressed | | $p = .098$ † |
| | Neutral: Vehicle vs C21 | | $p = .251$ n.s. |
| | Distressed: Vehicle vs C21 | | $p = .443$ n.s. |
| (Group: High) | Vehicle: Neutral vs Distressed | | $p = 1$ n.s. |
| | C21: Neutral vs Distressed | | $p < .01$ ** |
| | Neutral: Vehicle vs C21 | | $p = .231$ n.s. |
| | Distressed: Vehicle vs C21 | | $p = .056$ † |
| Pairwise t-test | Neutral: Control vs Low | | $p = 1$ n.s. |
| (Condition: Vehicle) | Neutral: Control vs High | | $p = 1$ n.s. |
| | Neutral: High vs Low | | $p = 1$ n.s. |
| | Distressed: Control vs Low | | $p = .224$ n.s. |
| | Distressed: Control vs High | | $p < .01$ ** |
| | Distressed: High vs Low | | $p = .176$ n.s. |
| (Condition: C21) | Neutral: Control vs Low | | $p = .097$ † |
| | Neutral: Control vs High | | $p < .05$ * |
| | Neutral: High vs Low | | $p = 1$ n.s. |
| | Distressed: Control vs Low | | $p = 1$ n.s. |
| | Distressed: Control vs High | | $p = 1$ n.s. |
| | Distressed: High vs Low | | $p = .627$ n.s. |

| | | | | | |
|---|---|---|---|---|---|
| S6E | Two-way ANOVA | Group | $F_{2,21} = 31.702$ | $p < .001$ | *** |
| | | Condition | $F_{1,21} = 0.078$ | $p = .783$ | n.s. |
| | | Group: Condition | $F_{2,21} = 11.065$ | $p < .001$ | *** |
| | Pairwise t-test | Control: Vehicle vs C21 | | $p < .05$ | * |
| | (Group) | Low: Vehicle vs C21 | | $p = .064$ | † |
| | | High: Vehicle vs C21 | | $p < .01$ | ** |
| | (Condition) | Vehicle: Control vs Low | | $p < .001$ | *** |
| | | Vehicle: Control vs High | | $p < .001$ | *** |
| | | Vehicle: Low vs High | | $p = 1$ | n.s. |
| | | C21: Control vs Low | | $p = .135$ | n.s. |
| | | C21: Control vs High | | $p = .137$ | n.s. |
| | | C21: Low vs High | | $p = 1$ | n.s. |
| S6F | Two-way ANOVA | Group | $F_{2,21} = 1.403$ | $p = .268$ | n.s. |
| | | Condition | $F_{1,21} = 4.210$ | $p = .053$ | † |
| | | Group: Condition | $F_{2,21} = 3.411$ | $p = .052$ | † |
| | Pairwise t-test | Control: Vehicle vs C21 | | $p = .545$ | n.s. |
| | (Group) | Low: Vehicle vs C21 | | $p < .01$ | ** |
| | | High: Vehicle vs C21 | | $p < .05$ | * |
| | (Condition) | Vehicle: Control vs Low | | $p = .059$ | † |
| | | Vehicle: Control vs High | | $p < .05$ | * |
| | | Vehicle: Low vs High | | $p = .634$ | n.s. |
| | | C21: Control vs Low | | $p = 1$ | n.s. |
| | | C21: Control vs High | | $p = 1$ | n.s. |
| | | C21: Low vs High | | $p = 1$ | n.s. |
| S6G | Two-way ANOVA | Group | $F_{2,21} = 1.923$ | $p = .171$ | n.s. |
| | | Condition | $F_{1,21} = 1.189$ | $p = .288$ | n.s. |
| | | Group: Condition | $F_{2,21} = 5.609$ | $p < .05$ | * |
| | Pairwise t-test | Control: Vehicle vs C21 | | $p = .087$ | † |
| | (Group) | Low: Vehicle vs C21 | | $p = .075$ | † |
| | | High: Vehicle vs C21 | | $p = .066$ | † |
| | (Condition) | Vehicle: Control vs Low | | $p = 1$ | n.s. |
| | | Vehicle: Control vs High | | $p = 1$ | n.s. |
| | | Vehicle: Low vs High | | $p = 1$ | n.s. |
| | | C21: Control vs Low | | $p < .01$ | ** |
| | | C21: Control vs High | | $p < .01$ | ** |
| | | C21: Low vs High | | $p = 1$ | n.s. |
| S6H | Two-way ANOVA | Group | $F_{2,22} = 1.080$ | $p = .357$ | n.s. |
| | | Condition | $F_{1,22} = 5.272$ | $p < .05$ | * |
| | | Group: Condition | $F_{2,22} = 4.041$ | $p < .05$ | * |
| | Pairwise t-test | Control: Vehicle vs C21 | | $p = .465$ | n.s. |

| | | | | | |
|---|---|---|---|---|---|
| | (Group) | Low: Vehicle vs C21 | | $p$ =.169 | n.s. |
| | | High: Vehicle vs C21 | | $p$ <.01 | ** |
| | (Condition) | Vehicle: Control vs Low | | $p$ =.128 | n.s. |
| | | Vehicle: Control vs High | | $p$ <.05 | * |
| | | Vehicle: Low vs High | | $p$ =1 | n.s. |
| | | C21: Control vs Low | | $p$ =1 | n.s. |
| | | C21: Control vs High | | $p$ =1 | n.s. |
| | | C21: Low vs High | | $p$ =1 | n.s. |
| S6I | Two-way ANOVA | Group | $F_{2,22}$ =0.375 | $p$ =.692 | n.s. |
| | | Condition | $F_{1,22}$ =4.755 | $p$ <.05 | * |
| | | Group: Condition | $F_{2,22}$ =7.622 | $p$ <.01 | ** |
| | Pairwise t-test | Control: Vehicle vs C21 | | $p$ <.05 | * |
| | (Group) | Low: Vehicle vs C21 | | $p$ <.05 | * |
| | | High: Vehicle vs C21 | | $p$ <.05 | * |
| | (Condition) | Vehicle: Control vs Low | | $p$ =.309 | n.s. |
| | | Vehicle: Control vs High | | $p$ =.119 | n.s. |
| | | Vehicle: Low vs High | | $p$ =1 | n.s. |
| | | C21: Control vs Low | | $p$ =.252 | n.s. |
| | | C21: Control vs High | | $p$ =1 | n.s. |
| | | C21: Low vs High | | $p$ =1 | n.s. |
| S6J | Two-way ANOVA | Group | $F_{2,22}$ =2.179 | $p$ =.137 | n.s. |
| | | Condition | $F_{1,22}$ =3.473 | $p$ =.076 | † |
| | | Group: Condition | $F_{2,22}$ =1.518 | $p$ =.228 | n.s. |
| | Pairwise t-test | Control: Vehicle vs C21 | | $p$ =.094 | † |
| | (Group) | Low: Vehicle vs C21 | | $p$ =.799 | n.s. |
| | | High: Vehicle vs C21 | | $p$ =.076 | † |
| | (Condition) | Vehicle: Control vs Low | | $p$ =1 | n.s. |
| | | Vehicle: Control vs High | | $p$ =.939 | n.s. |
| | | Vehicle: Low vs High | | $p$ =.664 | n.s. |
| | | C21: Control vs Low | | $p$ =.23 | n.s. |
| | | C21: Control vs High | | $p$ =1 | n.s. |
| | | C21: Low vs High | | $p$ =.107 | n.s. |

**Table S11. Detailed statistical results corresponding to Figure S6.**

| Figure | Analysis | Factor | Statistical value | p-value |
|---|---|---|---|---|
| S7B | One-sample t-test | Control (Vehicle) vs chance level | $t_{14}$ = 6.343 | $p$ <.001 *** |
| | | Control (C21) vs chance level | $t_{14}$ = 5.165 | $p$ <.001 *** |
| | | High (Vehicle) vs chance level | $t_{11}$ = 0.323 | $p$ =.753 n.s. |
| | | High (C21) vs chance level | $t_{11}$ = 0.270 | $p$ =.792 n.s. |
| | Two-way ANOVA | Group | $F_{1,25}$ = 22.391 | $p$ <.001 *** |
| | | Condition | $F_{1,25}$ = 1.223 | $p$ =.279 n.s. |
| | | Group: Condition | $F_{1,25}$ = 1.045 | $p$ =.316 n.s. |
| | Pairwise t-test (Group) | Control: Vehicle vs C21 | | $p$ =.187 n.s. |
| | | High: Vehicle vs C21 | | $p$ =.943 n.s. |
| | (Condition) | Vehicle: Control vs High | | $p$ <.001 *** |
| | | C21: Control vs High | | $p$ <.01 ** |
| S7C | Three-way ANOVA | Group | $F_{1,25}$ = 4.541 | $p$ <.05 * |
| | | Demonstrator | $F_{1,25}$ = 16.044 | $p$ <.001 *** |
| | | Condition | $F_{1,25}$ = 1.161 | $p$ =.292 n.s. |
| | | Group: Demonstrator | $F_{1,25}$ = 16.304 | $p$ <.001 *** |
| | | Group: Condition | $F_{1,25}$ = 1.689 | $p$ =.206 n.s. |
| | | Demonstrator: Condition | $F_{1,25}$ = 2.924 | $p$ =.100 n.s. |
| | | Group: Demonstrator: Condition | $F_{1,25}$ = 3.403 | $p$ =.077 † |
| | Two-way ANOVA (Group: Control) | Demonstrator | $F_{1,14}$ = 2.97 | $p$ <.001 *** |
| | | Condition | $F_{1,14}$ = 24.5 | $p$ =.107 n.s. |
| | | Demonstrator: Condition | $F_{1,14}$ = 4.28 | $p$ =.058 † |
| | (Group: High) | Demonstrator | $F_{1,11}$ = 0.024 | $p$ =.974 n.s. |
| | | Condition | $F_{1,11}$ = 0.001 | $p$ =.879 n.s. |
| | | Demonstrator: Condition | $F_{1,11}$ = 0.051 | $p$ =.825 n.s. |
| | Two-way ANOVA (Condition: Vehicle) | Group | $F_{1,25}$ = 5.83 | $p$ <.05 * |
| | | Demonstrator | $F_{1,25}$ = 13.2 | $p$ <.01 ** |
| | | Group: Demonstrator | $F_{1,25}$ = 13.8 | $p$ <.01 ** |
| | (Condition: C21) | Group | $F_{1,25}$ = 2.54 | $p$ =.123 n.s. |
| | | Demonstrator | $F_{1,25}$ = 8.83 | $p$ <.01 ** |
| | | Group: Demonstrator | $F_{1,25}$ = 8.45 | $p$ <.01 ** |
| | Two-way ANOVA (Demonstrator: Neutral) | Group | $F_{1,25}$ = 0.083 | $p$ =.776 n.s. |
| | | Condition | $F_{1,25}$ = 0.228 | $p$ =.637 n.s. |
| | | Group: Condition | $F_{1,25}$ = 0.137 | $p$ =.714 n.s. |
| | (Demonstrator:Distressed) | Group | $F_{1,25}$ = 10.6 | $p$ <.01 ** |
| | | Condition | $F_{1,25}$ = 2.51 | $p$ =.126 n.s. |
| | | Group: Condition | $F_{1,25}$ = 3.22 | $p$ =.085 † |
| | Pairwise t-test (Group: Control) | Vehicle: Neutral vs Distressed | | $p$ <.001 *** |
| | | C21: Neutral vs Distressed | | $p$ <.01 ** |
| | | Neutral: Vehicle vs C21 | | $p$ =.563 n.s. |

| | | | | |
|---|---|---|---|---|
| | | Distressed: Vehicle vs C21 | | $p < .05$ * |
| | (Group: High) | Vehicle: Neutral vs Distressed | | $p = .908$ n.s. |
| | | C21: Neutral vs Distressed | | $p = .95$ n.s. |
| | | Neutral: Vehicle vs C21 | | $p = .937$ n.s. |
| | | Distressed: Vehicle vs C21 | | $p = .847$ n.s. |
| | Pairwise t-test | Neutral: Control vs High | | $p = .871$ n.s. |
| | (Condition: Vehicle) | Distressed: Control vs High | | $p < .01$ ** |
| | (Condition: C21) | Neutral: Control vs High | | $p = .72$ n.s. |
| | | Distressed: Control vs High | | $p < .05$ * |
| S7D | Two-way ANOVA | Group | $F_{1,25} = 6.240$ | $p < .05$ * |
| | | Condition | $F_{1,25} = 2.932$ | $p = .099$ † |
| | | Group: Condition | $F_{1,25} = 0.117$ | $p = .735$ n.s. |
| | Pairwise t-test (Group) | Control: Vehicle vs C21 | | $p = .072$ † |
| | | High: Vehicle vs C21 | | $p = .341$ n.s. |
| | (Condition) | Vehicle: Control vs High | | $p < .05$ * |
| | | C21: Control vs High | | $p < .05$ * |
| S7E | Two-way ANOVA | Group | $F_{1,25} = 12.118$ | $p < .01$ ** |
| | | Condition | $F_{1,25} = 2.213$ | $p = .149$ n.s. |
| | | Group: Condition | $F_{1,25} = 1.774$ | $p = .195$ n.s. |
| | Pairwise t-test (Group) | Control: Vehicle vs C21 | | $p = .855$ n.s. |
| | | High: Vehicle vs C21 | | $p = .183$ n.s. |
| | (Condition) | Vehicle: Control vs High | | $p < .01$ ** |
| | | C21: Control vs High | | $p < .01$ ** |
| S7F | Three-way ANOVA | Group | $F_{1,25} = 19.702$ | $p < .001$ *** |
| | | Demonstrator | $F_{1,25} = 16.950$ | $p < .001$ *** |
| | | Condition | $F_{1,25} = 0.655$ | $p = .426$ n.s. |
| | | Group: Demonstrator | $F_{1,25} = 13.111$ | $p < .01$ ** |
| | | Group: Condition | $F_{1,25} = 0.164$ | $p = .689$ n.s. |
| | | Demonstrator: Condition | $F_{1,25} = 0.002$ | $p = .963$ n.s. |
| | | Group: Demonstrator: Condition | $F_{1,25} = 0.962$ | $p = .336$ n.s. |
| | Two-way ANOVA | Demonstrator | $F_{1,14} = 0.981$ | $p = .339$ n.s. |
| | (Group: Control) | Condition | $F_{1,14} = 24.6$ | $p < .001$ *** |
| | | Demonstrator: Condition | $F_{1,14} = 0.402$ | $p = .536$ n.s. |
| | (Group: High) | Demonstrator | $F_{1,11} = 0.061$ | $p = .809$ n.s. |
| | | Condition | $F_{1,11} = 0.21$ | $p = .656$ n.s. |
| | | Demonstrator: Condition | $F_{1,11} = 1$ | $p = .339$ n.s. |
| | Two-way ANOVA | Group | $F_{1,25} = 15.3$ | $p < .001$ *** |
| | (Condition: Vehicle) | Demonstrator | $F_{1,25} = 8.38$ | $p < .01$ ** |
| | | Group: Demonstrator | $F_{1,25} = 9.07$ | $p < .01$ ** |
| | (Condition: C21) | Group | $F_{1,25} = 17.8$ | $p < .001$ *** |

| | | | |
|---|---|---|---|
| | Demonstrator | $F_{1,25} = 17.6$ | $p < .001$ *** |
| | Group: Demonstrator | $F_{1,25} = 9.03$ | $p < .01$ ** |
| Two-way ANOVA | Group | $F_{1,25} = 6.17$ | $p < .05$ * |
| (Demonstrator: Neutral) | Condition | $F_{1,25} = 0.828$ | $p = .372$ n.s. |
| | Group: Condition | $F_{1,25} = 0.278$ | $p = .603$ n.s. |
| (Demonstrator:Distressed) | Group | $F_{1,25} = 25.1$ | $p < .001$ *** |
| | Condition | $F_{1,25} = 0.259$ | $p = .615$ n.s. |
| | Group: Condition | $F_{1,25} = 0.564$ | $p = .46$ n.s. |
| Pairwise t-test | Vehicle: Neutral vs Distressed | | $p < .01$ ** |
| (Group: Control) | C21: Neutral vs Distressed | | $p < .001$ *** |
| | Neutral: Vehicle vs C21 | | $p = .758$ n.s. |
| | Distressed: Vehicle vs C21 | | $p = .4$ n.s. |
| (Group: High) | Vehicle: Neutral vs Distressed | | $p = .91$ n.s. |
| | C21: Neutral vs Distressed | | $p = .306$ n.s. |
| | Neutral: Vehicle vs C21 | | $p = .4$ n.s. |
| | Distressed: Vehicle vs C21 | | $p = .856$ n.s. |
| Pairwise t-test | Neutral: Control vs High | | $p < .05$ * |
| (Condition: Vehicle) | Distressed: Control vs High | | $p < .001$ *** |
| (Condition: C21) | Neutral: Control vs High | | $p < .05$ * |
| | Distressed: Control vs High | | $p < .001$ *** |

**Table S12. Detailed statistical results corresponding to Figure S7.**

| Figure | Analysis | Factor | Statistical value | p-value |
|---|---|---|---|---|
| S8A | One-way ANOVA | Condition | $F_{1,11} = 0.061$ | $p = .809$ n.s. |
| S8B | Two-way ANOVA | Condition | $F_{1,11} = 0.221$ | $p = .647$ n.s. |
| | | Demonstrator | $F_{1,11} = 12.629$ | $p < .01$ ** |
| | | Condition: Demonstrator | $F_{1,11} = 20.011$ | $p < .01$ ** |
| | Pairwise t-test | Vehicle: Neutral vs Distressed | | $p < .01$ ** |
| | (Condition) | C21: Neutral vs Distressed | | $p < .01$ ** |
| | (Demonstrator) | Neutral: Vehicle vs C21 | | $p < .05$ * |
| | | Distressed: Vehicle vs C21 | | $p < .01$ ** |
| S8C | One-way ANOVA | Condition | $F_{1,10} = 6.691$ | $p < .05$ * |
| | Pairwise t-test | Vehicle vs C21 | | $p < .05$ * |
| S8D | Two-way ANOVA | Condition | $F_{1,10} = 4.374$ | $p = .063$ † |
| | | Demonstrator | $F_{1,10} = 14.435$ | $p < .01$ ** |
| | | Condition: Demonstrator | $F_{1,10} = 4.057$ | $p = .072$ † |
| | Pairwise t-test | Vehicle: Neutral vs Distressed | | $p < .01$ ** |
| | (Condition) | C21: Neutral vs Distressed | | $p < .05$ * |
| | (Demonstrator) | Neutral: Vehicle vs C21 | | $p < .01$ ** |
| | | Distressed: Vehicle vs C21 | | $p = .459$ n.s. |
| S8E | One-way ANOVA | Condition | $F_{1,11} = 6.141$ | $p < .05$ * |
| | Pairwise t-test | Vehicle vs C21 | | $p < .05$ * |
| S8F | One-way ANOVA | Condition | $F_{1,11} = 0.046$ | $p = .834$ n.s. |
| S8G | One-way ANOVA | Condition | $F_{1,11} = 1.998$ | $p = .185$ n.s. |
| S8H | One-way ANOVA | Condition | $F_{1,10} = 10.648$ | $p < .01$ ** |
| | Pairwise t-test | Vehicle vs C21 | | $p < .01$ ** |
| S8I | One-way ANOVA | Condition | $F_{1,10} = 0.593$ | $p = .459$ n.s. |
| S8J | One-way ANOVA | Condition | $F_{1,10} = 0.273$ | $p = .613$ n.s. |

**Table S13. Detailed statistical results corresponding to Figure S8.**

| Figure | Analysis | Factor | Statistical value | p-value |
| --- | --- | --- | --- | --- |
| S9B | Two-way ANOVA | Group | $F_{2,20} = 0.207$ | $p = .815$ n.s. |
| | | Demonstrator | $F_{1,20} = 0.408$ | $p = .530$ n.s. |
| | | Group: Demonstrator | $F_{2,20} = 0.061$ | $p = .941$ n.s. |
| S9C | Three-way ANOVA | Group | $F_{2,20} = 8.353$ | $p < .01$ ** |
| | | Demonstrator | $F_{1,20} = 63.729$ | $p < .001$ *** |
| | | Condition | $F_{1,20} = 3.073$ | $p = .095$ † |
| | | Group: Demonstrator | $F_{2,20} = 10.457$ | $p < .001$ *** |
| | | Group: Condition | $F_{2,20} = 0.881$ | $p = .430$ n.s. |
| | | Demonstrator: Condition | $F_{1,20} = 0.971$ | $p = .336$ n.s. |
| | | Group: Demonstrator: Condition | $F_{2,20} = 2.036$ | $p = .157$ n.s. |
| | Two-way ANOVA | Demonstrator | $F_{1,8} = 67.6$ | $p < .001$ *** |
| | (Group: Control) | Condition | $F_{1,8} = 2.55$ | $p = .149$ n.s. |
| | | Demonstrator: Condition | $F_{1,8} = 1.68$ | $p = .231$ n.s. |
| | (Group: Low) | Demonstrator | $F_{1,5} = 11.4$ | $p < .05$ * |
| | | Condition | $F_{1,5} = 0.009$ | $p = .927$ n.s. |
| | | Demonstrator: Condition | $F_{1,5} = 0.817$ | $p = .407$ n.s. |
| | (Group: High) | Demonstrator | $F_{1,7} = 4.43$ | $p = .073$ † |
| | | Condition | $F_{1,7} = 2.66$ | $p = .147$ n.s. |
| | | Demonstrator: Condition | $F_{1,7} = 4.46$ | $p = .073$ † |
| | Two-way ANOVA | Group | $F_{2,20} = 7.05$ | $p < .01$ ** |
| | (Condition: Vehicle) | Demonstrator | $F_{1,20} = 23.3$ | $p < .001$ *** |
| | | Group: Demonstrator | $F_{2,20} = 7.34$ | $p < .01$ ** |
| | (Condition: C21) | Group | $F_{2,20} = 5.50$ | $p < .05$ * |
| | | Demonstrator | $F_{1,20} = 33.8$ | $p < .001$ *** |
| | | Group: Demonstrator | $F_{2,20} = 4.51$ | $p < .05$ * |
| | Two-way ANOVA | Group | $F_{2,20} = 2.05$ | $p = .154$ n.s. |
| | (Demonstrator: Neutral) | Condition | $F_{1,20} = 0.733$ | $p = .402$ n.s. |
| | | Group: Condition | $F_{2,20} = 0.695$ | $p = .511$ n.s. |
| | (Demonstrator:Distressed) | Group | $F_{2,20} = 11.3$ | $p < .001$ *** |
| | | Condition | $F_{1,20} = 2.28$ | $p = .147$ n.s. |
| | | Group: Condition | $F_{2,20} = 1.65$ | $p = .218$ n.s. |
| | Pairwise t-test | Vehicle: Neutral vs Distressed | | $p < .001$ *** |
| | (Group: Control) | C21: Neutral vs Distressed | | $p < .01$ ** |
| | | Neutral: Vehicle vs C21 | | $p = .456$ n.s. |
| | | Distressed: Vehicle vs C21 | | $p = .163$ n.s. |
| | (Group: Low) | Vehicle: Neutral vs Distressed | | $p = .062$ † |
| | | C21: Neutral vs Distressed | | $p < .05$ * |
| | | Neutral: Vehicle vs C21 | | $p = .328$ n.s. |
| | | Distressed: Vehicle vs C21 | | $p = .543$ n.s. |

| | | | | | |
|---|---|---|---|---|---|
| | (Group: High) | Vehicle: Neutral vs Distressed | | $p = .75$ | n.s. |
| | | C21: Neutral vs Distressed | | $p < .05$ | * |
| | | Neutral: Vehicle vs C21 | | $p = .708$ | n.s. |
| | | Distressed: Vehicle vs C21 | | $p = .056$ | † |
| | Pairwise t-test | Neutral: Control vs Low | | $p = .154$ | n.s. |
| | (Condition: Vehicle) | Neutral: Control vs High | | $p = 1$ | n.s. |
| | | Neutral: High vs Low | | $p = .497$ | n.s. |
| | | Distressed: Control vs Low | | $p = .802$ | n.s. |
| | | Distressed: Control vs High | | $p < .01$ | ** |
| | | Distressed: High vs Low | | $p < .05$ | * |
| | (Condition: C21) | Neutral: Control vs Low | | $p = .669$ | n.s. |
| | | Neutral: Control vs High | | $p = .563$ | n.s. |
| | | Neutral: High vs Low | | $p = 1$ | n.s. |
| | | Distressed: Control vs Low | | $p < .05$ | * |
| | | Distressed: Control vs High | | $p < .05$ | * |
| | | Distressed: High vs Low | | $p = 1$ | n.s. |
| S9D | Two-way ANOVA | Group | $F_{2,22} = 3.921$ | $p < .05$ | * |
| | | Condition | $F_{1,22} = 0.538$ | $p = .923$ | n.s. |
| | | Group: Condition | $F_{2,22} = 0.383$ | $p = .560$ | n.s. |
| S9E | Three-way ANOVA | Group | $F_{2,22} = 4.927$ | $p < .05$ | * |
| | | Demonstrator | $F_{1,22} = 76.813$ | $p < .001$ | *** |
| | | Condition | $F_{1,22} = 0.763$ | $p = .392$ | n.s. |
| | | Group: Demonstrator | $F_{2,22} = 4.586$ | $p < .05$ | * |
| | | Group: Condition | $F_{2,22} = 0.723$ | $p = .497$ | n.s. |
| | | Demonstrator: Condition | $F_{1,22} = 0.176$ | $p = .679$ | n.s. |
| | | Group: Demonstrator: Condition | $F_{2,22} = 1.545$ | $p = .236$ | n.s. |
| | Two-way ANOVA | Demonstrator | $F_{1,9} = 42.4$ | $p < .001$ | *** |
| | (Group: Control) | Condition | $F_{1,9} = 0.267$ | $p = .618$ | n.s. |
| | | Demonstrator: Condition | $F_{1,9} = 0.034$ | $p = .858$ | n.s. |
| | (Group: Low) | Demonstrator | $F_{1,6} = 23.7$ | $p < .01$ | ** |
| | | Condition | $F_{1,6} = 1.26$ | $p = .304$ | n.s. |
| | | Demonstrator: Condition | $F_{1,6} = 1.71$ | $p = .239$ | n.s. |
| | (Group: High) | Demonstrator | $F_{1,7} = 14.9$ | $p < .01$ | ** |
| | | Condition | $F_{1,7} = 0.239$ | $p = .64$ | n.s. |
| | | Demonstrator: Condition | $F_{1,7} = 0.955$ | $p = .361$ | n.s. |
| | Two-way ANOVA | Group | $F_{2,22} = 2.30$ | $p = .124$ | n.s. |
| | (Condition: Vehicle) | Demonstrator | $F_{1,22} = 32.0$ | $p < .001$ | *** |
| | | Group: Demonstrator | $F_{2,22} = 3.58$ | $p < .05$ | * |
| | (Condition: C21) | Group | $F_{2,22} = 3.42$ | $p = .051$ | † |
| | | Demonstrator | $F_{1,22} = 58.6$ | $p < .001$ | *** |

| | | | | |
|---|---|---|---|---|
| | | Group: Demonstrator | $F_{2,22} = 2.15$ | $p = .141$ n.s. |
| | Two-way ANOVA (Demonstrator: Neutral) | Group | $F_{2,22} = 3.17$ | $p = .062$ † |
| | | Condition | $F_{1,22} = 0.527$ | $p = .476$ n.s. |
| | | Group: Condition | $F_{2,22} = 0.093$ | $p = .912$ n.s. |
| | (Demonstrator:Distressed) | Group | $F_{2,22} = 5.31$ | $p < .05$ * |
| | | Condition | $F_{1,22} = 0.583$ | $p = .453$ n.s. |
| | | Group: Condition | $F_{2,22} = 1.24$ | $p = .309$ n.s. |
| | Pairwise t-test (Group: Control) | Vehicle: Neutral vs Distressed | | $p < .01$ ** |
| | | C21: Neutral vs Distressed | | $p < .001$ *** |
| | | Neutral: Vehicle vs C21 | | $p = .528$ n.s. |
| | | Distressed: Vehicle vs C21 | | $p = .78$ n.s. |
| | (Group: Low) | Vehicle: Neutral vs Distressed | | $p < .01$ ** |
| | | C21: Neutral vs Distressed | | $p < .05$ * |
| | | Neutral: Vehicle vs C21 | | $p = .783$ n.s. |
| | | Distressed: Vehicle vs C21 | | $p = .254$ n.s. |
| | (Group: High) | Vehicle: Neutral vs Distressed | | $p = .082$ † |
| | | C21: Neutral vs Distressed | | $p < .05$ * |
| | | Neutral: Vehicle vs C21 | | $p = .732$ n.s. |
| | | Distressed: Vehicle vs C21 | | $p = .478$ n.s. |
| | Pairwise t-test (Condition: Vehicle) | Neutral: Control vs Low | | $p = .323$ n.s. |
| | | Neutral: Control vs High | | $p = 1$ n.s. |
| | | Neutral: High vs Low | | $p = 1$ n.s. |
| | | Distressed: Control vs Low | | $p = 1$ n.s. |
| | | Distressed: Control vs High | | $p = .085$ † |
| | | Distressed: High vs Low | | $p = .173$ n.s. |
| | (Condition: C21) | Neutral: Control vs Low | | $p = .131$ n.s. |
| | | Neutral: Control vs High | | $p = 1$ n.s. |
| | | Neutral: High vs Low | | $p = .432$ n.s. |
| | | Distressed: Control vs Low | | $p = .161$ n.s. |
| | | Distressed: Control vs High | | $p = .122$ n.s. |
| | | Distressed: High vs Low | | $p = 1$ n.s. |
| S9F | Two-way ANOVA | Group | $F_{2,20} = 11.687$ | $p < .001$ *** |
| | | Condition | $F_{1,20} = 0.441$ | $p = .514$ n.s. |
| | | Group: Condition | $F_{2,20} = 0.068$ | $p = .934$ n.s. |
| | Pairwise t-test (Group) | Control: Vehicle vs C21 | | $p = .723$ n.s. |
| | | Low: Vehicle vs C21 | | $p = .864$ n.s. |
| | | High: Vehicle vs C21 | | $p = .383$ n.s. |
| | (Condition) | Vehicle: Control vs Low | | $p < .05$ * |
| | | Vehicle: Control vs High | | $p < .05$ * |
| | | Vehicle: Low vs High | | $p = 1$ n.s. |

| | | | | | |
|---|---|---|---|---|---|
| | | C21: Control vs Low | | $p < .01$ | ** |
| | | C21: Control vs High | | $p < .05$ | * |
| | | C21: Low vs High | | $p = 1$ | n.s. |
| S9G | Two-way ANOVA | Group | $F_{2,20} = 0.616$ | $p = .550$ | n.s. |
| | | Condition | $F_{1,20} = 6.280$ | $p < .05$ | * |
| | | Group: Condition | $F_{2,20} = 0.159$ | $p = .159$ | n.s. |
| | Pairwise t-test | Control: Vehicle vs C21 | | $p = 1$ | n.s. |
| | (Group) | Low: Vehicle vs C21 | | $p = 1$ | n.s. |
| | | High: Vehicle vs C21 | | $p < .01$ | ** |
| | (Condition) | Vehicle: Control vs Low | | $p = 1$ | n.s. |
| | | Vehicle: Control vs High | | $p = .887$ | n.s. |
| | | Vehicle: Low vs High | | $p = 1$ | n.s. |
| | | C21: Control vs Low | | $p = 1$ | n.s. |
| | | C21: Control vs High | | $p = 1$ | n.s. |
| | | C21: Low vs High | | $p = .421$ | n.s. |
| S9H | Two-way ANOVA | Group | $F_{2,20} = 1.353$ | $p = .281$ | n.s. |
| | | Condition | $F_{1,20} = 2.679$ | $p = .117$ | n.s. |
| | | Group: Condition | $F_{2,20} = 1.764$ | $p = .197$ | n.s. |
| S9I | Two-way ANOVA | Group | $F_{2,20} = 6.882$ | $p < .01$ | ** |
| | | Condition | $F_{1,20} = 17.537$ | $p < .001$ | *** |
| | | Group: Condition | $F_{2,20} = 4.379$ | $p < .05$ | * |
| | Pairwise t-test | Control: Vehicle vs C21 | | $p = .063$ | † |
| | (Group) | Low: Vehicle vs C21 | | $p = .948$ | n.s. |
| | | High: Vehicle vs C21 | | $p < .001$ | *** |
| | (Condition) | Vehicle: Control vs Low | | $p = .157$ | n.s. |
| | | Vehicle: Control vs High | | $p < .05$ | * |
| | | Vehicle: Low vs High | | $p = 1$ | n.s. |
| | | C21: Control vs Low | | $p < .01$ | ** |
| | | C21: Control vs High | | $p = .201$ | n.s. |
| | | C21: Low vs High | | $p = .135$ | n.s. |
| S9J | Two-way ANOVA | Group | $F_{2,20} = 2.277$ | $p = .129$ | n.s. |
| | | Condition | $F_{1,20} = 20.171$ | $p < .001$ | *** |
| | | Group: Condition | $F_{2,20} = 4.531$ | $p < .05$ | * |
| | Pairwise t-test | Control: Vehicle vs C21 | | $p < .01$ | ** |
| | (Group) | Low: Vehicle vs C21 | | $p = .711$ | n.s. |
| | | High: Vehicle vs C21 | | $p < .01$ | ** |
| | (Condition) | Vehicle: Control vs Low | | $p = 1$ | n.s. |
| | | Vehicle: Control vs High | | $p = 1$ | n.s. |
| | | Vehicle: Low vs High | | $p = 1$ | n.s. |
| | | C21: Control vs Low | | $p = .213$ | n.s. |

| | | | | |
|---|---|---|---|---|
| | | C21: Control vs High | | $p$ =.473 n.s. |
| | | C21: Low vs High | | $p$ <.05 * |
| S9K | Two-way ANOVA | Group | $F_{2,20}$ = 0.849 | $p$ =.443 n.s. |
| | | Condition | $F_{1,20}$ = 3.283 | $p$ =.085 † |
| | | Group: Condition | $F_{2,20}$ = 2.010 | $p$ =.160 n.s. |

**Table S14. Detailed statistical results corresponding to Figure S9.**
